# CLICK-enabled analogues reveal pregnenolone interactomes in cancer and immune cells

**DOI:** 10.1101/800649

**Authors:** Sougata Roy, James Sipthorp, Bidesh Mahata, Jhuma Pramanik, Marco L. Hennrich, Anne-Claude Gavin, Steven V. Ley, Sarah A. Teichmann

## Abstract

Pregnenolone (P5) promotes prostate cancer cell growth, and *de novo* synthesis of intratumoural P5 is a potential cause of development of castration-resistance. Immune cells can also synthesize P5 *de novo*. Despite its biological importance, little is known about P5’s mode of actions, which appears to be context-dependent and pleiotropic. A comprehensive proteome-wide spectrum of P5-binding proteins that are involved in its trafficking and functionality remains unknown. Here, we describe an approach that integrates chemical biology for probe synthesis with chemoproteomics to map P5-protein interactions in live prostate cancer cells and murine CD8^+^ T cells. We subsequently identified P5-binding proteins potentially involved in P5-trafficking, and in P5’s non-genomic action that may drive the promotion of castrate-resistance prostate cancer and regulate CD8^+^ T cell function. We envisage that this methodology could be employed for other steroids to map their interactomes directly in a broad range of living cells, tissues and organisms.

## Introduction

Pregnenolone (P5) is the first bioactive steroid hormone and precursor of all other steroid hormones in steroid biosynthesis (steroidogenesis) pathways. P5 is synthesized from cholesterol by the enzyme CYP11A1 inside the mitochondria of steroidogenic cells^1^. A high capacity for P5 biosynthesis has been reported in adrenal tissue, gonads and placenta. Extra-adrenal and extra-gonadal P5 synthesis (also known as local steroidogenesis) has been reported in lymphocytes^2–4^, adipocytes^5^, the nervous system^6^, tumours^7^ and tumour-infiltrating immune cells^8^. The role played by this local P5 synthesis is poorly understood^9^, particularly in pathologies such as cancer. In prostate cancer and melanoma P5 promotes tumour growth^8, 10^, while in glioma it restricts tumour growth^11^. The mode-of-action of P5 in tumours is incompletely understood.

In the nervous system, P5 is known to regulate synapse formation, outgrowth of neurites and enhances myelinization^12^, improves cognitive and memory function^13^. During immune response against helminth parasite infection T helper cells synthesize P5 to restore immune homeostasis^4^. Up to now, we do not have a proteome-wide description of the P5-interacting molecules in any living cells.

Traditionally, steroid hormones have been considered to act by regulating transcription^14^. However, rapid activity of steroid hormones can be mediated by non-genomic pathways^15^. Non-genomic pathways appear to mediate P5 activity^16^ in a cell type-specific and context-dependent manner, indicating a need for proteome-wide studies to map the full spectrum of P5 functions.

Here, we have developed a chemical biology method to generate clickable P5-analogues for use in living cells. Exploiting these P5-probes in combination with quantitative mass-spectrometry, we profiled global P5-protein interactions directly in two distinct cell types: a steroid-sensitive cell line derived from a metastatic prostate cancer patient (i.e. LNCaP) and *de novo* P5-producing mouse CD8^+^ T cells.

Altogether, we identified 62 high-confidence P5 binding and about 387 potential P5-binding proteins localized in the nucleus, mitochondria and endoplasmic reticulum. These proteins include receptors, channels, transporters, cytoskeletal proteins such as vimentin and enzymes, of which many represent novel interactions. Overall, we identified P5-binding proteins potentially involved in inter- and intra-cellular P5-trafficking and P5’s non-genomic action that drives prostate cancer promotion of castration-resistance, and mediates CD8^+^ T cell regulation.

This study unravels prospective routes for understanding pregnenolone biochemistry in different cellular contexts such as prostate cancer progression and immune cell regulation. We demonstrate a general methodology to decipher P5-biochemistry in *de novo* steroidogenic and steroid-responsive cells or tissues.

## Results

### Design of clickable and photoactivatable P5 probes

To capture the P5 interactome in living cells, we utilise photoaffinity labelling combined with enrichment of tagged proteins using a bioorthogonal handle. This strategy required modification of the P5 core, whilst also retaining the primary pharmacology of P5. A minimalist photoaffinity enrichment linker has been introduced to kinase inhibitors to map their interactomes^17^. The linker utilises a diazirine as the UV-induced cross-linking agent and an alkyne tag, upon which bioorthogonal chemistry could be performed after photoaffinity labelling to enrich tagged proteins using azide-biotin (Figure 1a). The linker was selected due to its small size, reducing the risk of the tag altering the P5 interactome, and due to the proven effectiveness of this type of approach^17–19^. With these considerations in mind, we synthesized molecule **8** (Figure 1b) to conjugate to P5 at three different positions to ensure maximum coverage of molecular space, and avoid preclusion of binding by a particular linker. The P5-derived probes were named **P5-A**, **P5-B1**, **P5-B2** and **P5-C** (Figure 1c and S1), and tested for bioactivity and cell permeability.

**Figure 1.**
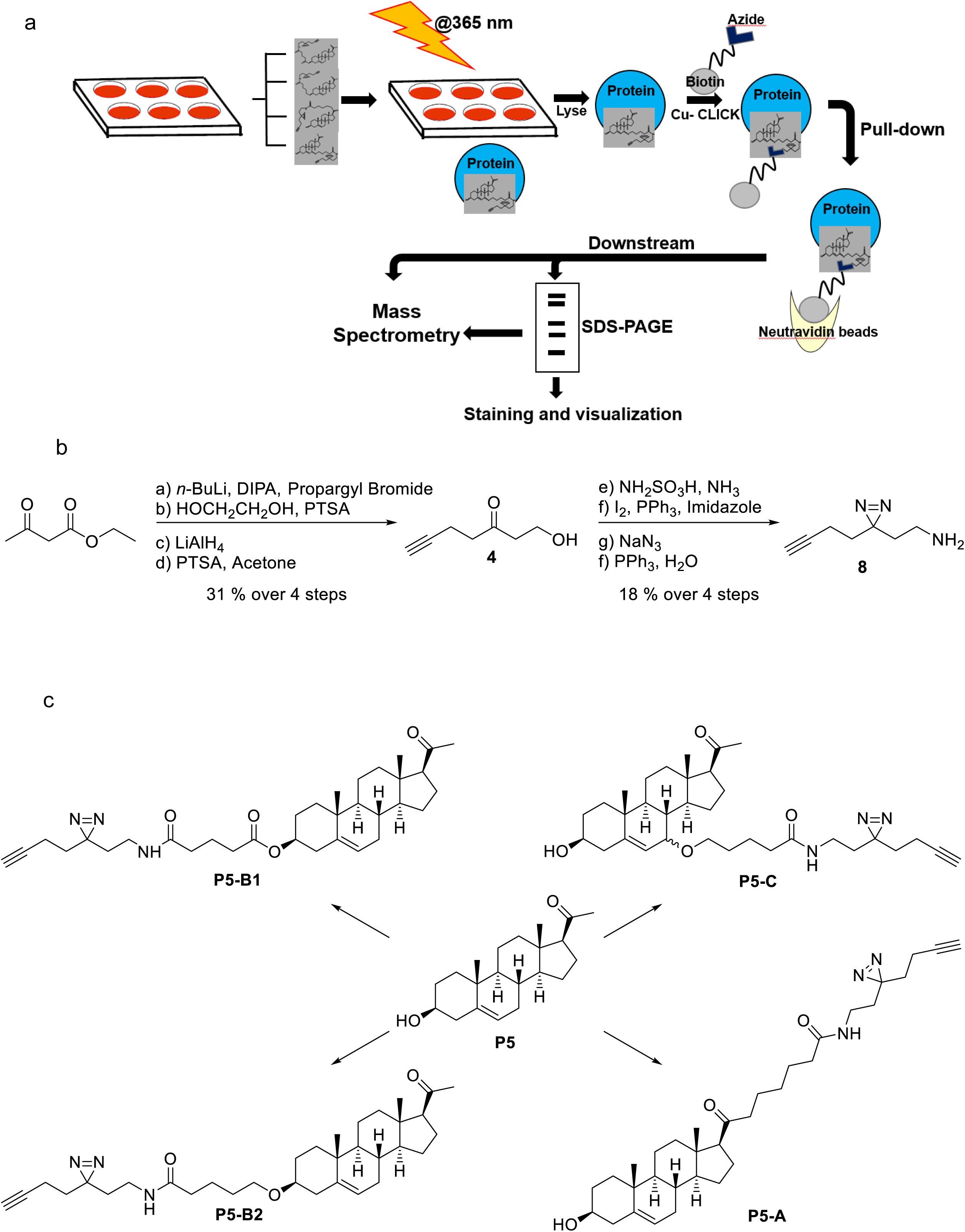
Design of clickable and photoactivable pregnenolone (P5) probes. a. Schematic diagram of our chemoproteomic approach to pull-down the P5-interacting proteome in live cells. b. Synthesis of the photoactivatable and clickable linker with diazirine and alkyne. c. Final chemical structures of the four different probes with their linker positions as compared to the mother molecule P5.

### P5 probes mimic biological activity of P5 in human cancer cell lines

P5 is known to stimulate LNCaP prostate cancer cell growth^10^ and inhibit U87MG glioma cell growth^11^. To test the biological activity of the **P5-A**, **P5-B1**, **P5-B2** and **P5-C** we performed cell viability assays using the LNCaP and U87MG cells. While LNCaP cells show increased cell viability in the presence of P5, it has a cytotoxic effect on U87MG cells (Figure 2a). For LNCaP cells, only 2 nM of **P5-A** is sufficient to induce a four-fold increase in cell proliferation, which corresponds to the effect of 20 nM of P5 (Figure S2 a). However, at higher concentration of **P5-A**, there is decrease in cell viability. For **P5-C**, cell viability follows exactly the same pattern as for P5. In both cases, cell proliferation gradually increases from 3 fold to 4 fold between 2 nM and 20 nM concentration of **P5-C** or P5 (Figure 2a top panel). **P5-B1** and **P5-B2** (although slightly better than **P5-B1**) can only induce cell growth to 3 fold corresponding to the highest concentration i.e. 20 nM (Figure S2a). In summary, **P5-C** shows the best retention of the parental pharmacology of P5.

**Figure 2.**
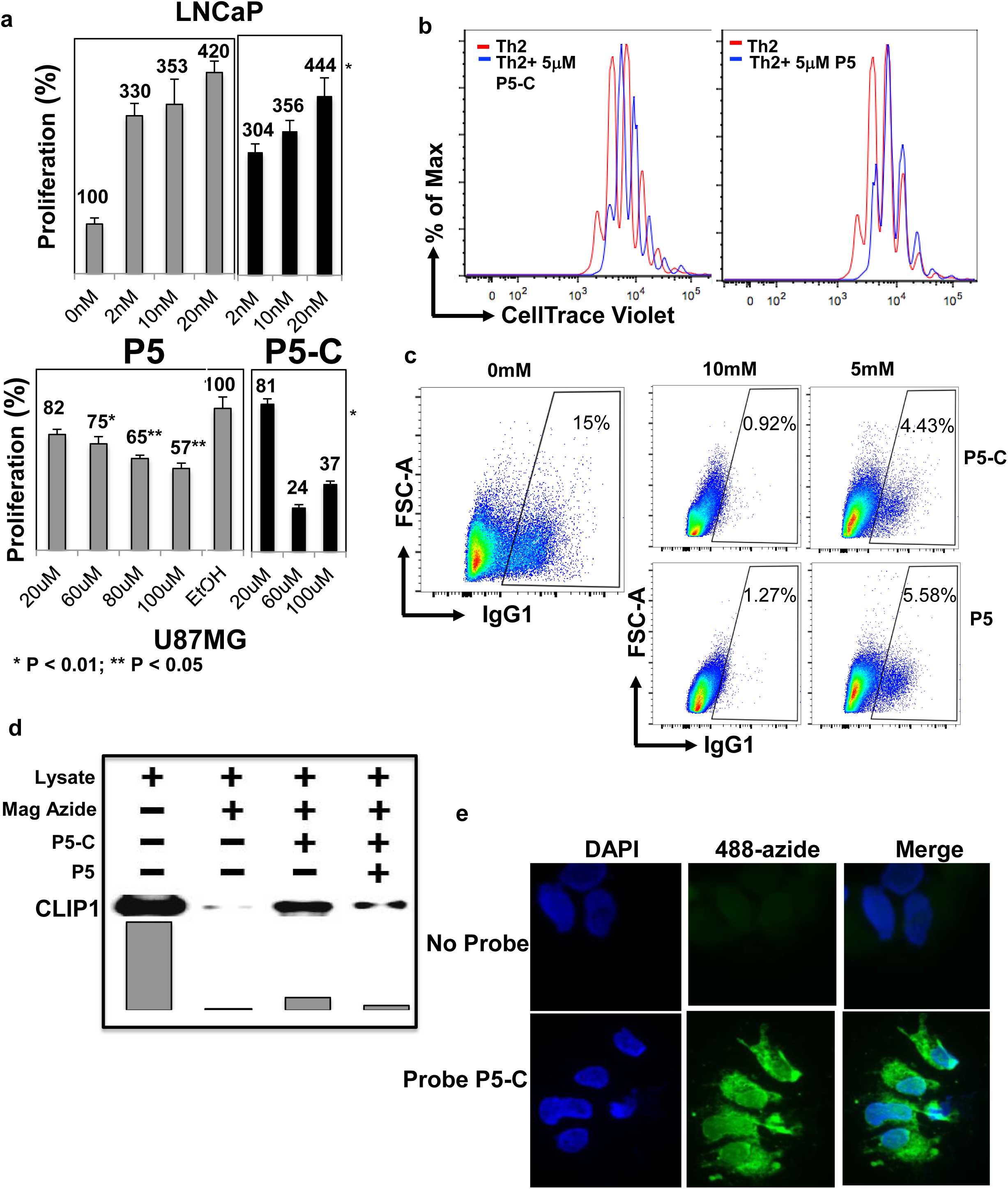
P5-C mimics biological activity of P5. **a. P5-C** mimics biological activity of P5 in human prostate cancer and glioma cell lines. The LNCaP (top panel) and U87MG (bottom panel) cells were incubated equivalent amounts of P5-C (right histograms) and P5 (left histogram). Cell proliferation was evaluated using the XTT cell viability assay in at least 5 independent experiments. **b. P5-C** mimics biological activity of P5 in mouse primary T cells and inhibits T cell proliferation. Naive CD4^+^ T cells were stained with CellTrace Violet and activated under Th2 differentiation conditions for 72 hrs in the presence (red histogram) or absence (blue histogram) of 5 µM P5 or P5-C. The cell proliferation profile was captured by a flow cytometry-based dye decay assay. Data shown are representative of three independent experiments with three mice in each experiment. **c. P5-C** mimics biological activity of P5 in mouse primary B cells and inhibits B cell immunoglobulin class switching. Naive resting B cells were induced with LPS and IL4 in the presence of different concentrations of P5 and P5-C (0, 5 and 10 µM). Cell-surface expression of IgG1 was analyzed by flow cytometry on the fifth day of stimulation. Data shown are representative of three independent experiments with three mice in each experiment. **d.** The **P5-C** probe mimics P5 interaction with the previously known P5-binding protein CLIP1, and is competed out by cold P5. HA-tagged CLIP1 was ectopically expressed in HEK cells and the protein content of the lysate was estimated. About 400 µg of whole cell lysate was used in 2 experiments simultaneously, one incubated with 50 nM **P5-C** and another with 500 nM P5 before 50 nM P5-C. The azide-coated magnetic beads were clicked and pulled down using magnets. The magnetic-azide beads were incubated with cell lysate to capture any background binding of CLIP1 to the beads. SDS-PAGE and Western blotting followed by incubation with HA antibody revealed P5-C binding to CLIP 1, and P5 competed out P5-C binding. **e. P5-C** is cell permeable. Live LNCaP cells are incubated with P5-C, which is clicked to alexa fluor 488 azide (in green) and imaged under a fluorescence microscope. Simultaneously, as a control, cells not incubated with P5-C were clicked and imaged as before. DAPI (in blue) was used in both the experiments to visualize the nucleus.

U87MG cells are known to undergo apoptosis in the presence of P5. All four probes induced apoptosis at lower concentrations than the parental P5. **P5-B2** showed the highest apoptotic activity with 3-fold reduction in cell viability at only 20 μM concentration; the same concentration of P5 does not induce any significant reduction of cells. In contrast, 100 μM of all the probes produces a significant reduction in cell number that also corresponds to the cell viability at 100 μM the cells with P5 (Figure 2a bottom panel and Figure S2 b).

### P5 probes mimic the biological activity of P5 in primary immune cells

P5 inhibits murine T helper cell proliferation and B cell class switching^4^. To test whether addition of the linker retains the biological activity of P5 in this context, we performed T helper cell proliferation assays and B cell immunoglobulin class switching assays in the presence of **P5-A, P5-B1, P5-B2** and **P5-C**. The presence of P5 in the *in vitro* Th2 culture conditions significantly restricts cell proliferation compared to vehicle-only-treated conditions (Figure 2b, right-side panel). We found that **P5-C**, like P5, inhibits T cell proliferation (Figure 2b, left-side panel). **P5-A**, **P5-B1** and **P5-B2** showed similar properties (Figure S3 a). In immunoglobulin class-switching experiments we observed that the **P5-C** and the other P5-probes are equally efficient as P5 (Figure 2c and S3 b).

### P5 probes mimic P5 protein interactions and is cell permeable

CAP-Gly domain containing linker protein 1 or cytoplasmic linker protein 1 (CLIP1, also known as CLIP170) was previously reported to be a specific P5 binding protein in zebrafish and human^19^. To test whether our P5 probes mimic P5’s binding activity with CLIP1, we expressed CLIP1 ectopically in HEK293 cells, and the lysate was used to analyse the CLIP1 binding affinity of the P5-probes. All the P5 probes were able to pull down the CLIP1 protein from the lysate with similar efficiency (Figure 2d and S4). Unlabelled P5 was able to compete out **P5-C** binding to the CLIP1 protein, indicating **P5-C**’s ability to capture P5 interactome (Figure 2d), and clearly reflecting its ability to bind known P5-binding proteins.

To confirm the cell permeability of **P5-C**, we cultured LNCaP cells in the presence or absence of **P5-C** followed by CLICK reaction with a fluorophore bearing azide. Subsequent fluorescence microscopy showed that the **P5-C** could enter live LNCaP cells (Figure 2e).

### P5-C enriches P5-binding proteins from cell extracts

To ascertain that **P5-C** can mimic P5’s binding ability, LNCaP cell extracts were incubated with **P5-C** in the presence or absence of competing amounts of P5. In parallel, two other controls were performed, one without UV irradiation and another without **P5-C** being present. All the samples were loaded onto an SDS-PAGE gel and silver stained (Figure 3a). 10-fold excess of P5 compared to **P5-C** was able to effectively compete out the **P5-C** binding, confirming our earlier observation (with CLIP1). **P5-C** captures P5 interacting proteins in a complex protein mixture. The samples without UV treatment and the neutravidin beads did not show any significant binding. The above benchmarking and quality control results confirmed the **P5-C** molecule as true analogue of P5, and an ideal probe to study *in vivo* P5 interactomes. LNCaP cells are steroid sensitive prostate cancer cell line, which has been used widely as a model of human prostate cancer and was previously shown to respond to P5^10^. Prostate cancer cells and tumour-infiltrating T cells produce P5 *de novo*, which can cause autocrine and paracrine responses in the tumour microenvironment^7, 8^. Therefore, we proceeded to use these two cell types to reveal the proteome-wide map of P5-interacting proteins.

**Figure 3.**
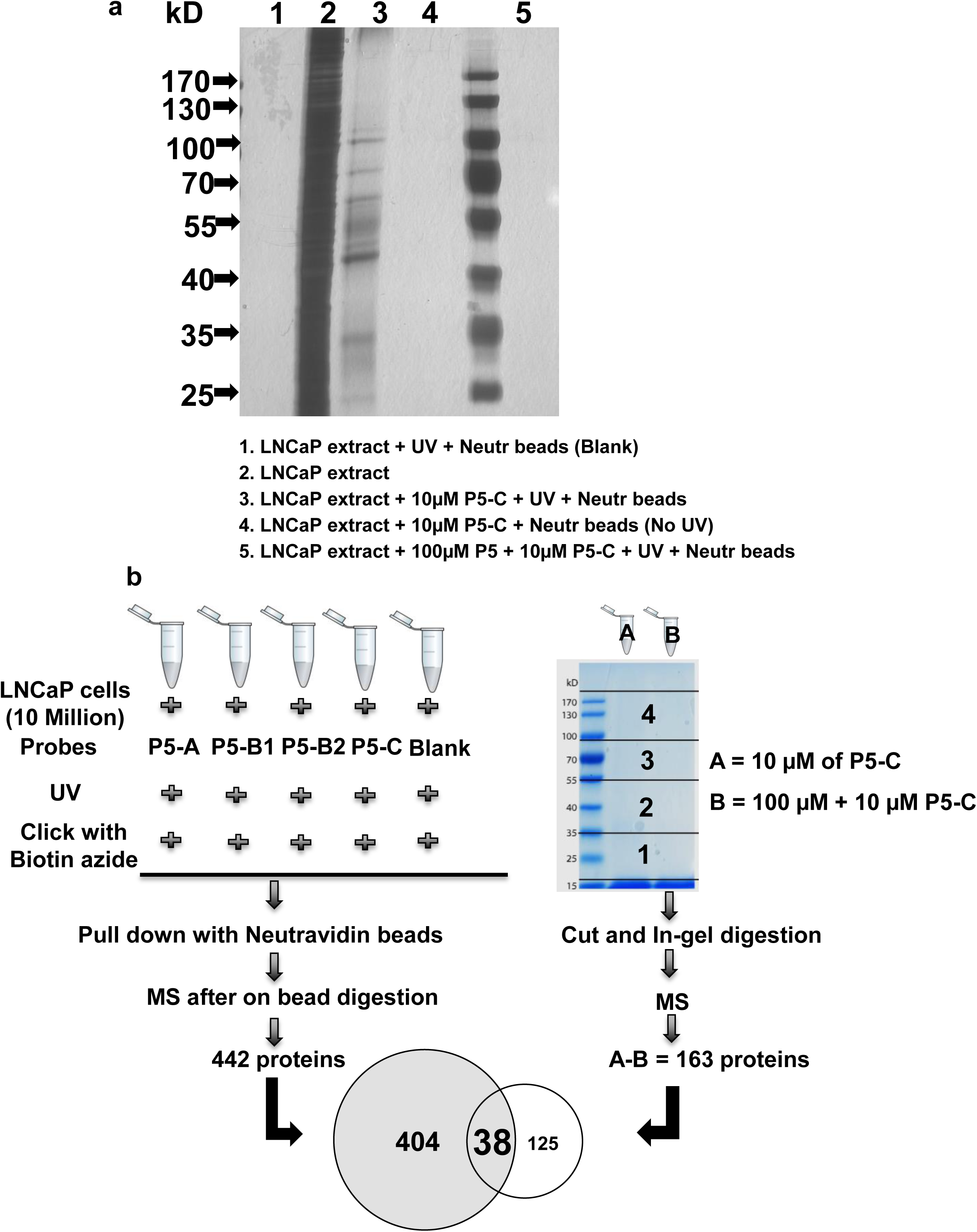
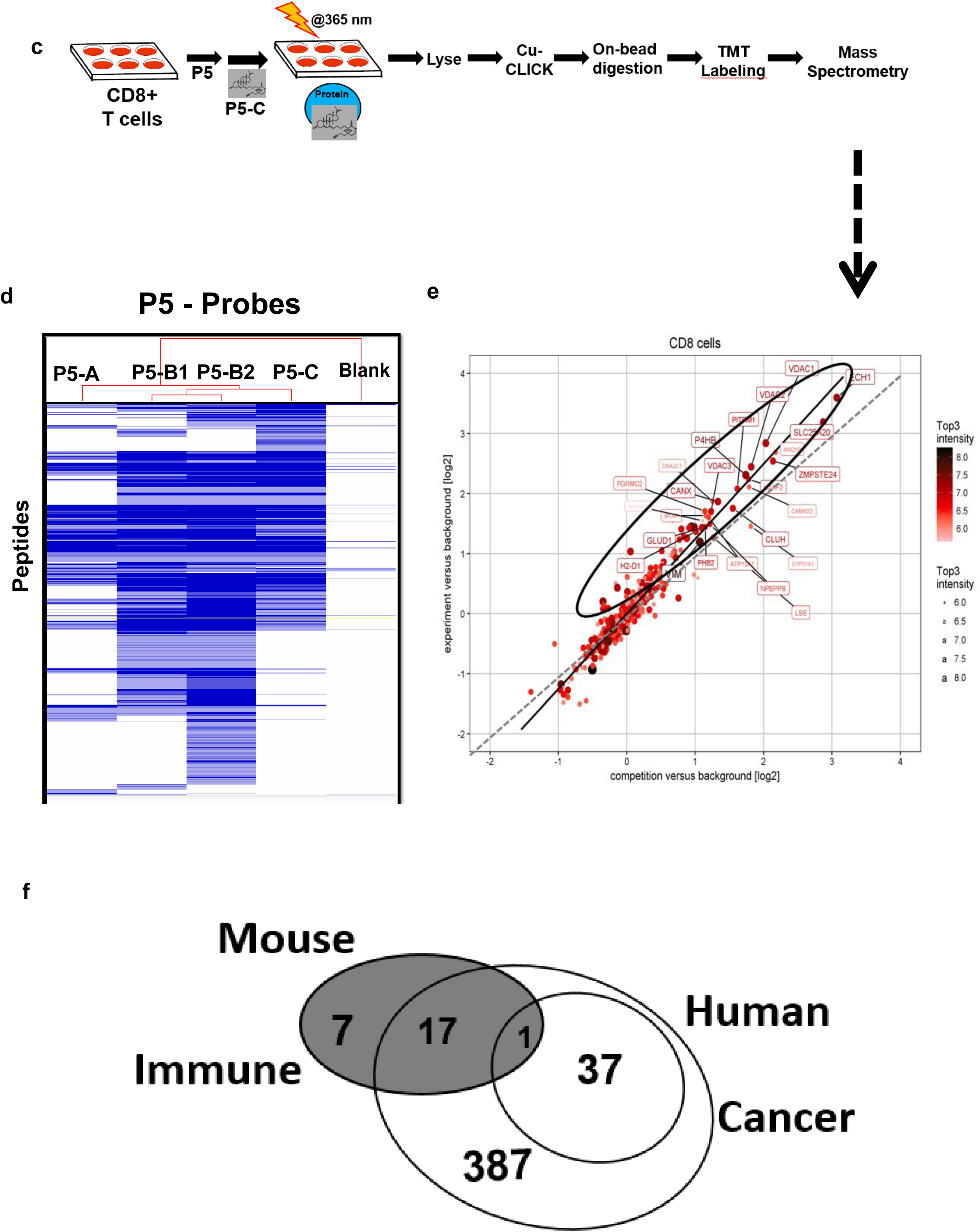
P5 probes enable mass-spectrometry profiling of P5-binding proteins in live cells. a. Gel profiling and specificity of P5-binding proteins with **P5-C** in LNCaP cell extracts. 400 µg of LNCaP protein *in vitro* was mixed either with P5-C alone or in the presence of 10X P5 (competition assay), exposed to UV and loaded onto lanes 3 and 5 respectively in a 10% SDS-PAGE. Two control experiments were run in parallel, one without P5-C to check background binding to the beads (lane 1). Another without UV to check the proper activation of the diazirine within the linker (lane 4). Lane 2 has 50 µg of total LNCaP proteins. The gel was stained with silver and imaged. b. Experimental design to reveal the identity of P5-interacting proteins in LNCaP cells using two approaches. c. Experimental design to reveal the identity of P5-interacting proteins in live CD8^+^ T cells. d. Hierarchical clustering of all proteins pulled down using all four probes shows the structure-dependent bioactivity of the probes. e. 23 P5 interacting proteins in CD8^+^ T cells. The proteins extracted with P5-C either alone or after competition with P5. P5-C binding proteins either in presence of P5 (competition assay) or its absence (experiment) were plotted on a log scale. The regression equation fitting the two variables is represented by the solid straight line, while the dotted line represent the X=Y linear relation showing no variation. The dots in ellipses represent the P5C-binding proteins whose binding can be competed out in the presence of P5. f. Common and specific P5-interating proteins in human prostate cancer and mouse immune cells. P5-interactomes from human prostate cancer (two concentric empty circles) and murine CD8^+^ T cells (filled circle) have 16 proteins in common. Among them, only 1 protein was common to CD8^+^ T cells and core ‘P5 binding proteins’ from LNCaP. Another 17 CD8^+^ T cell proteins were found in the 404 ‘potential P5 binding proteins’ from LNCaP. 7 and 424 (37 + 387) proteins are specific to CD8^+^ T cells and LNCaP respectively.

### P5 probes enable mass-spectrometry profiling of P5-binding proteins in LNCaP and murine CD8^+^ T cells

P5-interactome in LNCaP and CD8+ T cells were captured as per the schematics (Figure 3b and c). Hierarchical clustering showed the binding potential of the probes to be distinct, which might be associated with the availability of molecular space due to the distinct pattern of linker substitution (Figure 3d). **P5-B1** and **P5-B2** showed the closest correlation, which was expected as they were both substituted at the 3 – position of P5. The slight differences might be due to the addition of a carbonyl group in **P5-B2**. The protein pull-down efficiency of **P5-A** was lowest as compared to the others.

To generate consensus P5 interactome, we considered the four different probes as four replicates and only accepted those proteins with >2 unique peptides in at least 3 replicates as ‘potential P5 binding’ (described in detail in the Methods section under the heading ‘Analysis of the MS results’). This allowed us to obtain a list of 442 LNCaP proteins.

In a parallel approach, we used **P5-C** with or without competition with P5. After SDS-PAGE gel analysis, the gel lanes were cut out, and subsequently digested in-gel followed by LC-MS analysis to identify proteins (Figure 3b and S5). We then selected only those proteins that could be effectively competed by P5, thereby restricting to proteins with high specificity for P5. We captured 163 P5-interacting proteins. Among those, 38 were common to both approaches described above, and therefore termed as ‘P5 binding proteins’ (Table 1). The rest of the 403 out of the 441 proteins obtained from the on-bead digestion experiment were simultaneously analysed and named as ‘potential P5 binding proteins’ (Figure 3b). (125 out of the 163 proteins from the in-gel digestion experiment were excluded from further analysis.)

**Table1:**
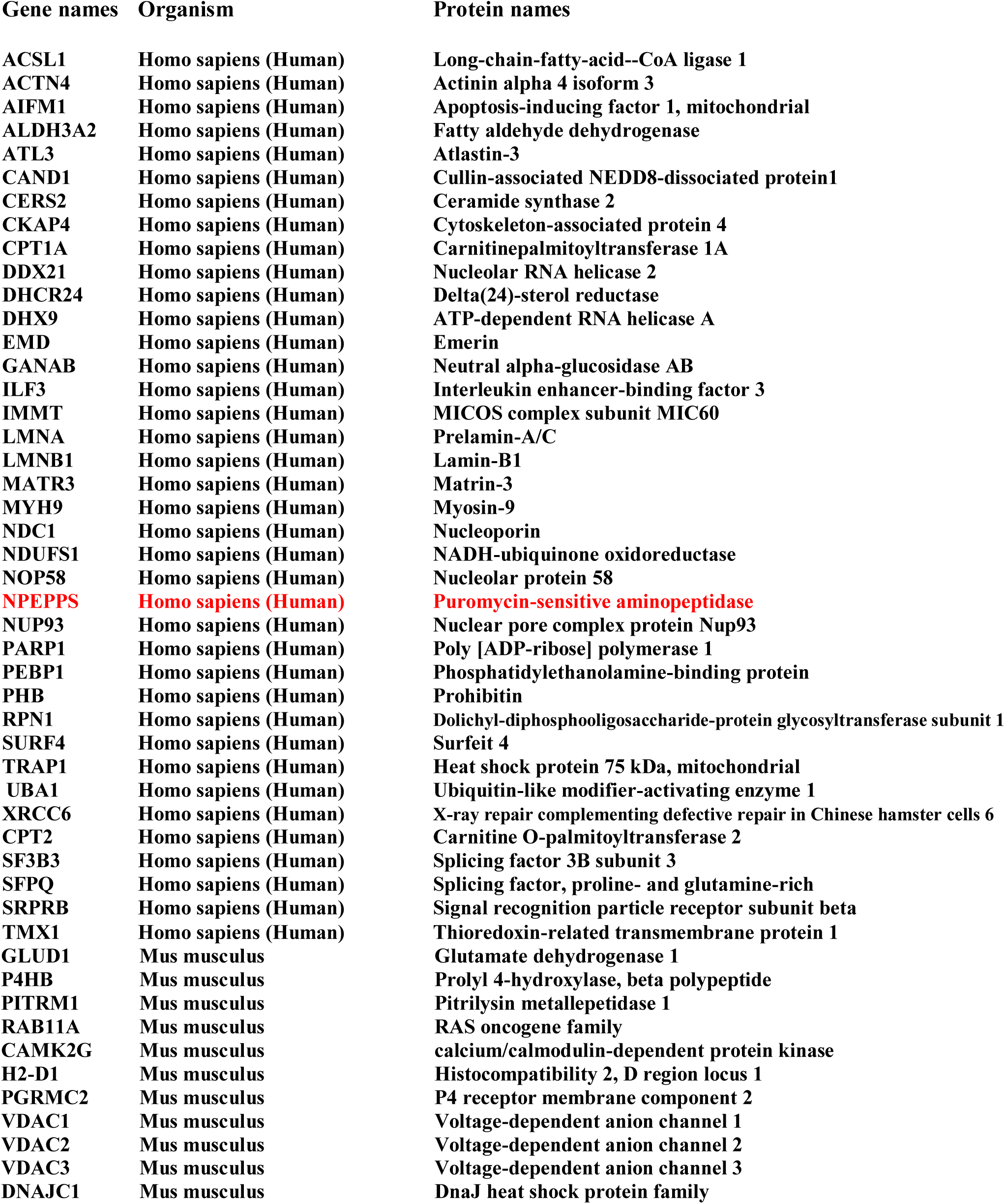
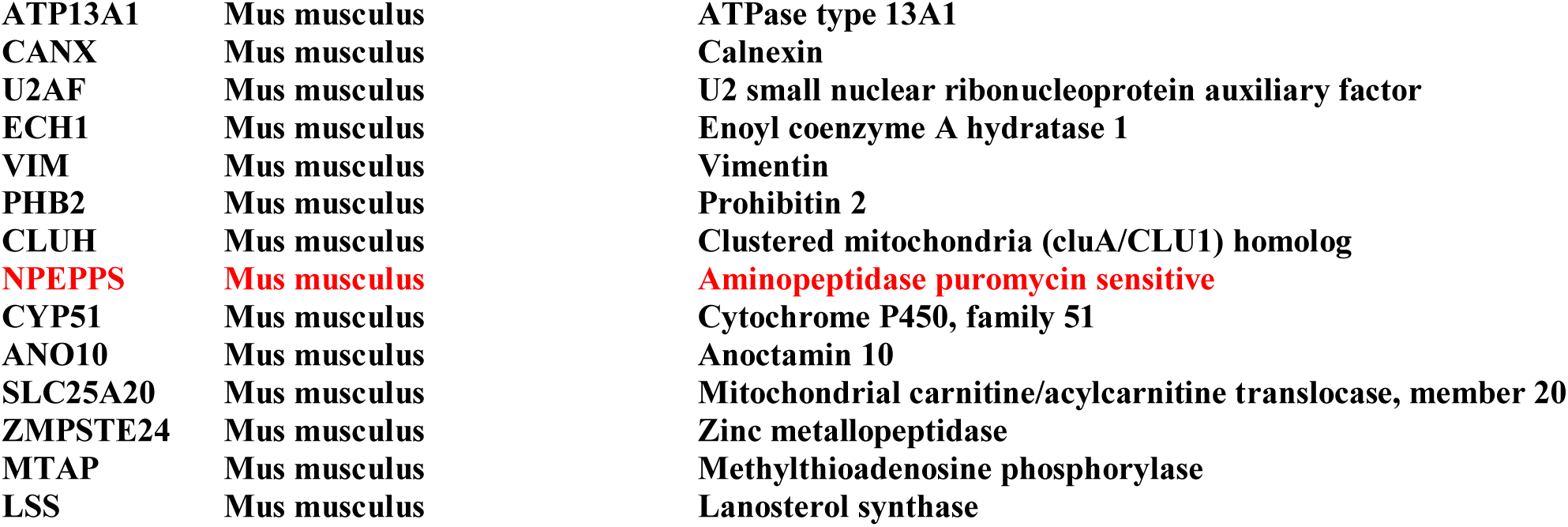
The details of 62 ‘P5 binding proteins’.

To identify proteins enriched with **P5-C** in murine immune cells, we carried out proteomics using the tandem mass tag (TMT) for relative quantification. The schematic in Figure 3c outlines the protocol. Analysis revealed 25 proteins in the P5-interactome in CD8^+^ T cells (Figure 3e). Overall, 18 of these 25 murine proteins were also found in human LNCaP. The remaining seven are CD8^+^ T cell-specific proteins (Figure 3f).

In total, we enriched a catalogue of 62 high confidence ‘P5 binding proteins’ and about 387 ‘potential P5 binding’ proteins from two different cell types.

### P5-Interacting proteins in cancer and immune cells are predominantly localized in organellar membranes

Quantitative mass spectrometry experiments successfully allowed us to profile and categorise P5-interacting proteins. Overall in the two cell types, 49 out of 62 (80%) are membrane proteins, with 27 of the 49 membrane proteins (∼55%) being transmembrane (Figure 4a left panel). In total nuclear, mitochondrial and ER-localized proteins represent 87% of the P5-binding proteins (Figure 4a right panel). Overall, 54 organellar proteins are almost equally distributed among nucleus, mitochondria and ER (Figure 4b). Interestingly, a majority of the P5 enriched proteins in prostate cancer are annotated as nuclear (65%), whereas ER and mitochondrial (68%) proteins form the majority in CD8^+^ T cells (Figure 4d and e).

**Figure 4.**
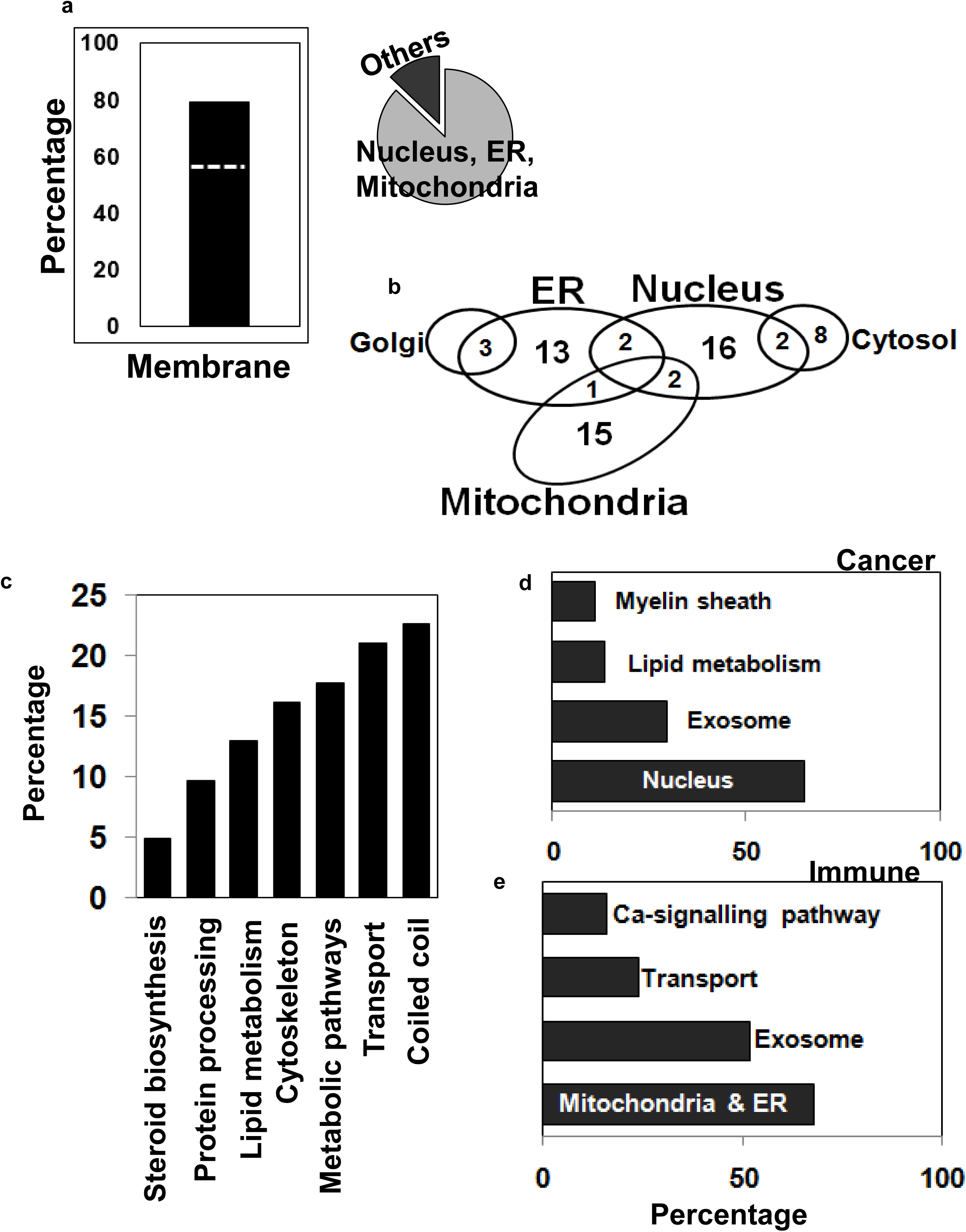
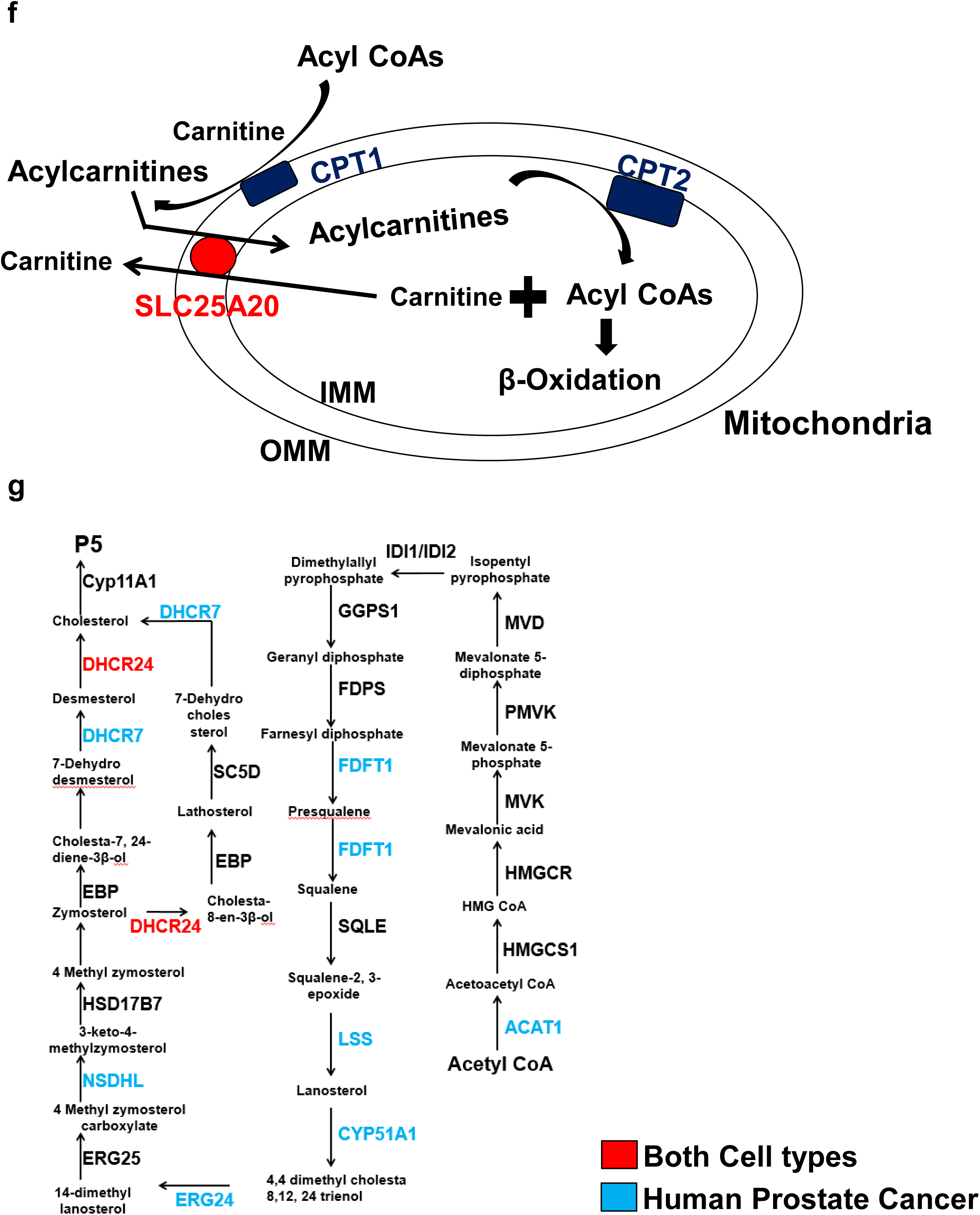
General and distinct P-5 interacting protein annotation in cancer and immune cells. a. (Left panel) P5-binding protein activity is mediated via membrane proteins, and the majority (55%) of membrane proteins in P5-interactome are transmembrane (dotted line). (Right panel) The P5-interactome acts through the nucleus, mitochondria and ER. b. Compartmentalization of the 54 organellar proteins enriched from the two cell types. c. Annotation of the 62 protein high confidence P5-interactome. d. Functional classification of P5-binding proteins from prostate cancer cells. e. Functional categorization of P5-binding proteins from CD8+ T cells. f. The main carnitine shuttle proteins that regulate β-oxidation are P5 targets. Colour codes are cell type-specific; (red) for both cell types and (green) for only cancer cells. g. Many cholesterol biosynthesis enzymes are P5 targets. The enzymes from each step of the pathway are colour coded (same as above) showing common and distinct cell type-specific P5 targets.

Consistent with a non-genomic mode of action of P5, none of the nuclear-localized proteins are annotated as putative steroid receptors, but we did retrieve the Progesterone receptor membrane component 2 (PGRMC2) in the P5 interactome from both cell types. The PGRMC2 family belongs to the unconventional membrane receptor family of the hormone progesterone (P4)^20^, a direct product of pregnenolone. *De novo* synthesis of P5 in CD8^+^ T cells indicated P5’s role in modulating immune cell plasticity and differentiation^21^. 17 out of 25 (70%) of the P5-interacting proteins in CD8+ T cells are localized in membranes, which were annotated to be either mitochondrial or ER.

### P5’s activity is mediated by metabolic pathways

In common across both cell types, about 18% of the P5 binding proteins represent metabolic enzymes, and 13% belong to lipid metabolism (Figure 4c). Three proteins (5%) are key enzymes that regulate lanosterol and cholesterol biosynthesis. Specific to the prostate cancer cells, 13% of the ‘P5 binding proteins’ belong to lipid metabolism pathways (Figure 4d), that are mostly absent in CD8^+^ T cells. The list contains key regulators of the fatty acid metabolism pathway, such as the carnitine palmitoyltransferase family proteins CPT1A and CPT2. These enzymes are from the outer and inner mitochondrial membrane respectively, and control the β-oxidation pathway in mitochondria (Figure 4f).

In addition, P5 binds to carnitine acyl carnitine translocase (SLC25A20), the carnitine and acylcarnitine transporter in both LNCaP and CD8^+^ T cells (Figure 4f). Acyl CoA synthetase ACSL1, which regulates the metabolism of fatty acids through its conversion to respective acyl CoAs, also binds to P5. P5-binding proteins were also found to be involved in ceramide and N-glycan biosynthesis pathways. These metabolic pathways are known to play key role in prostate cancer progression^22–24^. Many enzymes in the cholesterol biosynthesis pathway are found in LNCaP cells or both cell types (Figure 4g). A similar binding pattern has been noted for cholesterol^18^. Considering all 442 P5-binding proteins in LNCaP cells, we also note a significant over-representation of other metabolic pathways (Figure S6).

### Click-proteomics reveals P5 interactions with cellular transport and cytoskeletal proteins

The classical model of diffusion-mediated steroid transport is not sufficient to explain the steroid trafficking within and across cells, and therefore specific receptor-mediated transport and signalling mechanisms are gaining in importance^25, 26^. To date, the data regarding P5 inter- and intra-cellular transport is very limited. About a fifth (13 out of 62) of the P5 interactome proteins are related to transport (Figure 4c). The voltage-dependent ion channels, VDAC1, 2 and 3 are the primary regulators of metabolite exchange between the mitochondria and the rest of the cell. We retrieved VDAC1, 2 and 3 from both cell types.

In CD8^+^ T cells, 24% (6 of 25) of the P-interactome are related to transport (Figure 4e). All of these proteins belong to the key ion transport family, and about 16% are involved in calcium signalling (Figure 4e). Calcium signalling is important for ER-mediated stress response and protein folding^27^.

25% of the 442 LNCaP P5-interactome are related to transport of different classes (Figure S7). This includes three nuclear pore complex proteins, NUP93, NUP210 and NDC1, relevant to nuclear transportation, as well as TMED10, TMED9 and SEC22B, which belong to the ER and Golgi cisternae involved in vesicle trafficking.

We found enrichment of exosome-related proteins in both cell types. Exosomes are membrane-bound extracellular vesicles with key roles in intercellular communication and transport. Exosome exchange can regulate CD8^+^ T cell function and communication with other immune and tumour cells in the tumour microenvironment^28, 29^.

NPC1 and SCP2 are cholesterol binding membrane proteins essential for intercellular cholesterol trafficking in mammals. STEAP1, a highly expressed protein at prostate cancer cell-cell junctions also features in the P5 interactome. This protein is known to function as a channel or transporter at cell-cell junctions.

P5 modulates cytoskeleton by binding to cytoskeletal proteins^19, 30^. Overall about 16% of the P5-binding proteins are cytoskeletal (Figure 4c), which is consistent with P5’s reported role in cytoskeleton organization^19, 30^. Our P5-interactome capture revealed novel proteins such as Vimentin, an intermediate filament protein with a critical role in CD8^+^ T cell immune response^31^. We also find nuclear skeleton proteins associated with P5, including the lamins LMNA and LMNB1, which play a role in prostate cancer progression^32^.

## Discussion

Historically, P5 has been considered as an important molecule because it is the precursor of all functional steroid hormones, but the biological role of P5 itself is still emerging. Its immense importance as a bioactive molecule has been documented in many studies^4, 19, 33, 34^, yet its full interactome-map is unknown. Therefore, in this study we not only developed a method to profile P5-interacting proteins in live cells but also utilized it to reveal P5’s biochemistry in P5-sensitive cancer and P5-producing immune cells. This approach can be applied in any biological context where P5 has a role.

We chose prostate cancer cells as a model because prostate cancer is a major health concern globally^35, 36^, and previously it has been demonstrated definitively that P5 (and not P5 derivatives or P5-derived steroid hormones) drives LNCaP cell growth^10^. Moreover, intra-tumoural *de novo* steroidogenesis, either by neoplastic cancer cells^7, 37^, tumour-infiltrating immune cells^8^ or osteoblasts^38^ has been proposed as a driver of castrate-resistant prostate cancer. Therefore, we also studied P5-producing CD8^+^ T cells, which are primary anti-tumour effector cells in the tumour microenvironment.

Here, we developed biologically active P5 probes that mimic P5’s molecular, cellular and biological activities (Figure 2, S3 and S4). We exploited these probes to identify P5-binding proteins providing detailed insights of P5 function in prostate cancer and immune cells.

Among the three probes we synthesized, P5-C showed the most promising results, and structurally it is the best-suited probe for functional interactions, with both the hydroxyl and carbonyl groups available for reaction with protein molecules in the lysate. The probes enabled us to profile P5-binding proteins, which shed light on two major areas: (1) proteins that are involved in intra- and inter-cellular P5 trafficking and (2) how P5 may exert a biological effect via enzymatic activity as well as the cytoskeleton and membrane protein function.

The detailed mechanisms of how P5 is trafficked within and between cells has hitherto remained obscure. About 20% of the P5-interactome is related to transport (Figure 4c and e), which are relevant not only for P5’s own transport, but also affecting transport of other biomolecules (Figure S6). VDAC’s are mitochondrial membrane proteins mediating metabolite exchange between the mitochondria and cytosol. Cholesterol and neurosteroids, including P5 and its derivatives, are known to bind VDACs^18, 39^ and regulate their activity. Therefore, the presence of VDACs in the P5 interactome of both cell types suggests P5’s general mode of action is mediated by these ion channels.

P5’s interaction with cytoskeletal proteins is well established^19, 30^. The most interesting among the new cytoskeletal proteins we found is the intermediary filament protein Vimentin. This was round specifically in CD8+ T cells and has a key regulatory role in the immune response during T cell activation^31^.

P5’s mode of action seems to be mainly non-genomic, as reported previously for steroid hormones^15, 16, 40^. This is despite a radioactive P5 pull-down experiment that produced a signal around 110 kD^10^, a molecular weight corresponding to that of the Androgen Receptor (AR). We did not find evidence of any binding to transcription factors. Our detection of the unconventional P4 membrane receptors Progesterone Receptor Membrane Component (PGRMC) 1 and PGMRC2 (Table 1 and ST1) is intriguing, suggesting a possible regulation of P4 non-canonical membrane components by P5.

P5’s non-genomic role is evident from its interactome, which contains key proteins playing important roles in reprogramming the metabolic output of the cells. Cancer cells acquire metabolic flexibility, and one point of regulation is through a carnitine shuttle within the mitochondria^22^. The importance of lipid metabolism in prostate cancer has been well established^41^. In this context, the presence of carnitine palmitoyltransferase 1A and 2 (CPT1A and CPT2), vital enzymes of the fatty acid oxidation pathway^42^ and SLC25A20 the carnitine and acylcarnitine mitochondrial transporter (Figure 4f) is intriguing. The presence of several key enzymes of cholesterol biosynthesis (Figure 4g) and transport in P5-interactome indicates the allosteric role of P5 and feedback to its own synthesis pathway.

Consistent with the recognized role of P5 as a neurosteroid^43^, our study retrieved neurodegenerative pathway proteins (Figure 3f, and S7). Thus our results provide biochemical targets of P5 that are important not only to understand its function as a neurosteroid but also its regulatory role in neurodegenerative diseases.

Taken together, we have successfully mapped the P5 interactome and captured the general as well as cell type-specific binding of P5. Apart from P5’s emerging role in regulating cytoskeleton organization we find evidence of its potential role in key metabolic and transport pathways. This could lead to the understanding of steroid hormone regulation of prostate cancer progression and immune cell function. Our study demonstrates that a chemoproteomic approach can be extended to other *de novo* steroidogenic cell types (e.g. adipocytes, thymic epithelial cells, glial cells and neurons).

## Materials and Methods

### Chemical synthesis

#### General Methods

Solvents were purified and dried using standard methods prior to use: THF and Et2O by distillation from calcium hydride and lithium aluminium hydride; CH2Cl2, toluene and acetonitrile by distillation from calcium hydride. Petroleum ether 40-60 refers to petroleum ether distillate (BP = 40-60 °C). All other solvents were used as supplied unless otherwise stated. All reagents were used as supplied or purified using standard procedures as necessary. *n*-BuLi was titrated in triplicate using menthol and 1,10-phenanthroline in THF at 0°C prior to use. Flash column chromatography was performed using Breckland Scientific silica gel 60, particle size 40-63 nm under air pressure. All solvents used for chromatographic purification were distilled prior to use. Analytical thin layer chromatography (TLC) was performed using silica gel 60 F254 pre-coated glass backed plates and visualised by ultraviolet radiation (254 nm), potassium permanganate or ninhydrin as appropriate.^1^H NMR spectra were recorded on Bruker DPX-400 (400 MHz) or Bruker DRX-600 (600 MHz) spectrometers.

Chemical shifts are reported in ppm with the resonance resulting from incomplete deuteration of the solvent as the internal standard (CDCl3: 7.26 ppm). ^13^C NMR spectra were recorded on Bruker DPX-400 (100 MHz) or Bruker DRX-600 (150 MHz) spectrometers with complete proton decoupling. Chemical shifts are reported in ppm with the solvent resonance as the internal standard (^13^CDCl3: 77.2 ppm, triplet). Data are reported as follows: chemical shift δ ppm (integration (^1^H only), multiplicity (s = singlet, d = doublet, t = triplet, q = quartet, quin = quintet, br = broad, app = apparent, m = multiplet or combinations thereof, coupling constants J = Hz, assignment). ^13^C signals are singlets unless otherwise stated, where they are shown in the same format as ^1^H NMR.High resolution mass spectrometry (HRMS) was performed on a Waters Micromass LCT spectrometer using electrospray ionisation and Micromass MS software. HRMS signals are reported to 4 decimal places and are within 5 ppm of theoretical values.Infrared spectra were recorded neat as thin films on a Perkin-Elmer Spectrum One FTIR spectrometer andonly selected peaks are reported (s = strong, m = medium, w = weak, br = broad).

Chemical abbreviations: DIPA – diisopropylamine; DIPEA – *N*-*N*-diisopropylethylamine; DMAP – 4-dimethylaminopyridine; DMF – *N*-*N*-dimethylformamide; HBTU - (2-(1*H*-benzotriazol-1-yl)-1,1,3,3-tetramethyluronium hexafluorophosphate; PTSA – *para*-toluenesulfonic acid

#### Ethyl 3-oxohept-6-ynoate (1)

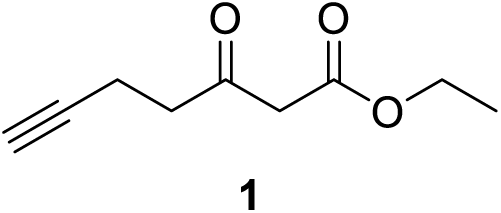

Diisopropylamine (42.0 mL, 300 mmol) was added to anhydrous THF (250 mL) under an atmosphere of argon and then cooled to 0 °C. *n-*BuLi (120 mL, 2.5 M in hexane, 300 mmol) was then added to the stirred solution dropwise *via* cannula syringe. After 15 min, ethyl acetoacetate (19.0 mL, 150 mmol) was added dropwise then left for 30 min. Propargyl bromide (16.7 mL, 150 mmol) was added dropwise to the solution and then stirred for 2 h at 0 °C. The solution was allowed to warm to room temperature then stirred for a further 15 min before quenching with glacial AcOH (9 mL) and the solution stirred for a further 15 min. The solution was diluted with Et2O (500 mL), washed with H2O (2 × 200 mL) and brine (200 mL) then dried over MgSO4. The solvent was removed *in vacuo* and the crude mixture was purified by distillation using a kugelrohr (1.7 Torr, 108 °C) to give **1** (10.8 g, 64.3 mmol, 43% yield) as a colourless oil.

IR (thin film, cm^−1^): 3284 (w), 1741 (s), 1714 (s); ^1^H NMR (600 MHz, CDCl3) δ = 1.29 (3H, t, J = 7.1 Hz), 1.96 (1H, t, J = 2.3 Hz), 2.48 (2H, td, J = 7.2 Hz, 2.5 Hz), 2.82 (2H, t, J = 7.3 Hz), 3.47 (2H, s), 4.21 (2H, q, J = 7.2 Hz); ^13^C NMR (150 MHz, CDCl3) δ = 12.9, 14.2, 41.8, 49.5, 61.6, 69.2, 82.5, 167.3, 200.8; HRMS (*m/z*): [M+Na]^+^calcd. for C9H12NaO3, 191.0679; found, 191.0678.

#### Ethyl 2-(2-(but-3-yn-1-yl)-1,3-dioxolan-2-yl)acetate (2)

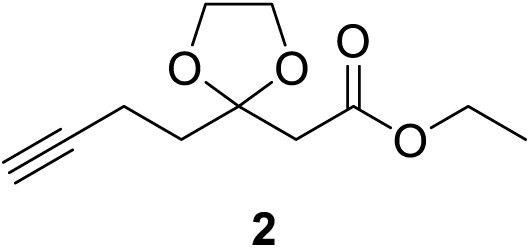

**1**(8.00 g, 47.6 mmol) was mixed with *para*-toluene sulfonic acid (0.24 g, 1.27 mmol) and ethylene glycol (13.3 mL, 238 mmol) in toluene (100 mL) in a dean stark apparatus and refluxed for 5 h. After this time the flask was cooled to room temperature then washed with sat. NaHCO3 (3 × 80 mL) and brine (30 mL) then dried over MgSO4. The solvent was removed *in vacuo*to give **2** (8.01 g, 37.8 mmol, 79% yield) as a pale yellow oil.

IR (thin film, cm^−1^): 3292 (w), 1731 (s); ^1^H NMR (600 MHz, CDCl3) δ = 1.17 (3H, t, J = 7.1 Hz), 1.86 (1H, m), 2.00 (2H, m), 2.21 (2H, m), 2.55 (2H, m), 3.90 (4H, m), 4.06 (2H, q, J = 7.1 Hz); ^13^C NMR (150 MHz, CDCl3) δ = 12.7, 14.1, 36.2, 42.5, 60.5, 65.1, 68.1, 83.8, 108.1, 169.1; HRMS (*m/z*): [M+Na]^+^calcd. for C11H16NaO4, 235.0941; found, 235.0955.

#### 2-(2-(But-3-yn-1-yl)-1,3-dioxolan-2-yl)ethan-1-ol (3)

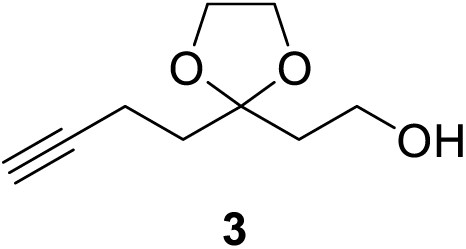

LiAlH4 (1.43 g, 37.7 mmol) was suspended in anhydrous ether (250 mL) under an atmosphere of argon and the stirred suspension cooled to 0 °C. **2** (8.00 g, 37.7 mmol) was added dropwise to the flask and then left for 15 min. To quench the residual LiAlH4, H2O (1.43 mL) was added slowly followed by a 15% (w/v) solution of NaOH (1.43 mL) and then H2O (4.29 mL). The mixture was stirred for 1 h until a white precipitate formed. MgSO4 was added to the flask then the suspension filtered. Solvents were removed *in vacuo* to give **3** (6.25 g, 36.8 mmol, 97% yield) as a pale yellow oil.

IR (thin film, cm^−1^): 3500-3200 (br w), 3288 (w); ^1^H NMR (600 MHz, CDCl3) δ = 1.93 (5H, m), 2.25 (2H, m), 2.71 (1H, br s), 3.72 (2H, m), 3.98 (4H, m); ^13^C NMR (150 MHz, CDCl3) δ = 13.1, 35.8, 38.2, 58.6, 64.9, 68.3, 83.9, 111.0; HRMS (*m/z*): [M+H]^+^ calcd. for C9H15O3, 171.1016; found, 171.1015.

#### 1-Hydroxyhept-6-yn-3-one (4)

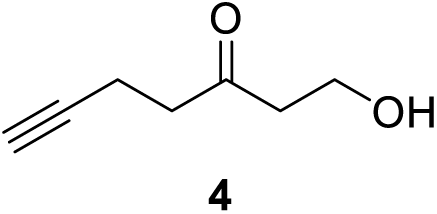

**3** (6.27 g, 36.8 mmol) was added dropwise to a solution of *para*-toluene sulfonic acid (1.75 g, 9.21 mmol) in acetone (100 mL) and H2O (1.5 mL) and stirred at room temperature for 5 h. The reaction was quenched with sat. NaHCO3 (150 mL), extracted with CH2Cl2 (5 × 150 mL) then the organic layer was washed brine (100 mL) and dried over MgSO4. Solvents were removed *in vacuo* to give **4** (4.33 g, 34.3 mmol, 93% yield) as a pale yellow oil.

IR (thin film, cm^−1^): 3550-3200 (br w), 3291 (w), 1708 (s); ^1^H NMR (600 MHz, CDCl3) δ = 1.94 (1H, t, J = 2.7 Hz), 2.43 (2H, td, J = 7.3 Hz, 2.7 Hz), 2.67 (5H, m), 3.83 (2H, t, J = 5.6 Hz); ^13^C NMR (150 MHz, CDCl3) δ = 12.7, 41.7, 44.6, 57.6, 64.9, 82.8, 209.1. HRMS (*m/z*): [M+H]^+^ calcd. for C7H11O2, 127.0754; found, 127.0753.

#### 2-(3-(But-3-yn-1-yl)-3H-diazirin-3-yl) ethan-1-ol (5)

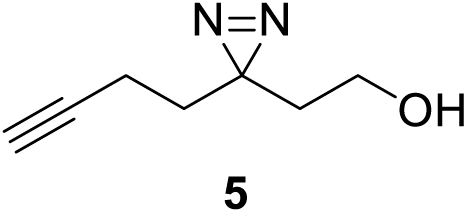

**4** (2.80 g, 22.2mmol) was added to a flask and placed under an atmosphere of argon then ammonia (30 mL) was condensed into the flask at −78 °C. The solution was refluxed at −35 °C for 5 h then cooled to −78 °C and a solution of hydroxylamine sulfonic acid (2.90 g, 25.6 mmol) in MeOH (20 mL) was added dropwise. The reaction was then refluxed at −35 °C for 1 h. The reaction was then allowed to warm to room temperature and left open to the atmosphere and stirred overnight. The suspension was then filtered, washing with MeOH with the liquor retained. The solvent was then removed *in vacuo* and the resultant solid dissolved in CH2Cl2 (10 mL). The solution was cooled to 0 °C and iodine (5.60 g, 22.2 mmol) was added portion wise until the solution retained a brown colour. The reaction mixture was diluted with CH2Cl2 (80 mL) then washed with 10% HCl solution (30 mL), sat. sodium thiosulfate (80 mL), H2O (40 mL) and brine (40 mL) respectively. The solution was then dried over MgSO4 and solvent removed *in vacuo* and the crude mixture purified using flash column chromatography (hexanes:EtOAc) to give **5** (0.965 g, 6.98 mmol, 32% yield) as a colourless oil.

IR (thin film, cm^−1^): 3550-3200 (br w), 3294 (w); ^1^H NMR (600 MHz, CDCl3) δ = 1.69 (5H, m), 2.00 (1H, t, J = 2.7 Hz), 2.04 (2H, td, J = 7.4 Hz, 2.7 Hz), 3.49 (2H, t, J = 6.2 Hz); ^13^C NMR (150 MHz, CDCl3) δ = 13.2, 26.6, 32.6, 35.5, 57.3, 69.2, 82.8; HRMS (*m/z*): [M-H]^−^calcd. for C7H9N5O, 137.0720; found, 137.0714.

#### 3-(But-3-yn-1-yl)-3-(2-iodoethyl)-3H-diazirine (6)

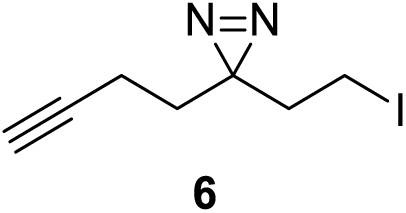

Iodine (1.54 g, 6.08 mmol) was added to a solution of PPh3 (1.46 g, 5.58 mmol) and imidazole (1.04 g, 68.1 mmol) in CH2Cl2 (16 mL) in a flask protected from light and stirred for 10 min at 0 °C. **5** (0.700 g, 5.07 mmol) in CH2Cl2 (2 mL) was then added and the reaction stirred for 4 h. The excess iodine was quenched by sat. sodium thiosulfate solution (40 mL) and the mixture extracted with EtOAc (2 × 50 mL). The organic layer was washed with H2O (40 mL) brine (40 mL) and dried over MgSO4. The solvents were removed *in vacuo* and the crude mixture purified using flash column chromatography (hexanes: 3% EtOAc) to obtain **6** (0.907 g, 3.65 mmol, 72% yield) as a colourless oil.

IR (thin film, cm^−1^): 3296 (w); ^1^H NMR (600 MHz, CDCl3) δ = 1.68 (2H, t, J=7.2 Hz), 2.02 (3H, m), 2.11 (2H, J=7.7 Hz), 2.88 (2H, t, J=7.7 Hz); ^13^C NMR (150 MHz, CDCl3) δ = −4.0, 13.2, 28.6, 31.8, 37.5, 69.4, 82.4.

#### 3-(2-Azidoethyl)-3-(but-3-yn-1-yl)-3H-diazirine (7)

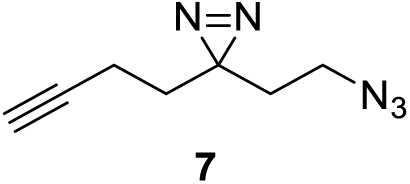

**6** (0.907 g, 3.65 mmol) was added to a solution of DMF (10 mL) and NaN3 (0.286 g, 4.40mmol) under an atmosphere of argon, then the solution was heated to 75 °C with stirring for 2 h. The reaction was allowed to cool to room temperature then was diluted with H2O (30 mL) and EtOAc (75 mL). The organic layer was retained and then washed with 5% (w/v) LiCl solution (2 × 30 mL), H2O (30 mL) and brine (30 mL) then dried over MgSO4. The solvent was removed *in vacuo* to give **7** (0.503 g, 3.07 mmol, 82% yield) as a pale yellow oil.

IR (thin film, cm^−1^): 3299 (w), 2093 (s); ^1^HNMR (600 MHz, CDCl3)δ = 1.69 (4H, m), 2.02 (3H, m), 3.16 (2H, t, J= 6.8 Hz);^13^CNMR (150 MHz, CDCl3) δ =13.2, 26.4, 32.2, 32.4, 45.9, 63.4, 82.5;HRMS (*m/z*): [M-H]^−^calcd. for C7H8N5, 162.0785; found, 162.0783.

#### 2-(3-(But-3-yn-1-yl)-3H-diazirin-3-yl)ethan-1-amine (8)

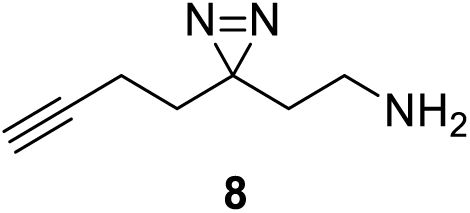

PPh3 (0.734 g, 2.80mmol) and **7** (0.400 g, 2.45 mmol) were added to a 10:1 solution of THF:H2O (5 mL) then stirred for 5 h at room temperature. 1 N HCl (10 mL) was added and the aqueous phase was washed withCH2Cl2 (2 × 15 mL). The solution was then made basic with 1 N NaOH (20 mL) and then extracted with CH2Cl2 (2 × 15 mL) and the organic layer dried overMgSO4. The solvent was removed *in vacuo* to give **8** (0.245 g, 1.79 mmol, 73% yield) as a pale yellow oil.

^1^HNMR (600 MHz, CDCl3)δ =1.14 (2H, s),1.65 (4H, m), 2.01 (3H, m), 2.51 (2H, t, J = 7.1 Hz);^13^C NMR (150 MHz, CDCl3) δ =13.3, 26.9, 32.6, 36.2, 36.7, 69.1, 82.7; HRMS (*m/z*): [M+H]^+^calcd. for C7H12N3, 138.1026; found, 138.1024.

In **Scheme 2A**, starting from the pregnenolone acetate (obtained from Sigma Aldrich) the synthesis of the probe **P5-B1** has been shown.

##### Pregnenolone (P5)

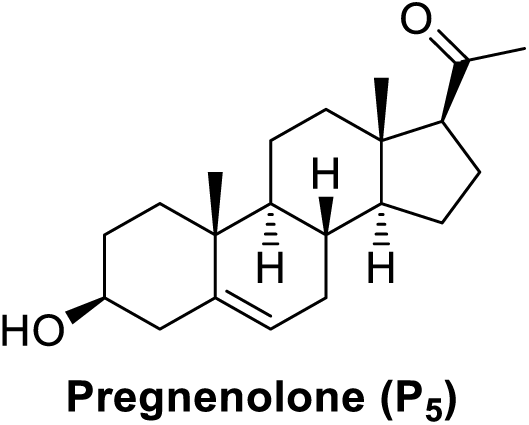

Pregnenolone acetate (1.00g, 2.80mmol) was added to *t*-BuOH (20 mL) which had been warmed to 42°C. To this an aqueous solution of KOH (1.71 mL, 1.06 g, 16.07 mmol) was added and the mixture was stirred overnight at 42 °C. The reaction was quenched with 5% HCl solution (30 mL) and extracted with Et2O (2 × 30 mL). The organic phase was dried with MgSO4, filtered and evaporated. The resultant white solid was recrystallized in methanol to obtain pregnenolone (0.761g, 2.40mmol, 86%yield).

IR (thin film, cm^−1^): 3507 (m), 1683 (s); ^1^H NMR (600 MHz, CDCl3) δ =0.68 (3H, s), 0.99 (1H, m), 1.01 (3H, s), 1.09-1.19 (2H, m), 1.24 (1H, m), 1.44-1.56 (5H, m), 1.60-1.71 (3H, m),1.86 (2H, m), 1.99-2.07 (2H, m), 2.14 (3H, s), 2.15-2.27 (2H, m), 2.35 (1H, m), 2.54 (1H, t, J = 9.0 Hz), 3.54 (1H, m), 5.36 (1H, m); ^13^C NMR (150MHz, CDCl3) δ = 13.2, 19.4, 21.1, 22.8, 24.5, 31.5, 31.6, 31.8, 31.9, 36.5, 37.3, 38.8, 42.3, 44.0, 50.0, 56.9, 63.7, 71.7, 121.4, 140.7, 209.5; HRMS (*m/z*): [M+Na]^+^calcd. for C21H32NaO2, 339.2295; found, 339.2290.

##### Compound 9

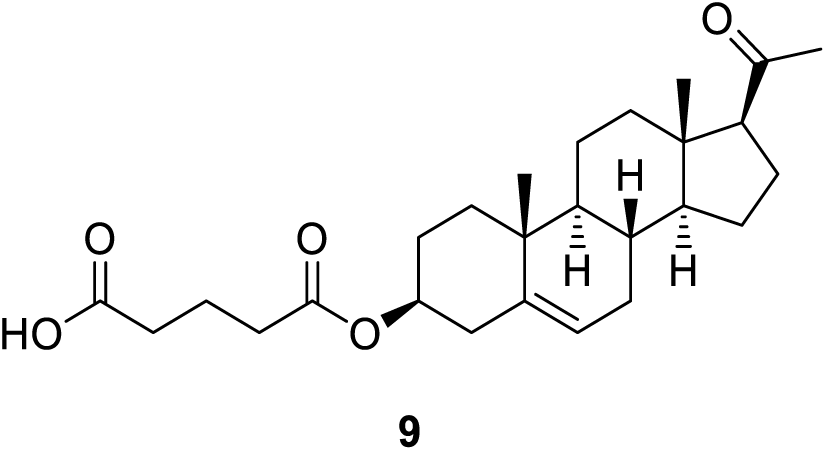

Pregnenolone (0.640 g, 2.02mmol), glutaric anhydride (0.920 g, 8.04mmol) and 4-(dimethylamino) pyridine (0.042 g, 0.34 mmol) were added to pyridine (6 mL) which had been placed under an atmosphere of argon. The solution was stirred overnight at 60°C then allowed to cool to room temperature.1 N HCl (15 mL) was added and then the solution was extracted with Et2O (2 × 40 mL). The organic layer was then washed once with water (40 mL) and brine (40 mL) then dried over MgSO4. The solvent was then removed *in vacuo* then purified by flash column chromatography (hexane:ethyl acetate, 4:1) to yield **9** (0.512g, 1.19 mmol, 57% yield) as an off white powder.

IR (thin film, cm^−1^):3100-2850 (br), 1729 (s), 1667 (s); ^1^H NMR (600 MHz, CDCl3) δ= 0.60 (3H, s), 1.00-1.04 (4H, m), 1.12-1.28 (3H, m), 1.42-1.52 (3H, m), 1.55-1.72 (5H, m), 1.87 (2H, m), 1.94-2.06 (4H, m), 2.13 (3H, s), 2.18 (1H, m), 2.32 (2H, m), 2.38 (2H, t, J = 7.3 Hz), 2.44 (2H, t, J = 7.3 Hz), 2.54 (1H, t, J = 9.0 Hz), 4.63 (1H, m), 5.38 (1H, d, J = 5.0 Hz);^13^C NMR (150MHz, CDCl3) δ = 13.2, 19.3, 19.9, 21.0, 22.8, 24.5, 27.7, 31.5, 31.7, 31.8, 32.9, 33.5, 36.6, 37.0, 38.1, 38.8, 44.0, 49.9, 56.8, 63.7, 74.0, 122.4, 139.6, 172.3, 178.5, 209.7.; HRMS (*m/z*): [M+Na]^+^calcd. for C26H38NaO5, 453.2611; found, 453.2609.

##### Probe P5-B1

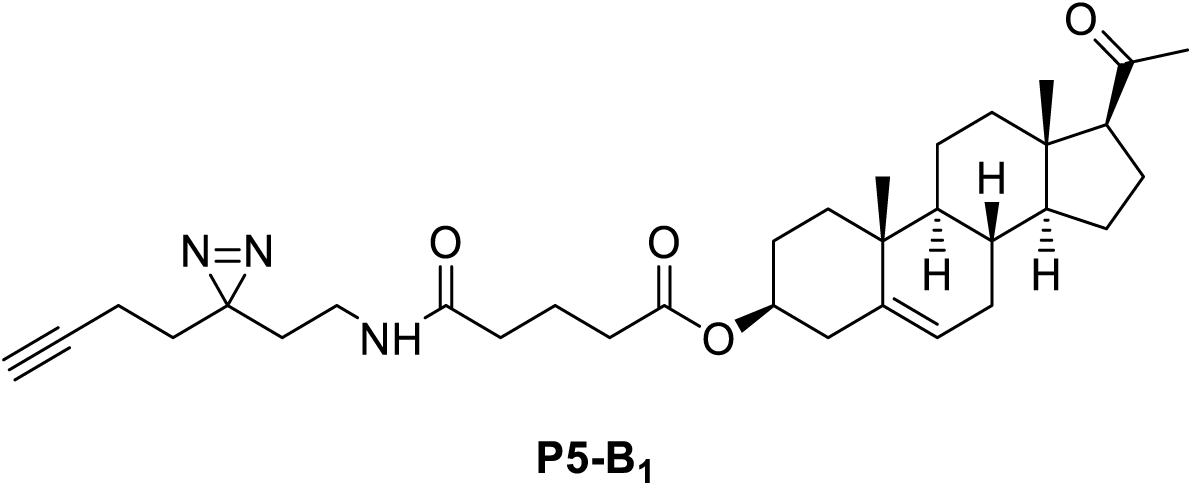

HBTU (0.232 g, 0.632mmol) and **9** (0.112 g, 0.252 mmol) were added to a flask and placed under argon. DMF (6 mL), DIPEA (0.110 mL, 0.632 mmol) and **8** (0.040 g, 0.292 mmol) were added by syringe. The solution was stirred for 16 h at room temperature then H2O (30 mL) was added to the solution then it was extracted with Et2O (2 × 40 mL). This was washed with 5% (w/v) LiCl (40 mL), H2O (3 × 30 mL), brine (40 mL) and then dried over Na2SO4. The solvent was removed *in vacuo* then purified by flash column chromatography (hexane:ethyl acetate: 7:3) to obtain **P5-B1** (0.053 g, 0.096 mmol, 38% yield). The product appeared approximately 90% pure by ^1^HNMR.

IR (thin film, cm^−1^):3352 (w), 3299 (m), 1728 (s), 1702 (s), 1648 (s); ^1^H NMR (600 MHz, CDCl3) δ =0.64 (3H, s), 1.03 (4H, m), 1.12-1.19 (2H, m), 1.20-1.24 (1H, m), 1.43-1.52 (3H, m), 1.56-1.64 (3H, m), 1.64-1.68 (4H, m), 1.69-1.72 (2H, m), 1.83-1.87 (2H, m), 1.94-2.08 (7H, m), 2.14 (3H, s), 2.19 (1H, m), 2.25 (2H, t, J = 7.4 Hz), 2.32 (2H, m), 2.37 (2H, t, J = 7.2 Hz), 2.54 (1H, t, J = 9.0 Hz), 3.11 (2H, app q, J = 6.3 Hz), 4.62 (1H, m), 5.38 (1H, d, J = 4.9Hz), 5.66 (1H, br s); ^13^C NMR (150 MHz, CDCl3) δ =13.2 (2C, m), 19.3, 20.9, 21.0, 22.8, 24.5, 26.8, 27.8, 31.5, 31.76, 31.80, 32.1, 32.5, 33.6, 34.3, 35.5, 36.6, 37.0, 38.1, 38.8, 44.0, 49.9, 56.8, 63.7, 69.4, 73.9, 82.7, 122.4, 139.6, 172.2, 172.6, 209.5;HRMS (*m/z*): [M+H]^+^calcd. for C33H48O4N3, 550.3639; found, 550.3632.

**Scheme 2B** depicts the synthesis of **P5-B2** starting from **10** which was synthesised by NewChem Technologies Ltd, Durham, UK.

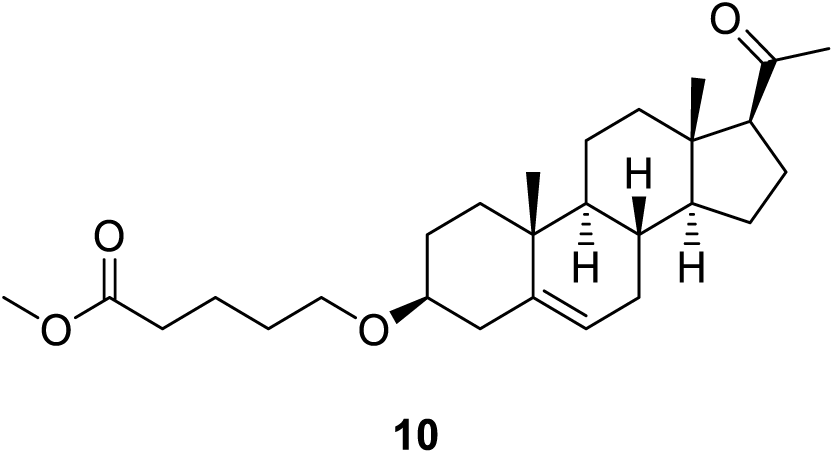

##### Compound 11

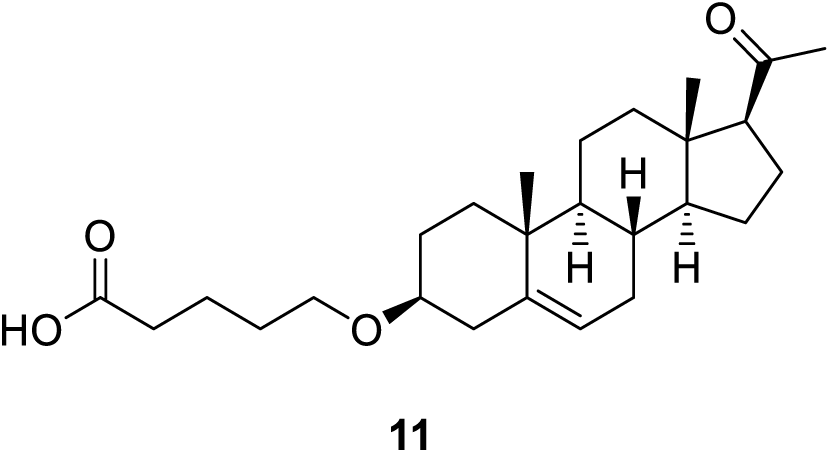

**10** (0.101 g, 0.230mmol) was added to a mixture of MeOH:THF:H2O1:1:1 (6 mL) LiOH.H2O (0.041 g, 0.977mmol) was added and the solution was stirred at room temperature for 2 h. Subsequently the reaction was acidified with 1M HCl solution (10 mL) then extracted using Et2O (3 × 40 mL). The organic layer was washed with H2O (40 mL), brine (40 mL) and dried over Na2SO4. The solvents were removed *in vacuo* to give **11** (0.084g, 0.201 mmol, 88% yield) as a white powder.

IR (thin film, cm^−1^):3300-2960 (br), 1731 (s), 1704 (s);^1^H NMR (600 MHz, CDCl3) δ = 0.60 (3H, s), 0.98 (4H, m), 1.04 (1H, m), 1.15 (1H, m), 1.21-1.27 (1H, m), 1.41-1.72 (12H, m), 1.88 (2H, m), 2.02 (2H, m), 2.12 (3H, s), 2.19 (2H, m), 2.38 (3H, m), 2.53 (1H, t, J = 9.0 Hz), 3.13 (1H, m), 3.49 (2H, m), 5.34 (1H, m); ^13^C NMR (150 MHz, CDCl3) δ = 13.2, 19.4, 21.1, 21.7, 22.8, 24.5, 28.4, 29.4, 31.5, 31.8, 31.9, 33.7, 36.9, 37.3, 38.8, 39.1, 44.0, 50.1, 56.9, 63.7, 67.5, 79.0, 121.2, 141.0, 178.7, 209.6; HRMS (*m/z*): [M+Na]^+^calcd. forC26H40NaO4, 439.2819; found, 439.2811.

##### Probe P5-B2

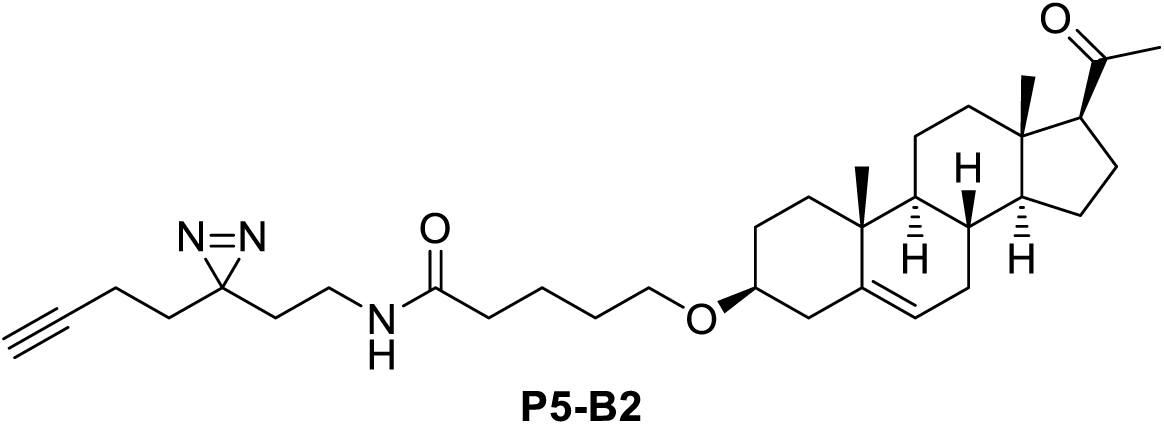

DMF (6 mL) was added to a flask containing **11** (0.080 g, 0.192mmol) and HBTU (0.174 g, 0.458mmol) that had been placed under an atmosphere of argon. DIPEA (0.08 mL, 0.459mmol) and **8** (0.032 g, 0.233mmol) were then added to the flask and it was stirred for 16 h. The reaction was then diluted with H2O (40 mL) then the solution extracted with Et2O (3 × 40 mL). The organic layer was washed with 5% (w/v) LiCl solution (60 mL), H2O (40 mL) and brine (40 mL) then dried over Na2SO4. The crude mixture was concentrated *in vacuo* then purified by flash column chromatography (hexane:ethyl acetate: 7:3) to give **P5-B2** (0.045 g, 0.084 mmol, 44% yield) as a colourless gum.

IR (thin film, cm^−1^): 3311 (m), 1702 (s), 1646 (s); ^1^H NMR (600 MHz, CDCl3)δ =0.62 (3H, s), 0.97 (1H, m), 0.99 (3H, s) 1.04 (1H, m), 1.11-1.18 (1H, m), 1.20-1.28 (2H, m), 1.41-1.52 (4H, m) 1.55-1.74 (13H, m), 1.84-1.91 (2H, m), 1.97-2.05 (5H, m), 2.11 (3H, s), 2.14-2.32 (4H, m), 2.36 (1H, m), 2.52 (1H, t, J = 9.0 Hz), 3.08-3.15 (3H, m), 3.45-3.50 (2H, m), 5.33 (1H, m), 5.73 (1H, br s);^13^C NMR (150 MHz, CDCl3) δ = 13.22, 13.23, 19.4, 21.1, 22.7, 22.8, 24.5, 26.8, 28.4, 29.5, 31.5, 31.78, 31.83, 32.2, 32.6, 34.2, 36.4, 36.9, 37.2, 38.8, 39.1, 44.0, 50.0, 56.9, 63.7, 67.7, 69.4, 79.0, 82.7, 121.2, 140.9, 172.9, 209.5; HRMS (*m/z*): [M+H]^+^calcd. ForC33H50N3O3, 536.3847; found, 536.3854.

Please note that within the proton NMR for this compound we have reported 51 proton peaks, whilst it contains only 49. It is believed this is due to a minor impurity within the sample. Due to the clustering of signals from the steroidal core, these cannot be isolated and therefore we have reported the spectra that are observed as we cannot determine where the extra protons are within the spectra. It should be noted that all other data is consistent for the reported product.

**Scheme 3** shows the synthesis of **P5-C** which starts from compound **12** which was synthesised by NewChem Technologies Ltd, Durham, UK.

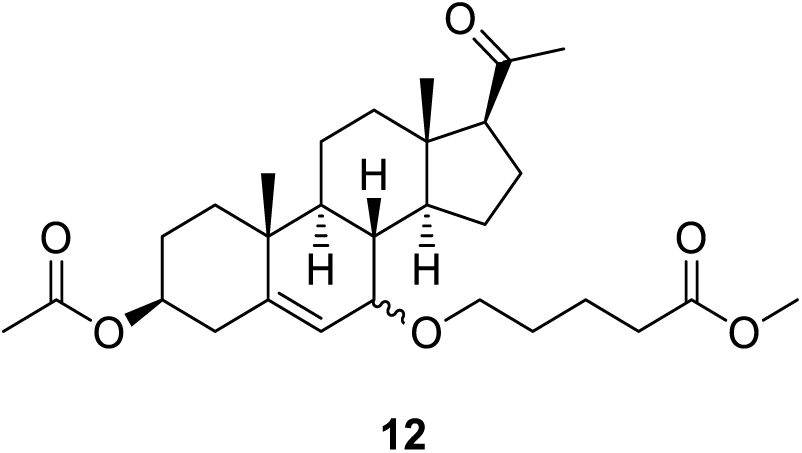

##### Compound 13

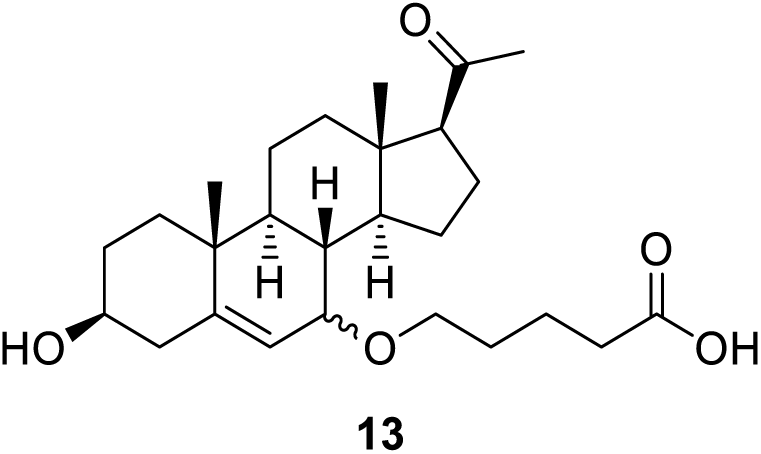

LiOH.H2O (0.052 g, 1.20mmol) was added to solution of **12** as a 9:1 mix of diastereomers (0.101 g, 0.204 mmol) in a solution of MeOH:THF:H2O 1:1:1 (6 mL) and stirred for 4 h. The solution was acidified with 1M HCl (15 mL) then the solution extracted with Et2O (3 × 30 mL). The organic layer was washed with H2O (40 mL) and brine (40 mL) then dried over Na2SO4. The solvents were removed *in vacuo* to give **13** (0.085g, 0.196 mmol, 98% yield) as colourless gum, as a mixture of the two diastereomers, with one predominant isomer (approximately 9:1 by ^1^HNMR based on protons at position 6 of the steroid core).

IR (thin film, cm^−1^):3550-2990 (br), 1701 (s, with shoulder); ^1^H NMR (600 MHz, CDCl3) δ = 0.62 (3H, s), 0.98 (3H, s), 1.14-1.26 (3H, m), 1.41 (1H, m), 1.45-1.55 (4H, m), 1.59-1.76 (9H, m), 1.83-1.88 (2H, m), 2.00 (1H, m), 2.14 (3H, s), 2.18 (1H, m), 2.27-2.36 (2H, m), 2.39 (2H, t, J = 7.4 Hz), 2.61 (1H, t, J = 8.9 Hz), 3.32 (1H, dt, J = 9.1 Hz, 6.3 Hz), 3.40 (1H, m), 3.58-3.67 (2H, m), 5.68 (1H, dd, J = 4.9 Hz, 1.6 Hz); ^13^CNMR (150 MHz, CDCl3) δ = 12.9, 18.3, 20.8, 21.8, 22.8, 24.5, 29.5, 31.4, 31.6, 33.6, 36.8, 37.2, 37.4, 38.2, 42.3, 42.6, 43.8, 49.3, 63.6, 68.3, 71.4, 72.3, 121.3, 145.8, 178.0, 209.9; HRMS (*m/z*): [M+Na]^+^calcd. For C26H40NaO5, 455.2768; found, 455.2752.

Due to diastereotopic mixture the NMR cannot be comprehensively assigned. As such the peaks described here are those observed.

##### Probe P5-C

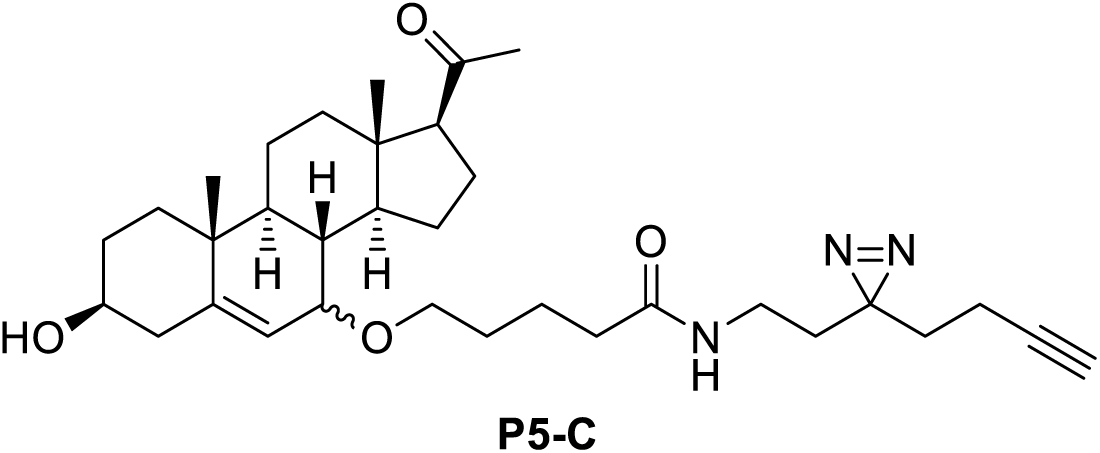

A flask containing **13** (0.085 g, 0.196mmol) and HBTU (0.182 g, 0.480mmol) was placed under an atmosphere of argon then DMF (6 mL), DIPEA (0.084 mL, 0.482mmol) and compound **8** (0.033 g, 0.242mmol) were added. The solution was stirred for 16 hat room temperature. The reaction was diluted with H2O (40 mL) then extracted with Et2O (3 × 40 mL). The organic layer was washed with 5% (w/v) aqueous LiCl solution (60 mL), H2O (40 mL) and brine (40 mL) respectively then dried over Na2SO4. The solvents were removed *in vacuo* and then purified by flash column chromatography (hexane:ethyl acetate: 7:3) was performed to give**P5-C** (0.075 g, 0.136 mmol, 68% yield) as a yellow gum.

IR (thin film, cm^−1^): 3305 (m), 1697 (s), 1649 (s); ^1^H NMR: (600 MHz, CDCl3)δ = 0.62 (3H, s), 0.98 (3H, s), 1.15 (1H, m), 1.22-1.31 (1H, m), 1.36-1.41 (1H, m), 1.42-1.54 (4H, m), 1.55-1.74 (14H, m),1.83-1.90 (2H, m), 1.99-2.05 (4H, m), 2.14 (3H, s), 2.15-2.24 (3H, m), 2.27-2.36 (2H, m), 2.60 (1H, t, J = 9.0 Hz), 3.11 (2H, appq, J = 6.4 Hz), 3.30 (1H, m), 3.40 (1H, m), 3.57-3.67 (2H, m), 5.62 (1H, brm), 5.69 (1H, m); ^13^C NMR (150 MHz, CDCl3) δ = 12.9, 13.2, 18.2, 20.8, 22.7, 22.9, 24.5, 26.9, 29.7, 31.4, 31.6, 32.1, 32.6, 34.3, 36.4, 36.8, 37.2, 37.4, 38.2, 42.3, 42.6, 43.8, 49.4, 63.5, 68.6, 69.4, 71.3, 72.2, 82.7, 121.3, 145.8, 172.9, 209.8; HRMS (*m/z*): [M+H]^+^calcd. forC33H50N3O4, 552.3796; found, 552.3790.

Due to diastereotopic mixture the NMR cannot be comprehensively assigned. As such the peaks described here are those observed.

**Scheme 4** is the diagrammatic representation for the synthesis of **Probe P5-A** from **14** which was obtained from NewChem Technologies Ltd, Durham, UK.

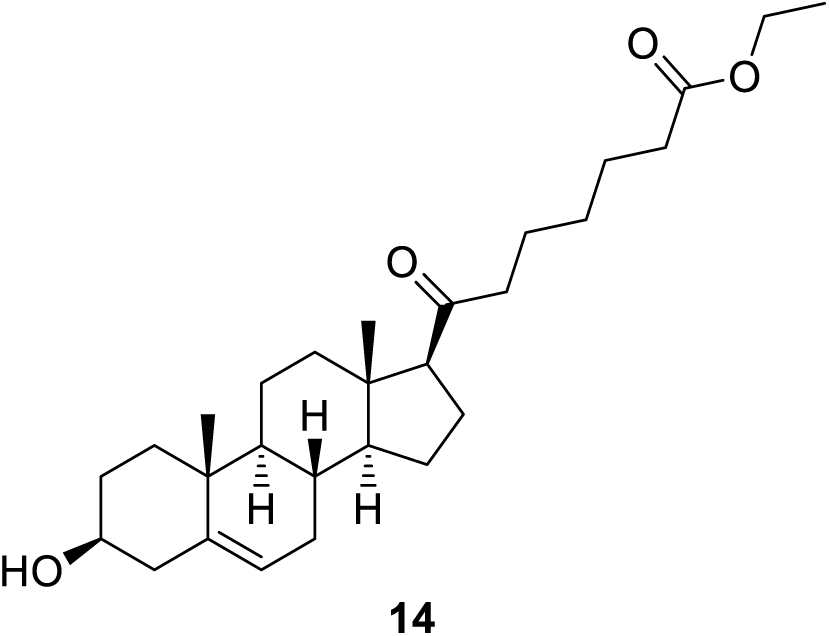

##### Compound 15

##### Probe P5-A

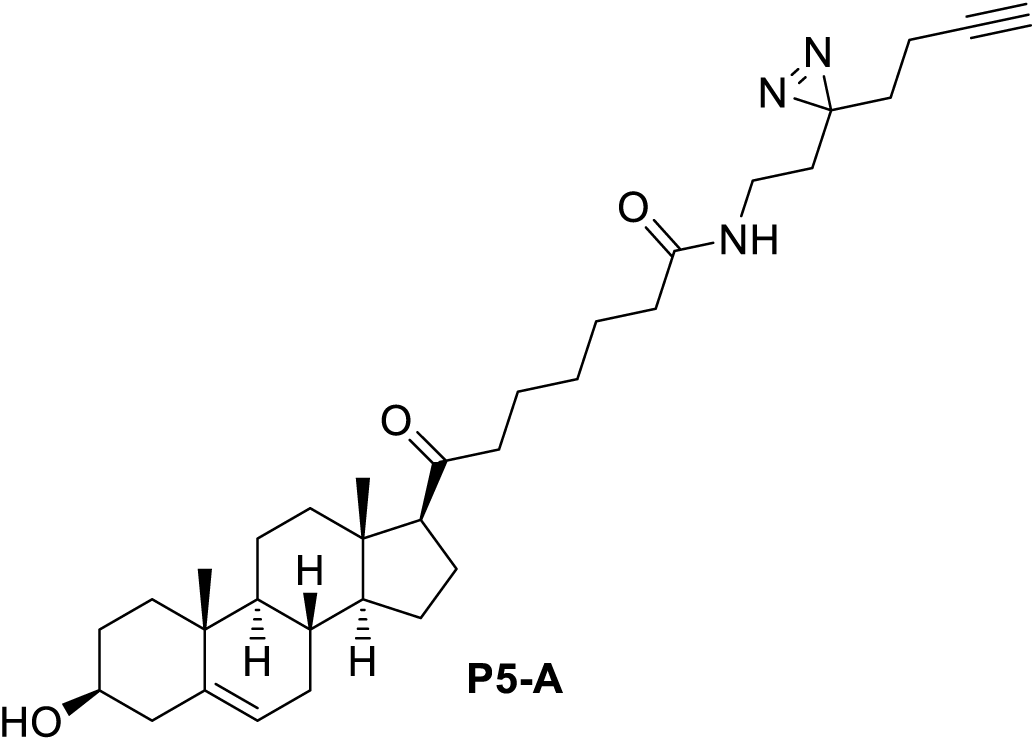

LiOH.H2O (0.052 g, 1.24mmol) was added to solution of**14** (0.100 g, 0.224 mmol) in a solution of MeOH:THF:H2O 1:1:1 (6 mL) and stirred for 4 h. The solution was acidified with 1 N HCl (15 mL) then the solution extracted withEt2O (3 × 30 mL). The organic layer was washed with H2O (40 mL) and brine (40 mL) then dried over Na2SO4. The solvents were removed *in vacuo* to give**15** as a white powder that was used immediately in the next step. HRMS (*m/z*): [M+H]^+^calcd. for C26H41NO4, 417.2999; found, 417.3004.

A flask containing the crude solid of **15** and HBTU (0.182 g, 0.483mmol) was placed under an atmosphere of argon then DMF (6 mL), DIPEA (0.084 mL, 0.483mmol) and **8** (0.033 g, 0.242mmol) were added. The solution was stirred for 16 h at room temperature. The reaction was diluted with H2O (40 mL) then extracted with Et2O (3 × 40 mL). The organic layer was washed with 5% (w/v) aqueous LiCl solution (60 mL), H2O (40 mL) and brine (40 mL) respectively then dried over Na2SO4. The solvents were removed *in vacuo* and then purified by flash column chromatography (hexane:ethyl acetate: 7:3) was performed to give **P5-A** (0.073 g, 0.136 mmol, 61% yield over two steps) as a colourless gum.

IR (thin film, cm^−1^): 3308(m), 1701 (m), 1647 (s); ^1^H NMR: (600 MHz, CDCl3) δ = 0.62 (3H, s), 0.67 (0.7H, s), 0.99 (1H, m), 1.01 (3.7H, s), 1.08-1.17 (2.6H, m), 1.18-1.35 (4.3H, m), 1.42-1.77 (21.8H, m), 1.85-1.89 (2.6H, m), 1.99-2.05 (6.2H, m), 2.18-2.26 (3.9H, m), 2.32-2.36 (2.6H, m), 2.39 (2H, t, J = 7.1 Hz), 2.51 (1H, t, J = 8.8 Hz), 3.11 (2.4H, app q, J = 6.4 Hz), 3.52 (1.2H, m), 4.06 (0.4H. m), 5.40 (1.2H, m), 5.60 (1.2H, br s);^13^C NMR (150 MHz, CDCl3) δ = 13.2, 13.4, 19.4, 21.1, 23.0, 23.1, 24.5, 25.4, 26.8, 28.7, 31.6, 31.76, 31.84, 32.2, 32.5, 34.2, 36.4, 36.5, 37.3, 38.9, 42.2, 44.0, 44.2, 50.0, 57.0, 62.9, 69.4, 71.7, 82.7, 121.4, 140.8, 172.9, 211.5; HRMS (*m/z*): [M+H]^+^calcd. forC33H50N3O3, 536.3847; found 536.3859.

The product is a 5:1 mixture of the reported product and an unknown impurity. As such the peaks described here are those observed with partial integrals included. **P5-A** was used in subsequent experiments despite this known impurity and the appropriate controls were put in place to ensure data obtained with this probe was valid.

#### Cell Culture and reagents

The human androgen sensitive prostate cancer cell line LNCaP, glioblastoma cell line U87MG (Gift from Dr Mathew Garnett, Wellcome Sanger institute) were first checked for mycoplasma (Surrey Diagonostics). The mycoplasma free LNCaP, U87MG and HEK293T cells were cultured in RPMI 1640 (supplemented with 10% FBS and 1% penicillin-streptomycin), DMEM/F-12 (supplemented with 10% charcoal stripped FBS and 1% penicillin-streptomycin) and DMEM (supplemented with FBS and 1% penicillin-streptomycin) respectively. Pregnenolone and ethanol, were obtained from Sigma. Stock solutions, 100mM of A, B1, B2, and C and 50mM of pregnenolone in ethanol were prepared. For LNCaP cells, 5000 cells per well were cultured in 100 μL of complete medium in a 96 well flat-bottom clear poly-L-lysine coated plates for 24 hrs at 37°C. After a day, the medium was replaced with phenol-red free RPMI 1640, supplemented with charcoal stripped FBS and 1% penicillin-streptomycin. The cells were grown either in presence of 2 to 20nM of indicated probes or the vehicle, ethanol. The cultures were maintained for 6 days and replacing the medium every 48 hours. For U87MG cells, 5000 cells were cultured in 100μL of medium in 96 wells flat-bottomed clear plates. The next day the medium was replaced and appropriate amount of either the probes or the vehicle ethanol was added. The culture was maintained for 4 days with medium replacement every 48 hrs.

#### Cloning and expression of CLIP1

The sequence verified cDNA clone of human CLIP1 (from Dharmacon) was PCR amplified with primers CLIP FW – TTTTGGCCCACAGGGCCTTAGTATGCTAAAGCCAAGTGG, CLIP RW - TTTTGGCCGATATGGCCTCAGAAGGTTTCGTCGTCATTGC and subcloned in frame into a mammalian expression vector (kind gift from Martin Beck, EMBL) with triple tag (HA, StrepII and His) after digestion with Sfi I (NEB) and extraction from agarose gel (using Qiagen kit). The correct incorporation was verified by sequencing and then transfected into HEK293T cells using Lipofectamine (Invitrogen) as per the manufacturer’s protocol. The transfected cells were harvested after 48 hours, washed once with PBS and stored at −80°C until further use.

#### Protein Binding Assay and proteomics

To test the *in vitro* binding of the probes (**P5-A, P5-B1, P5-B2** and **P5-C**) to CLIP1, the protein was transiently expressed in HEK cells. The transfected HEK cells were homogenized in 10 mM Tris pH 8, 1.5 mM EDTA, 10% glycerol, 1 μl Phosphatase and 1 μl Protease inhibitor (from ThermoFisher) using a 26G ½ needle fitted in a 1mL syringe. Roughly 400 μg of clear whole cell lysate (WCL) (measured in a NanoDrop spectrophotometer) was mixed with 50 nM probes and incubated at 37°C for 1 hour. The mixture was then irradiated at 365nm with a lamp (from UVP) on ice and then clicked with 100 nmoles magnetic azide (Turbobeadsazide, Sigma) in the presence of 1 mM CuSO4 (Sigma), 0.1 mM TBTA (Sigma) and 1mM TCEP (Sigma) for 1 hour at 37°C in 500 μL of PBS/0.1% SDS. A blank assay without the probe was carried out with all the above steps to ascertain the background binding to the magnetic beads. In a parallel competition assay, pregnenolone was added to the WCL and incubated at 37°C for 30 minutes before adding the probe and then the above procedure was followed. After clicking the beads were pulled down washed thoroughly 3 times with 1 mL PBS/0.1% SDS and then 3 times with 1 mL PBS. Subsequently, the beads were suspended in the SDS buffer and boiled for 10 minutes at 95°C. The samples were loaded on a 7.5% TGX gels (BioRad) and after electrophoresis the proteins were transferred to a nitrocellulose membrane using a wet transfer unit (BioRad). The membrane was blocked for 4 hours in 5% non-fat dry milk in 1X PBS and then incubated overnight at 4°C in 1X PBS with 5% non-fat dry milk and HA primary antibody (Abcam). After 3 washes with 1X PBS-0.05% Tween-20 the membrane was incubated with anti-rabbit secondary antibody (Santa Cruz Biotechnology) for 1 hour at RT. After 4 five minutes washes the membrane was incubated with SuperSignal West Dura (ThermoFisher) for five minutes and the resulting chemiluminescence was captured using the LAS4000 Image Quant (GE).

To test the global binding efficiency and specificity, probe C was used with LNCaP cells extracts. Three samples were prepared in parallel where 400 μg of protein, measured in a NanoDrop spectrophotometer was incubated with 10 μM of **P5-C** at 37°C for 30 minutes, irradiated with 365 nm UV for 30 minutes at 4°C and then clicked with 0.5 mM Biotin Azide (Sigma) in presence of 1 mM CuSO4 (Sigma), 0.1mM TBTA (Sigma) and 1mM TCEP (Sigma) for 1 hour at 37°C in 500μL of PBS/0.1 % SDS. In the second one 100 μM (10X) pregnenolone was added to extract and incubated for 1 hour at 37 °C before adding the probe, and the third sample was prepared without the UV irradiation. Another sample was prepared without the probe to determine the background binding of the beads. To get rid of the excess probes and biotin, 1 volume of the protein samples were washed with chloroform:methanol:water (1:4:3 volumes) and after removal of the aqueous top layer the protein containing interface was precipitated with 4 volumes of methanol. The precipitate was dried for 10 minutes and then resuspended in 200μL of PBS/1%SDS at 70°C for 10 minutes. Once the protein pellet was completely dissolved, the SDS concentration was diluted to 0.2% with PBS. The solution was centrifuged at 3000 × g for 3 minutes at RT to remove any undissolved particles and then the supernatant was mixed with 100 μl of 50 % slurry of neutravidin beads (ThermoFisher) pre-equilibrated in PBS/0.2% SDS. The mixture was incubated with slow rotation at RT for 1 hour. The beads were collected at 100 × g for 1 minute and then washed 3 times with 1 mL PBS/0.1% SDS, and then 3 times with only PBS. 25 μL of 2 × Laemmlibuffer was added to beads and boiled at 95 °C for 5 minutes, the supernatant was then loaded on to a 10% PAGE with SDS. The gel was stained with silver (ThermoFisher) as per manufacturer’s protocol and imaged LAS4000 Image Quant (GE).

For proteomic experiments, on the one hand 10μM of all 4 probes were incubated with 10 million LNCaP cells in phenol red free RPMI1640 for 1 hour in a 24 well plate alongside cells with no probes. The media was removed, cells washed once with cold PBS and then the cells were irradiated at 365nm for 15 minutes in 200 μL of cold PBS at 4°C. After removing the PBS the cells were washed once with PBS and then were extracted for proteins with PBS containing 0.2% SDS. After centrifugation, the clear lysates were used for copper-click and affinity purification. Following 3 times PBS/0.1% SDS, the beads were washed 3 times with 1 mL of PBS and then LNCaP cells were sent to the Wellcome Sanger Institute proteomics facility for on-bead digestion and MS sequencing. In parallel, to complement the above experiment, two samples, one with only 10 μM **P5-C** and another competition assay, where live cells were fed with 100μM (10X) P5 before adding the probe, were treated identically as above. After washing the beads with PBS/0.1%SDS the beads were boiled in Laemmli buffer at 95 °C for 5 minutes. The samples were loaded in adjacent lanes of an SDS-PAGE gel. After completion of the run the gel was stained with colloidal Coomassie Brilliant Blue. Each lane was cut into 4 pieces and from each piece and proteins were extracted, digested and sequenced in the Wellcome Sanger Institute’s proteomics facility.

10 million murine CD8^+^ T cells were used in three different experiments with two replicates for each. In the first experiment, CD8^+^ T cells in phenol-free RPMI were incubated with only the vehicle (without probe), the second experiment had 10 µM **P5-C** and the third was incubated with 100 µM (10X) of P5 before 10 µM of **P5-C** incubation. The following protocol was the same for all the experiment and its replicates. The cells were centrifuged, washed twice with ice cold PBS and then exposed to UV and the subsequent cell lysate preparation, click reaction and pull-down procedure was same as for LNCaP cells in the above section. The washed neutravidin beads were sent to EMBL proteomics core facility for TMT labelling, peptide fractionation and mass spectrometry.

#### Cell Viability Assay

XTT assay (Cayman Chemical) was used for assessing the effect of the probes on cell proliferation. For LNCaP cells, after 6 days a background absorbance, a reading was taken at 485 nm before adding 50 µL of activated XTT reagent to each of the wells. The cells were incubated at 37°C in an incubator for 5-6 hours before measuring the absorbance at 485nm. For U87MG cells, after 4 days of culture, the medium was replaced by sterile 100 μL of phosphate buffered saline, and then a background reading was measured at 485nm. 50 μL of activated XTT reagent was then added to each well and the plate was incubated at 37°C for 6 hours. The absorbance was measured at 485nm in a plate reader (Biotek).

#### Mass Spectrometry

##### LNCap Cell proteomics

For gel-LC/MS samples, the gel lane was excised to 4 bands, and destained with 50% CH3CN/50% of 50mM ammonium bicarbonate, then digested by trypsin (Roche) overnight. Peptides were then extracted by 50% CH3CN/50% of 0.5% formic acid, then dried in SpeedVac.

For on-bead digestion samples, beads were washed with 100mM TEAB twice, then resuspended in 250 µl of 100mM TEAB buffer. 2µl of 500mM TCEP and 2µl of 500mM iodoaceamide were added then incubated at 25°C for 45minutes. Then 0.4 µg of trypsin was added and the mixture was incubated at 37°C for 18 h. Beads were then transferred to a spin column, centrifuged at 400 × g for 1 min to collect the flow through. 100 µl of 1M TEAB was added to the beads again and sand collect as above, and both collections were pooled, then dried in SpeedVac.

Peptides were resuspended in 0.5% formic acid before LC-MS/MS analysis on an Ultimate 3000 RSLC nano System (Dionex) coupled to an LTQ Orbitrap Velos (Thermo Fisher) mass spectrometer equipped with a nanospray source.

The peptides were first loaded and desalted on a PepMap C18 trap (0.1 mm id × 20 mm, 5µm) then separated on a PepMap 75 µm id × 25 cm column (2 µm) over a 60 min linear gradient of 5– 42% B / 90 min cycle time, where B is 80% CH3CN/0.1% FA for gel band sample, but 5-40%B in 120min /150min cycle time for on-bead digestion sample. The LTQ Orbitrap Velos was operated in the “top 10” data-dependant acquisition mode while the preview mode of FT master scan was enabled. The Orbitrap full scan was set at m/z 380 – 1500 with the resolution at 30,000 at m/z 400 and AGC at 1×10^6^ with a maximum injection time at 200 msec. The 10 most abundant multiply-charged precursor ions, with a minimal signal above 2000 counts, were dynamically selected for CID fragmentation (MS/MS) in the LTQ ion trap, which has the AGC set at 5000 with the maximum injection time at 100ms.The dynamic exclusion was set at ± 10 ppm for 45 sec.

Data from gel bands were processed in Proteome Discoverer 1.4 (Thermo) using the Mascot (V 2.5) search engine against a Uniprot human database (version May 2013) combined with the common contaminate database. The precursor mass tolerance is set at 20 ppm, and fragments at 0.5 Da. Trypsin with full specificity and 2-missed cleavage sites was used. The dynamic modifications are set as acetyl (protein N-term), carbamidomethyl (C), deamidated (NQ) and Oxidation (M). The FDR setting used q-value, where the strict is at 0.01, and relaxed is at 0.05. Only peptides at high confidence were selected for protein groups.

Data from on-bead digestion were processed by MaxQuant (version 1.5.3.30) (http://www.coxdocs.org/). The human protein database was downloaded from Uniprot (version June 2016) and a common contaminate database was used as well. The parameters were the default except the carbamidomethyl (C) was set as variable as above. Other variable modifications were also the same as above. Both PSM and protein FRD were set at 0.01.

##### For CD8+ cell proteomics

###### Sample preparation and TMT labeling

The supernatant was removed from the beads and 20 µl of 4x Laemmli buffer was added to the ∼ 60 µl beads, vortexed, and kept shaking for 15 minutes at 95°C, vortexed again and subsequently kept shaking for additional 15 minutes at 95°C. The samples were cooled to room temperature and filtered using Mobi columns with a 90 µm filter.

The resulting samples (25 µl) were diluted with 50 µl 50 mM HEPES solution and treated with 2 µl 200 mM dithiothreitol in 50 mM HEPES at pH 8.5 for 30 min at 56°C to reduce disulfide bridges. The accessible cysteine residues were carbamidomethylated for 30 min in the dark after addition of 4 µl 400 mM 2-chloroacetamide in 50 mM HEPES at pH 8.5.

Protein clean up and digestion was done by SP3^44^. Two microliter of a 1:1 mixture of hydrophilic and hydrophobic Sera-Mag SpeedBeads (ThermoFisher) prewashed with water and at a concentration of 10 µg/µl were added to each sample. After addition of 83 µl acetonitrile, the suspensions were kept for 8 min prior to putting the vials on the magnet for 2 more minutes. The supernatants were discarded and the beads were washed twice with 200 µl 70% ethanol and once with 180 µl acetonitrile. After removing the acetonitrile, the beads were dried on air. After addition of 150 µg trypsin in 10 µl 50 mM HEPES buffer, the bound proteins were digested overnight. On the next day, the bead suspensions were sonicated for 5 minutes and vortexed prior to putting the vials on the magnet. The supernatants containing the peptides were transferred to new vials. The beads were rinsed with 10 µl 50 mM HEPES buffer and the resulting supernatants were combined with the first 10 µl. The individual samples were labeled by addition of 4 µl TMT-6plex reagent (ThermoFisher) in acetonitrile and incubated for one hour at room temperature. The reactions were quenched with a 5% hydroxylamine solution and acidified with 50 µl 0.05% formic acid. The 6 samples of each replicate were combined and the resulting two 6-plex samples were cleaned using an OASIS HLB µElution Plate (Waters). The wells were first washed twice with 0.05% formic acid in 80% acetonitrile and twice with 0.05% formic acid in water. The samples were loaded on the wells and washed with 0.05% formic acid in water. After elution with 0.05% formic acid in 80% acetonitrile the samples were dried and reconstituted in 4% acetonitrile and 1% formic acid in water.

###### Peptide fractionation

After adjusting the pH of the samples to pH 10 with ammonium hydroxide, the TMT-labeled peptides were fractionated on an Agilent 1200 Infinity HPLC system equipped with a degasser, quaternary pump, autosampler, variable wavelength UV detector (set to 254 nm), and fraction collector. Separation was performed on a Phenomenex Gemini C18 (100 × 1.0 mm; 3 μm; 110 Å) column using 20 mM ammonium formate pH 10 in water as mobile phase A and 100% acetonitrile as mobile phase B. The column was used in combination with a Phenomenex Gemini C18, 4 × 2.0 mm SecurityGuard cartridge. The flow rate was 0.1 ml/min. After 2 min isocratic separation at 100% A, a linear gradient to 35% B at minute 59 was used, followed by washing at 85% B and reconstitution at 100% A. In total 32 two minute fractions were collected and pooled to result in 6 samples. These were dried and reconstituted in 4% acetonitrile and 1% formic acid.

###### Mass spectrometry data acquisition

The fractionated samples were analyzed on an UltiMate 3000 nano LC system (Dionex) coupled to a QExactive plus (Thermo) mass spectrometer via a Nanospray Flex source (Thermo) using a Pico-Tip Emitter (New Objective; 360 µm OD × 20 µm ID; 10 µm tip). The peptides were first trapped on a C18 PepMap 100 µ-Precolumn (300 µm × 5 mm, 5 µm, 100 Å) prior to separation on a Waters nanoEase C18 75 µm × 250 mm, 1.8 µm, 100 Å column. The applied flow rates were 30 µl/min for trapping and 300 nl/min for separation. The mobile phase A was 0.1% formic acid in water and the mobile phase B was 0.1% formic acid in acetonitrile. After an initial isocratic step at 2% B for 2.9 minutes the multi-step gradient started with a gradient to 4% B at minute 4 followed by a linear increase to 8% B at minute 6. Subsequently, a shallow gradient to 28% B at minute 43 was followed by a steep gradient to 40% B at minute 52, a washing step at 80% B, and reconstitution at 2% B.

All spectra were acquired in positive ion mode. Full scan spectra were recorded in profile mode in a mass range of 375-1200 *m/z*, at a resolution of 70,000, with a maximum ion fill time of 10 ms and an AGC target value of 3×10^6^ ions. A top 20 method was applied with the normalized collision energy set to 32, an isolation window of 0.7, the resolution at 17,500, a maximum ion fill time of 50 ms, and an AGC target value of 2×10^5^ ions. The fragmentation spectra were recorded in profile mode with a fixed first mass of 100 *m/z*. Unassigned charge states as well as charge states of 1, 5-8, and >8 were excluded and the dynamic exclusion was set to 30 seconds.

#### Th2 Cell Proliferation Assay

We followed the method as described previously in Ref^45^. Negatively purified splenic naive CD4^+^ T cells were stained with CellTrace Violet following the CellTrace Violet Cell Proliferation Kit (Invitrogen) protocol and cultured under Th2 activation/differentiation conditions as described previously in the presence or absence of pregnenolone and linker tagged pregnenolene (**P5-A, P5-B1, P5-B2** and **P5-C**) for 3 days. The cell proliferation profile was captured by a flow cytometry-based dye decay assay on BD Fortessa. Data were analyzed in FlowJo.

#### Immunoglobulin Class Switch Recombination Assay

The detailed method is described previously^4^. Splenic naïve B cells from 8-12 week-old mice were purified by depletion of CD43^+^ cells using anti-CD43-coupled magnetic microbeads (Miltenyi Biotec) and seeded into 96-wellplates in RPMI supplemented with 10% FBS, 0.05 mM 2-mercaptoethanol, 25 ng/ml recombinant mouse IL4 (R&D Systems), and 40μg/ml LPS (Sigma-Aldrich). On day 5 of stimulation, B cell Fc receptors were blocked with PBS containing 2% rat serum and 10 mM EGTA and stained with FITC-conjugated anti-IgG1 (BD Biosciences). Flow cytometry was performed using a Fortessa (BD). Data were analyzed with FlowJo software.

#### Analysis of the MS results

After the LC-MS/MS data was obtained it was filtered to classify the proteins. The on-bead protein pull-down experiments were done in parallel with four samples (**P5-A, P5-B1, P5-B2** and **P5-C**) and were treated as four replicates and only those proteins with 2 or more unique peptides in at least three samples were selected for downstream analysis. The control experiment was used to eliminate the non-specifically bound proteins from the pool. This yielded a total of 442 proteins. The gel band analysis experiment was used to crosscheck and find the common proteins from the on-bead digestion experiment. The binding of the probe to LNCaP proteins was done in parallel to the probe binding in the presence of the competitor, P5. The expectation was that pull-down in the presence of cold P5 binding will lower the peptide count of specific binding proteins. 163 proteins were found to satisfy the criteria and among them, only 38 proteins were found to be common across both experiments, and were classified as the ‘P5 binding proteins’. The remainder of the 404 proteins from the on-bead digestion experiment were classified as ‘potential P5 binding proteins’. We did not analyze the 125 proteins from the in-gel digestion experiment.

##### MS data analysis for murine CD8^+^ T cells

IsobarQuant^46^ and Mascot (v2.2.07) were used to process the acquired data, which was searched against the Uniprot reference database of *Mus musculus* after the addition of common contaminants and reversed sequences. The following modifications were included into the search parameters: Carbamidomethyl (C) and TMT10 (K) (fixed modification), Acetyl (N-term), Oxidation (M) and TMT10 (N-term) (variable modifications). A mass error tolerance of 10 ppm was applied to full scan spectra and 0.02 Da to fragmentation spectra. Trypsin was selected as protease with an allowance of maximum two missed cleavages, a minimum peptide length of seven amino acids, and at least two unique peptides were required for protein identification. The false discovery rate (FDR) on peptide and protein level was set to 0.01.

The protein.txt output files from IsobarQuant were further processed using the R language. As quality filters, only proteins that were quantified with at least two unique peptides and have been identified in both biological replicates (analyzed in separate MS analysis) were used for further downstream analysis (416 proteins). The ‘signal_sum’ columns were used and potential batch-effects were removed using the respective function from the limma package^47^. Subsequently, the data was normalized using a variance stabilization normalization (vsn)^48^. The limma package was employed again to test for differential abundance between the various experimental conditions. T-values of the limma output were pasted into fdrtool^49^ in order to estimate false discovery rates (q-values were used as FDR).

Hierarchial clustering of all the peptides from five different samples was done using cluster 3.0^50^ and the results viewed with java treeview 3.0^51^. For functional annotation DAVID 6.8 software^52–54^ was used. For analysis we chose only those characters with P value > 0.05.

#### Microscopy

About 20000 LNCaP cells were grown on 1.5 coverslips in 24 well plates in complete RPMI 1640 medium with 10% FBS. After 48 hours the medium was aspirated and the cells were washed twice with PBS. 500 µL of phenol free RPMI 1640 supplemented with 10% charcoal stripped FBS was added to each well and incubated for an hour. 1 µM of **P5-C** was added to each well and incubated in the dark for 2 hours. Then media was removed from the wells and washed with cold PBS (special care was taken to avoid too much light exposure). 200 µL of cold PBS was added to each well and irradiated at 365 nm for 30 minutes at 4°C. After crosslinking PBS was removed and the cells were fixed in 0.5 mL of cold MeOH for 15 minutes at −20°C. After 6 one minute extraction with (10:55:0.75) chloroform:methanol:acetic acid the cells were clicked with 0.1 µM Alexa Fluor 488 azide (A10266, Thermofisher) in 200 µL PBS with freshly prepared 1mM TCEP, 100 µM TBTA and 1mM CuSO_4_ for 1 hour at RT with shaking. The coverslips were washed thrice in wells with PBS followed by two washes with water. Another parallel experiment with similar treatments but no probe was carried out as a control. The coverslips were mounted on slides with Prolong diamond anti-fade mountant (P36965, Thermofisher) with DAPI and visualized under Leica DM50000B fluorescent microscope equipped with narrow band-pass filters for DAPI-FITC with an ORCA-03G CCD camera (Hamamatsu). Digital images were captured using SmartCapture (Digital Scientific, UK).

## Acknowledgements

We thank Mathew Garnett for providing the LNCaP and U87MG cell lines; Fengtang Yang and the molecular cytogenetics group for their technical help with fluorescence microscopy; Jyoti Choudhary and Lu Yu, formerly of the proteomics mass spectrometry group of the Wellcome Sanger Institute. We acknowledge the EMBL Proteomics Core Facility for expert help.

## Funding

The ERC consolidator grant (ThDEFINE, Project ID: 646794) supported this study. SR was supported by an EIPOD fellowship and Ashoka University Individual Research grant for research visits. BM was supported by a CRUK Cancer Immunology grant (Ref. 20193).

## Author contribution

SR and SAT designed the experiments. SVL, SAT, SR and JS designed the experiments related to chemical synthesis of the probes. JS helped SR synthesize the P5 probes and JS did the analysis of the NMR, IR and figures of the synthesized probes. SR performed the experiments. BM and JP designed and performed the Th2 and immunoglobulin class switching experiments, helped SR culturing LNCaP cells occasionally. SR wrote the manuscript with help from BM. MLH did the on-bead digestion and mass spectrometry of CD8^+^ T cells. SAT, SVL and A-CG supervised the study. All authors commented on and approved the draft manuscript before submission.

## Competing financial interests

The authors declare no competing financial interests.

## Additional Information

Supplementary information and chemical compound information.

