## Supplementary figures for "CLICK-enabled analogues reveal pregnenolone interactomes in cancer and immune cells"

### Slide 1
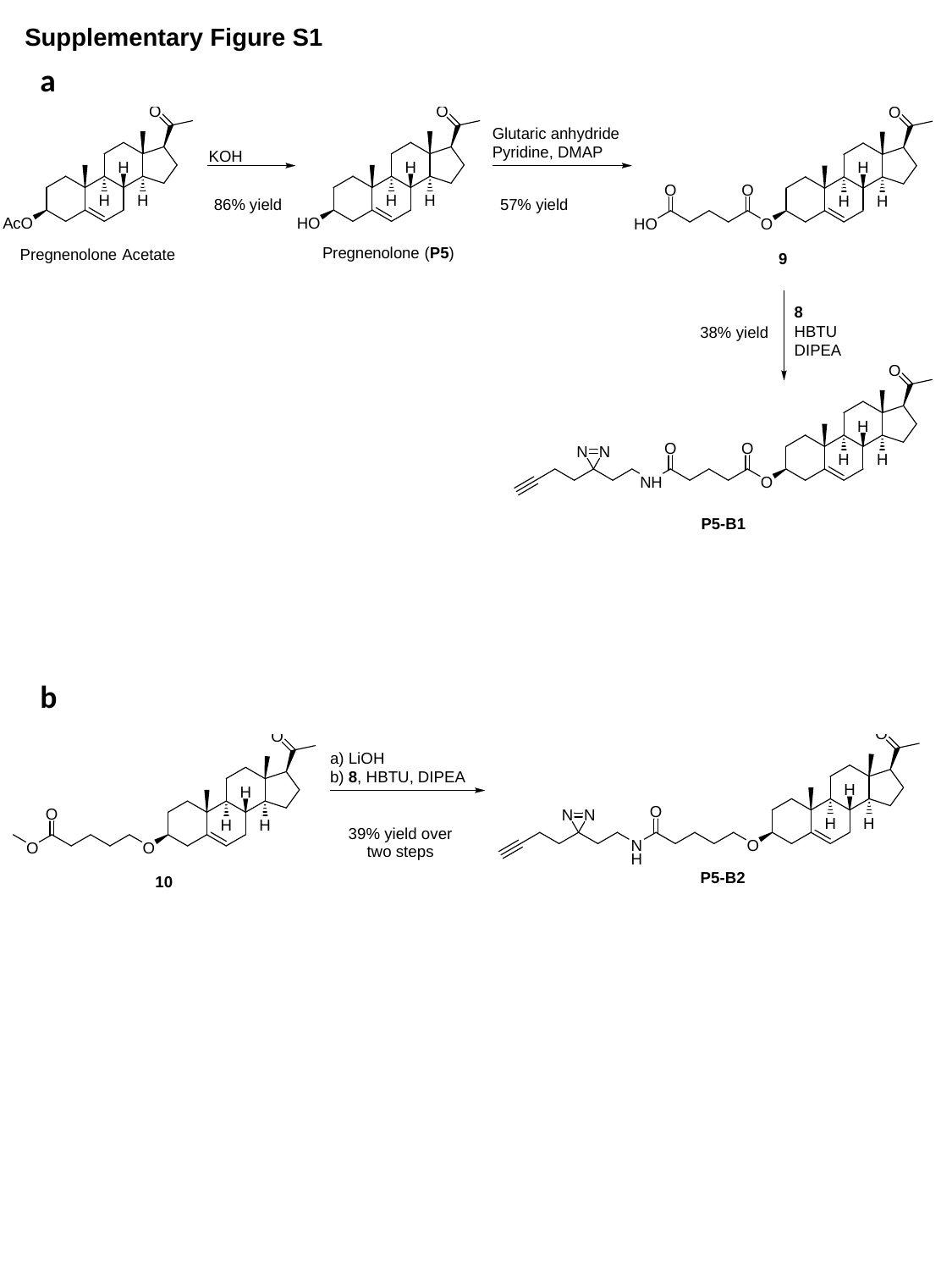

Supplementary Figure S1
a
b

### Slide 2
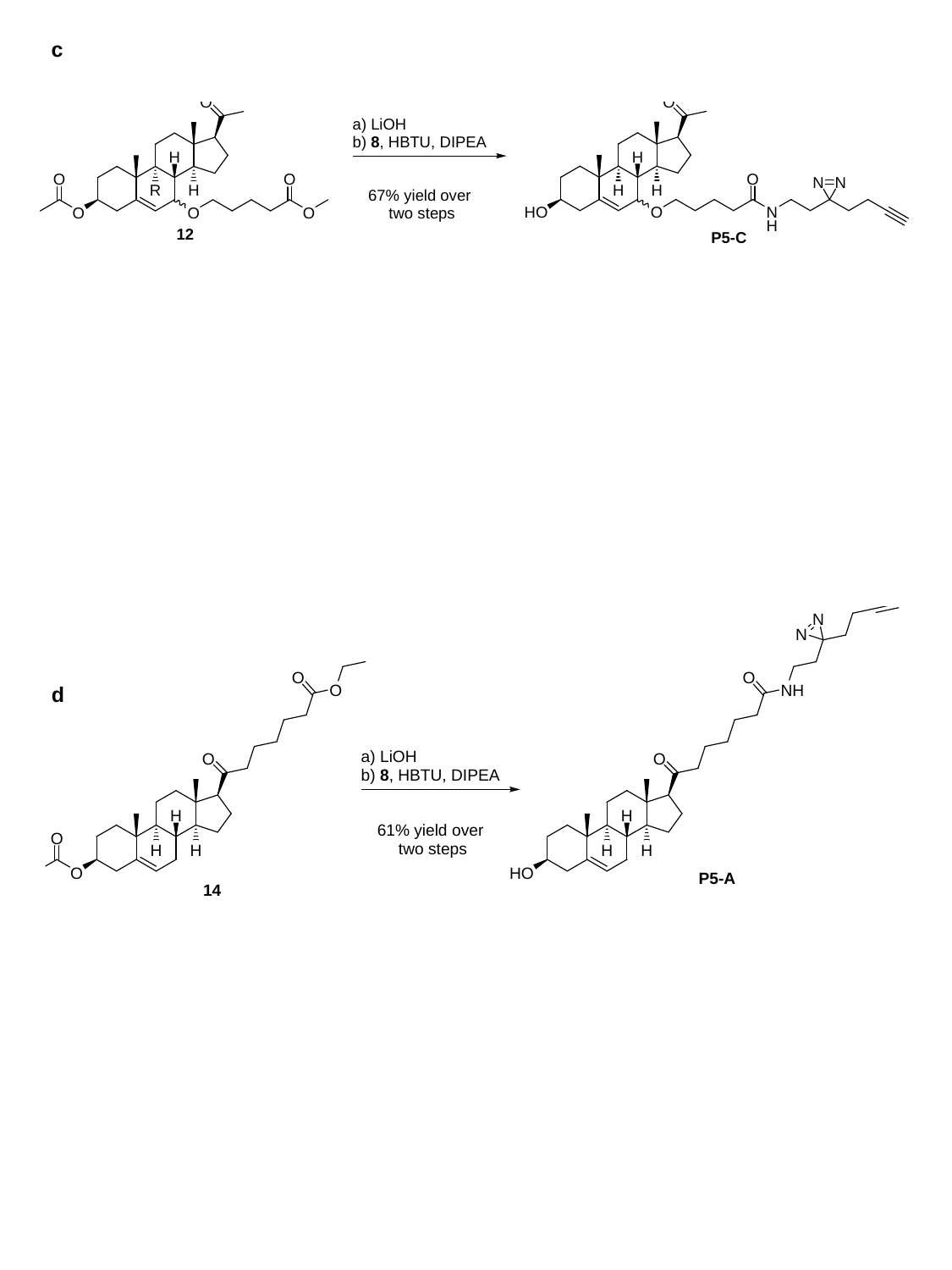

c
d

### Slide 3
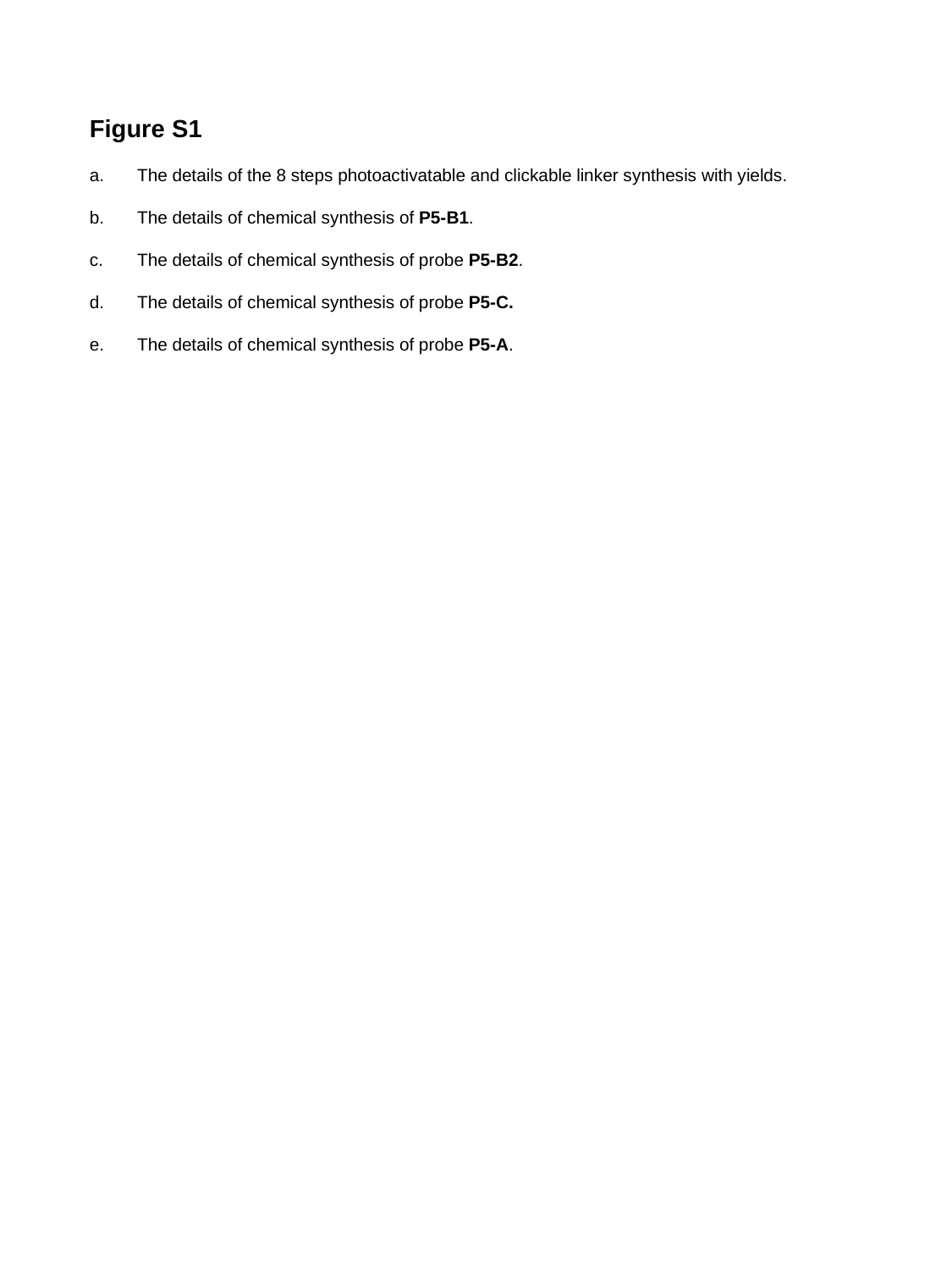

Figure S1
The details of the 8 steps photoactivatable and clickable linker synthesis with yields.
The details of chemical synthesis of P5-B1.
The details of chemical synthesis of probe P5-B2.
The details of chemical synthesis of probe P5-C.
The details of chemical synthesis of probe P5-A.

### Slide 4
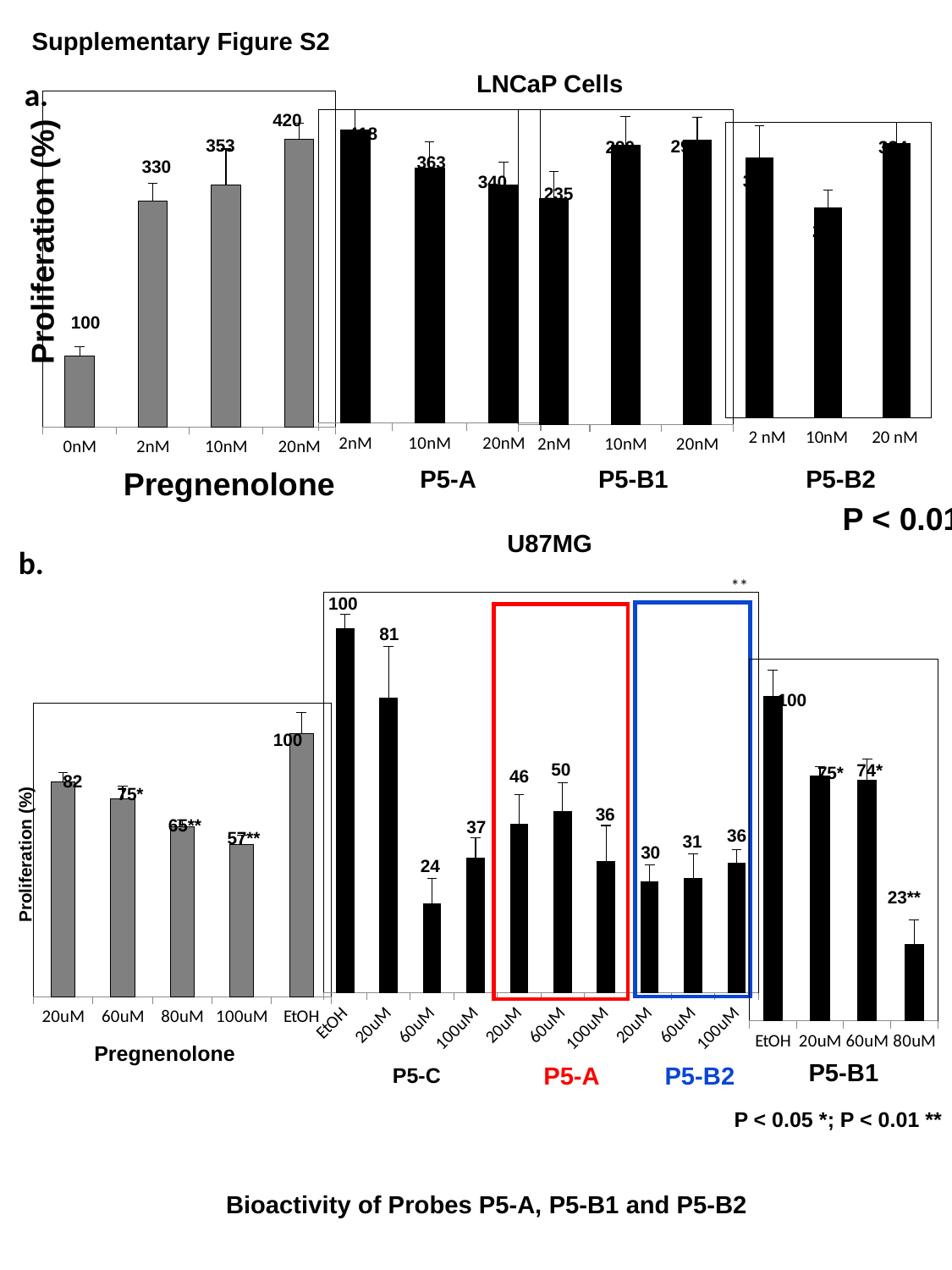

Supplementary Figure S2
LNCaP Cells
a.
#### Chart
| Category | |
|---|---|
| 0nM | 0.025375000000000012 |
| 2nM | 0.08062500000000056 |
| 10nM | 0.08650000000000001 |
| 20nM | 0.10274999999999998 |420
#### Chart
| Category | |
|---|---|
| 2nM | 0.42137500000000094 |
| 10nM | 0.36587500000000106 |
| 20nM | 0.3418750000000003 |
#### Chart
| Category | |
|---|---|
| 2nM | 0.05750000000000002 |
| 10nM | 0.071 |
| 20nM | 0.07225 |295
290
235
#### Chart
| Category | |
|---|---|
| 2mM | 0.079125 |
| 10mM | 0.063875 |
| 20mM | 0.08350000000000027 |334
317
256
418
353
363
330
340
Proliferation (%)
100
2 nM 10nM 20 nM
P5-A
P5-B1
P5-B2
Pregnenolone
P < 0.01
U87MG
b.
#### Chart
| Category | |
|---|---|
| EtOH | 0.45542857142857224 |
| 20uM | 0.36842857142857277 |
| 60uM | 0.11085714285714295 |
| 100uM | 0.16828571428571387 |
| 20uM | 0.210142857142857 |
| 60uM | 0.226142857142857 |
| 100uM | 0.164142857142857 |
| 20uM | 0.1383333333333335 |
| 60uM | 0.142666666666667 |
| 100uM | 0.16228571428571387 |**
100
81
#### Chart
| Category | |
|---|---|
| EtOH | 0.6277500000000024 |
| 20uM | 0.4731428571428584 |
| 60uM | 0.4660000000000001 |
| 80uM | 0.14750000000000021 |100
#### Chart
| Category | |
|---|---|
| 20uM | 0.512 |
| 60uM | 0.47075 |
| 80uM | 0.4058750000000003 |
| 100uM | 0.36175 |
| EtOH | 0.6277500000000024 |100
50
74*
75*
46
82
75*
36
65**
37
36
57**
31
30
Proliferation (%)
24
23**
Pregnenolone
P5-B1
P5-B2
P5-A
P5-C
P < 0.05 *; P < 0.01 **
Bioactivity of Probes P5-A, P5-B1 and P5-B2

### Slide 5
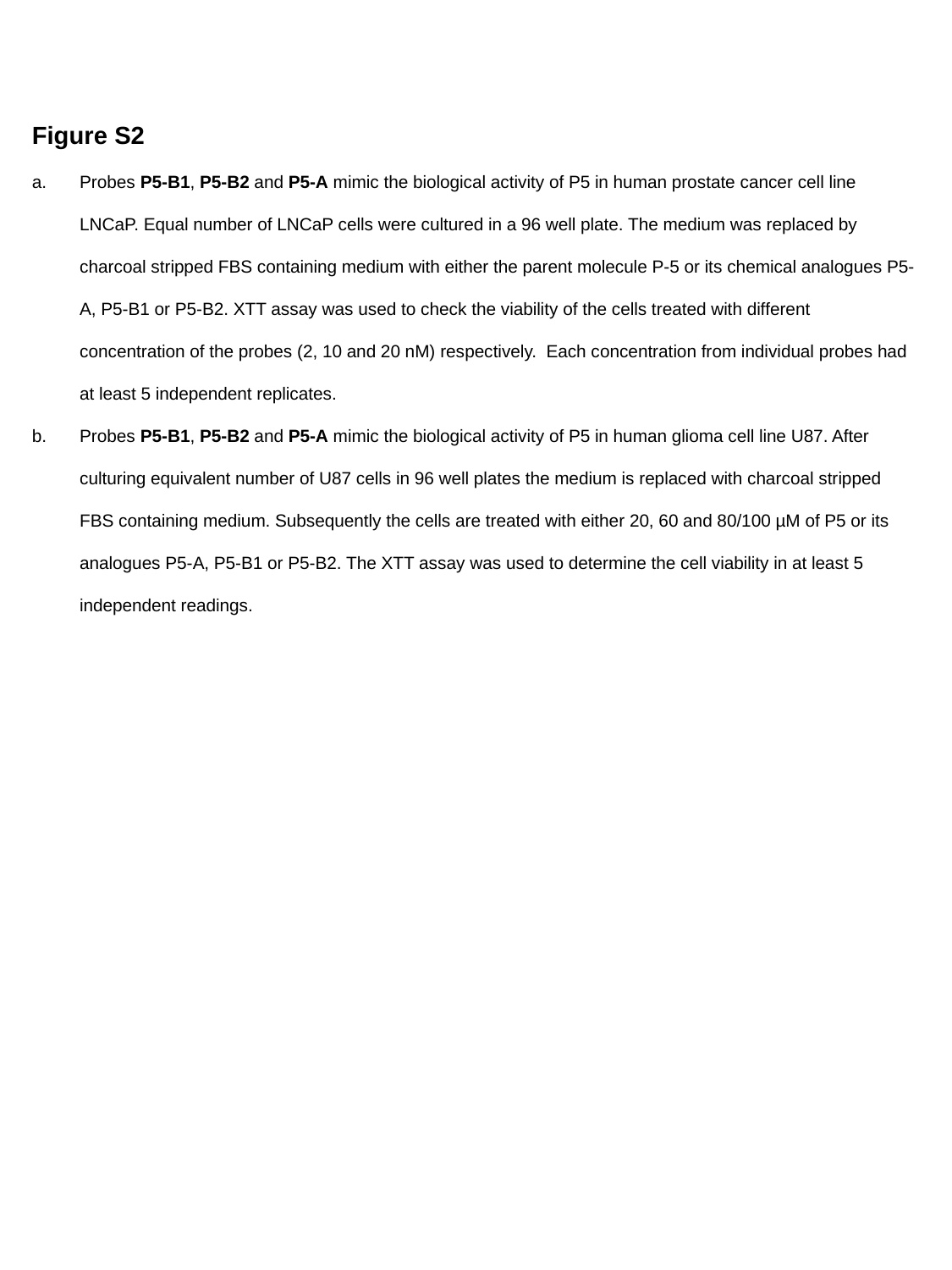

Figure S2
Probes P5-B1, P5-B2 and P5-A mimic the biological activity of P5 in human prostate cancer cell line LNCaP. Equal number of LNCaP cells were cultured in a 96 well plate. The medium was replaced by charcoal stripped FBS containing medium with either the parent molecule P-5 or its chemical analogues P5-A, P5-B1 or P5-B2. XTT assay was used to check the viability of the cells treated with different concentration of the probes (2, 10 and 20 nM) respectively. Each concentration from individual probes had at least 5 independent replicates.
Probes P5-B1, P5-B2 and P5-A mimic the biological activity of P5 in human glioma cell line U87. After culturing equivalent number of U87 cells in 96 well plates the medium is replaced with charcoal stripped FBS containing medium. Subsequently the cells are treated with either 20, 60 and 80/100 µM of P5 or its analogues P5-A, P5-B1 or P5-B2. The XTT assay was used to determine the cell viability in at least 5 independent readings.

### Slide 6
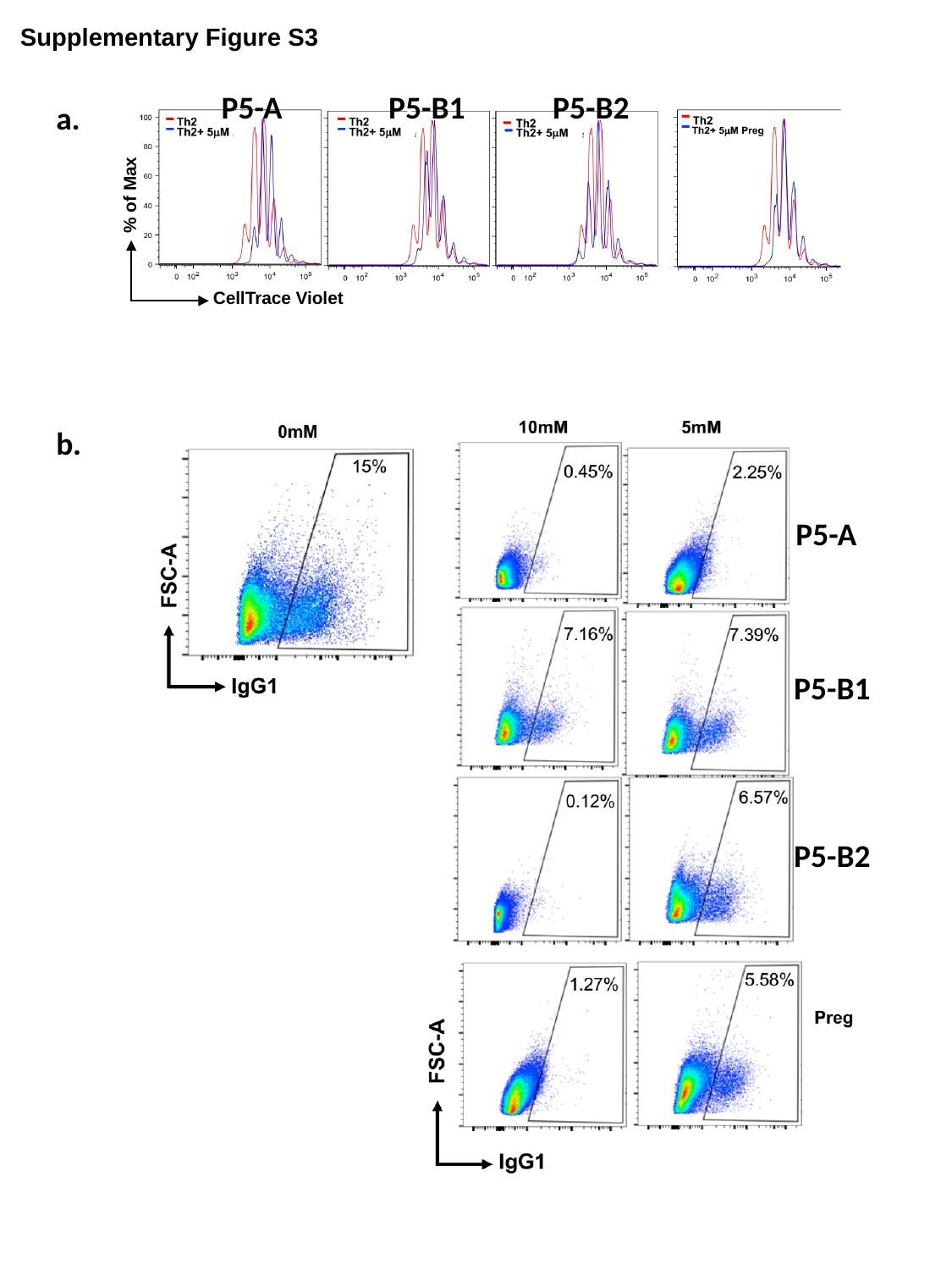

Supplementary Figure S3
P5-B1
P5-B2
P5-A
a.
% of Max
CellTrace Violet
b.
P5-A
P5-B1
P5-B2

### Slide 7
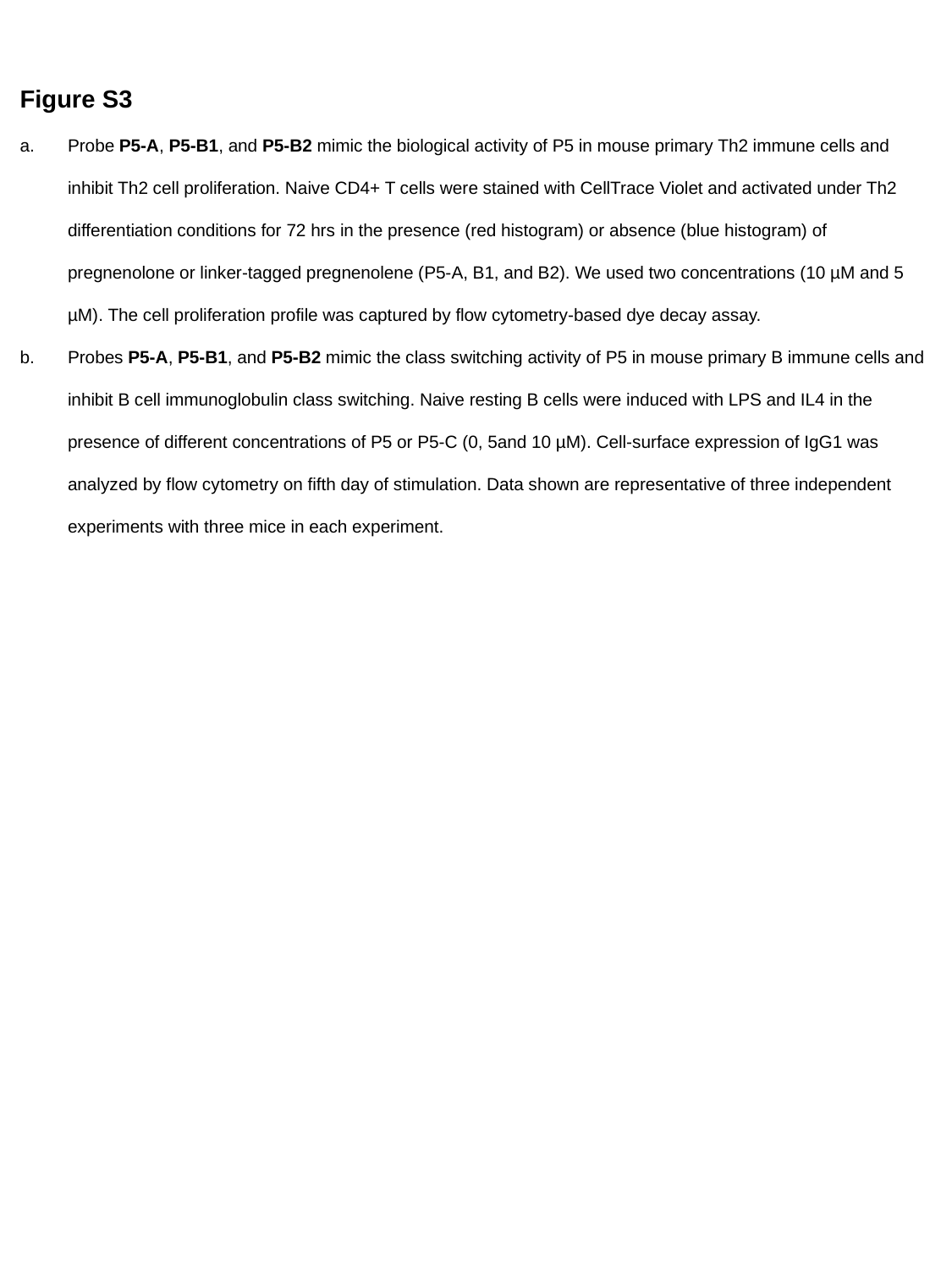

Figure S3
Probe P5-A, P5-B1, and P5-B2 mimic the biological activity of P5 in mouse primary Th2 immune cells and inhibit Th2 cell proliferation. Naive CD4+ T cells were stained with CellTrace Violet and activated under Th2 differentiation conditions for 72 hrs in the presence (red histogram) or absence (blue histogram) of pregnenolone or linker-tagged pregnenolene (P5-A, B1, and B2). We used two concentrations (10 µM and 5 µM). The cell proliferation profile was captured by flow cytometry-based dye decay assay.
Probes P5-A, P5-B1, and P5-B2 mimic the class switching activity of P5 in mouse primary B immune cells and inhibit B cell immunoglobulin class switching. Naive resting B cells were induced with LPS and IL4 in the presence of different concentrations of P5 or P5-C (0, 5and 10 µM). Cell-surface expression of IgG1 was analyzed by flow cytometry on fifth day of stimulation. Data shown are representative of three independent experiments with three mice in each experiment.

### Slide 8
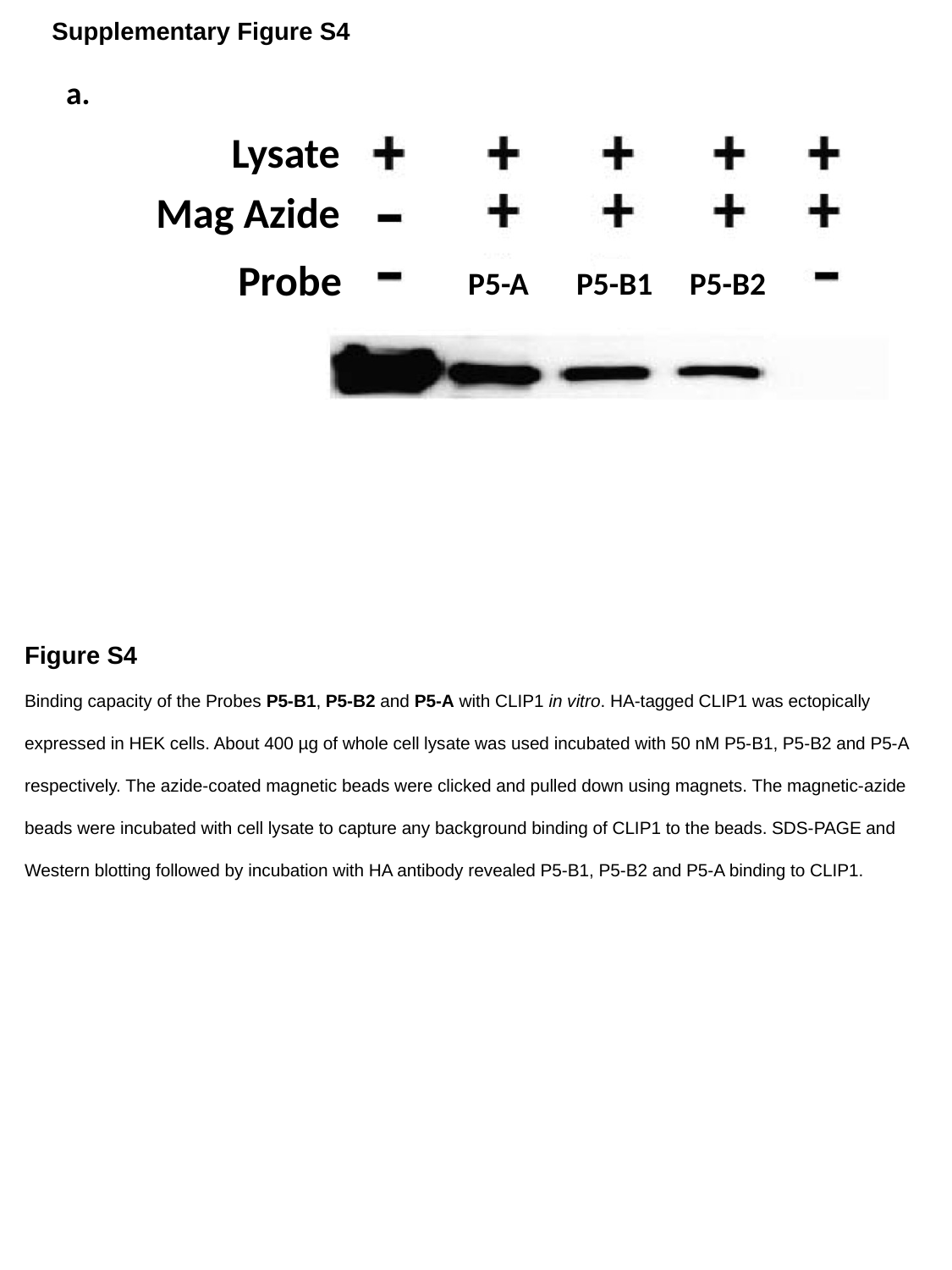

Supplementary Figure S4
a.
Lysate
Mag Azide
Probe
P5-B1
P5-A
P5-B2
Figure S4
Binding capacity of the Probes P5-B1, P5-B2 and P5-A with CLIP1 in vitro. HA-tagged CLIP1 was ectopically expressed in HEK cells. About 400 µg of whole cell lysate was used incubated with 50 nM P5-B1, P5-B2 and P5-A respectively. The azide-coated magnetic beads were clicked and pulled down using magnets. The magnetic-azide beads were incubated with cell lysate to capture any background binding of CLIP1 to the beads. SDS-PAGE and Western blotting followed by incubation with HA antibody revealed P5-B1, P5-B2 and P5-A binding to CLIP1.

### Slide 9
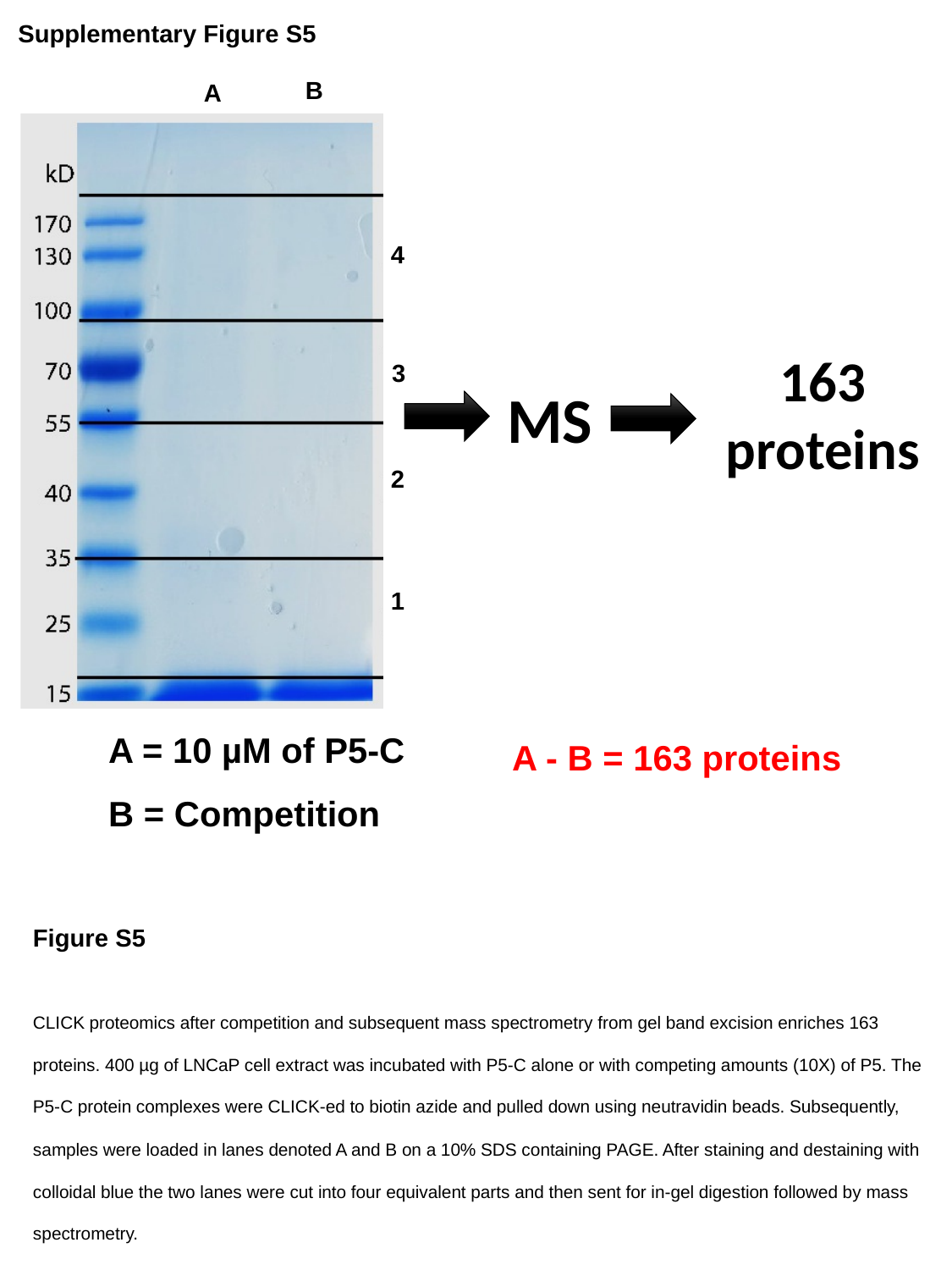

Supplementary Figure S5
B
A
4
163 proteins
3
MS
2
1
A = 10 µM of P5-C
B = Competition
A - B = 163 proteins
Figure S5
CLICK proteomics after competition and subsequent mass spectrometry from gel band excision enriches 163 proteins. 400 µg of LNCaP cell extract was incubated with P5-C alone or with competing amounts (10X) of P5. The P5-C protein complexes were CLICK-ed to biotin azide and pulled down using neutravidin beads. Subsequently, samples were loaded in lanes denoted A and B on a 10% SDS containing PAGE. After staining and destaining with colloidal blue the two lanes were cut into four equivalent parts and then sent for in-gel digestion followed by mass spectrometry.

### Slide 10
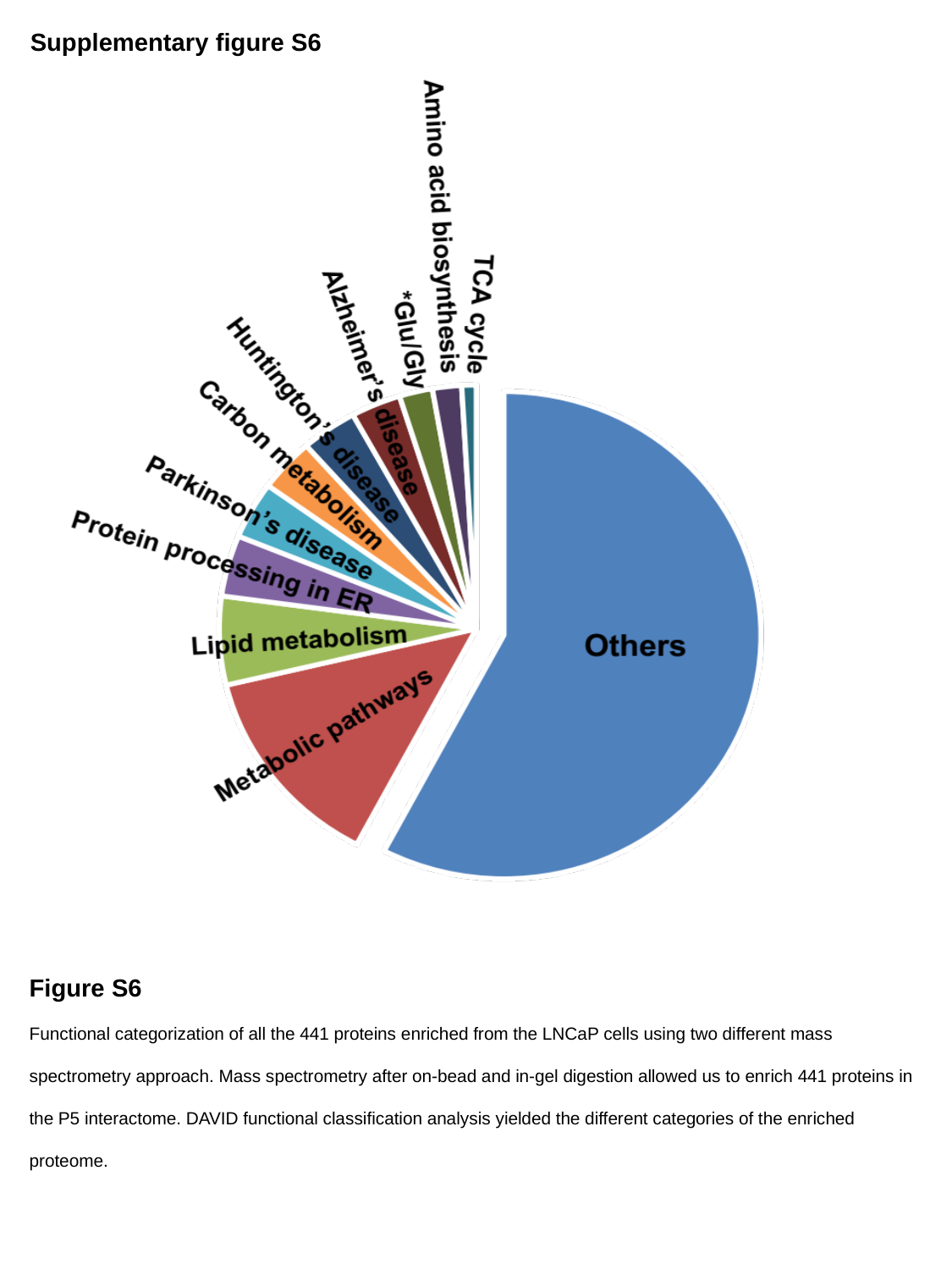

Supplementary figure S6
Non-Membranous
Figure S6
Functional categorization of all the 441 proteins enriched from the LNCaP cells using two different mass spectrometry approach. Mass spectrometry after on-bead and in-gel digestion allowed us to enrich 441 proteins in the P5 interactome. DAVID functional classification analysis yielded the different categories of the enriched proteome.

### Slide 11
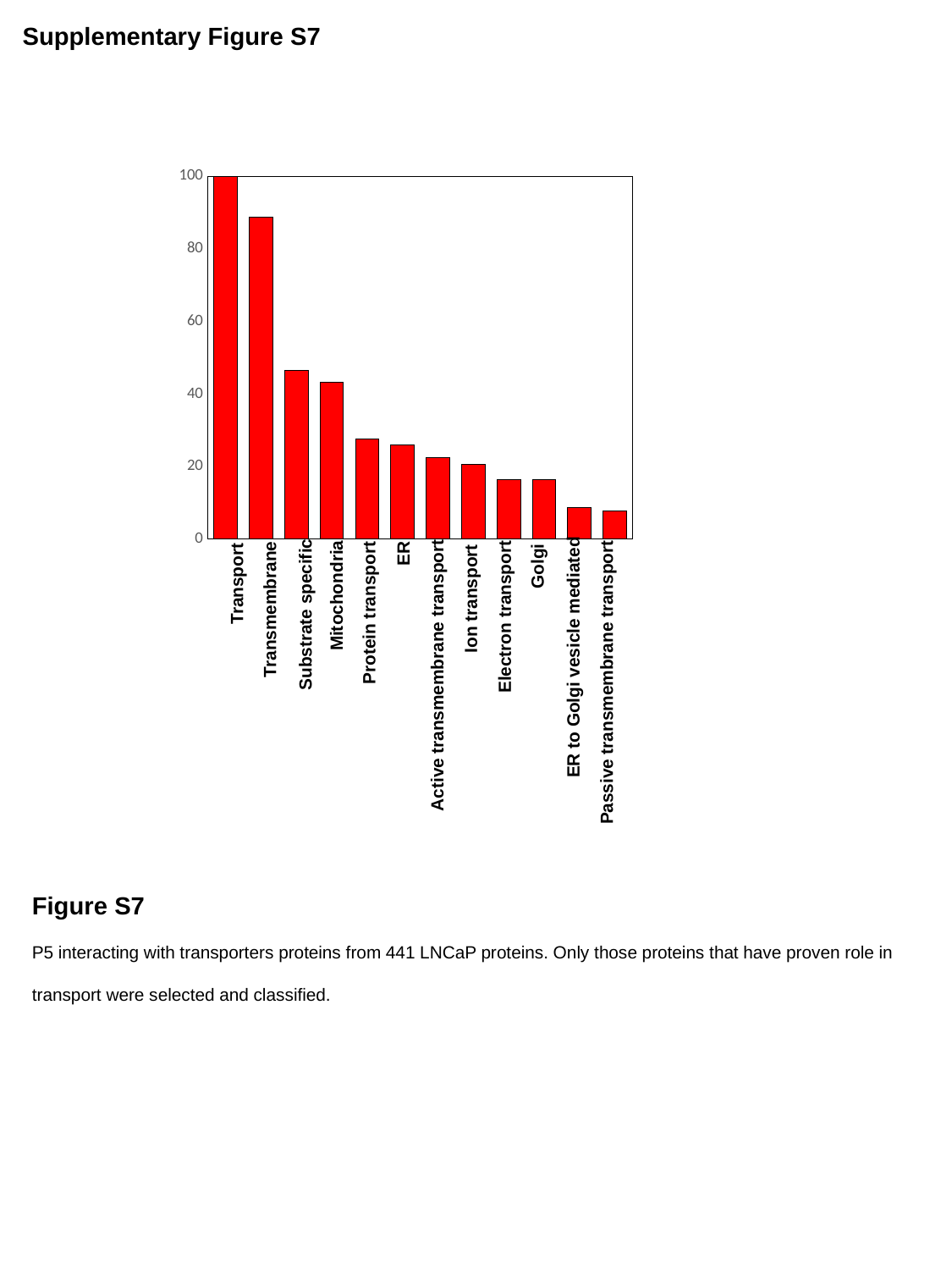

Supplementary Figure S7
#### Chart
| Category | |
|---|---|ER
Golgi
Transport
Mitochondria
Ion transport
Transmembrane
Protein transport
Substrate specific
Electron transport
ER to Golgi vesicle mediated
Active transmembrane transport
Passive transmembrane transport
Figure S7
P5 interacting with transporters proteins from 441 LNCaP proteins. Only those proteins that have proven role in transport were selected and classified.

### Slide 12
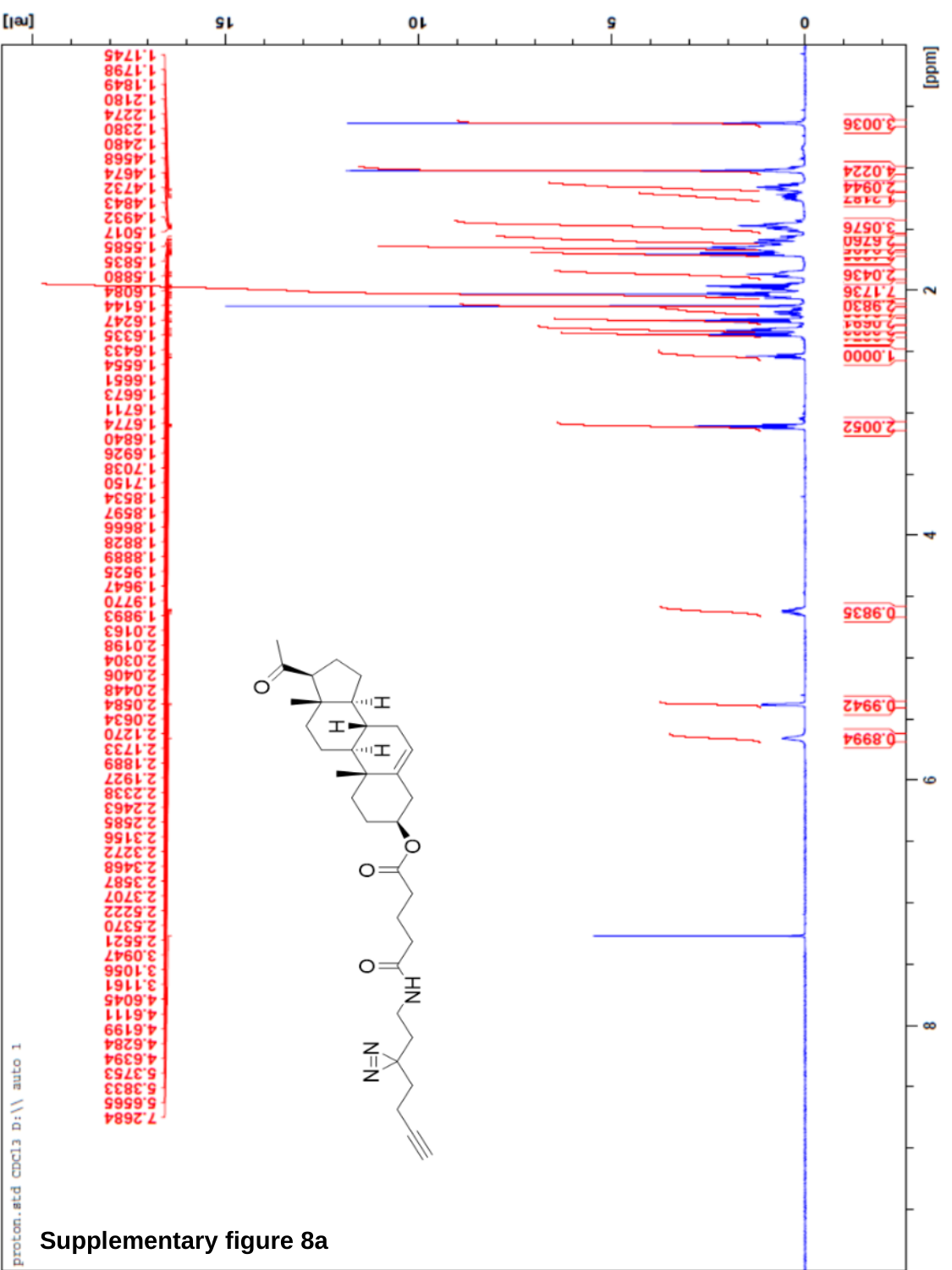

Supplementary figure 8a

### Slide 13
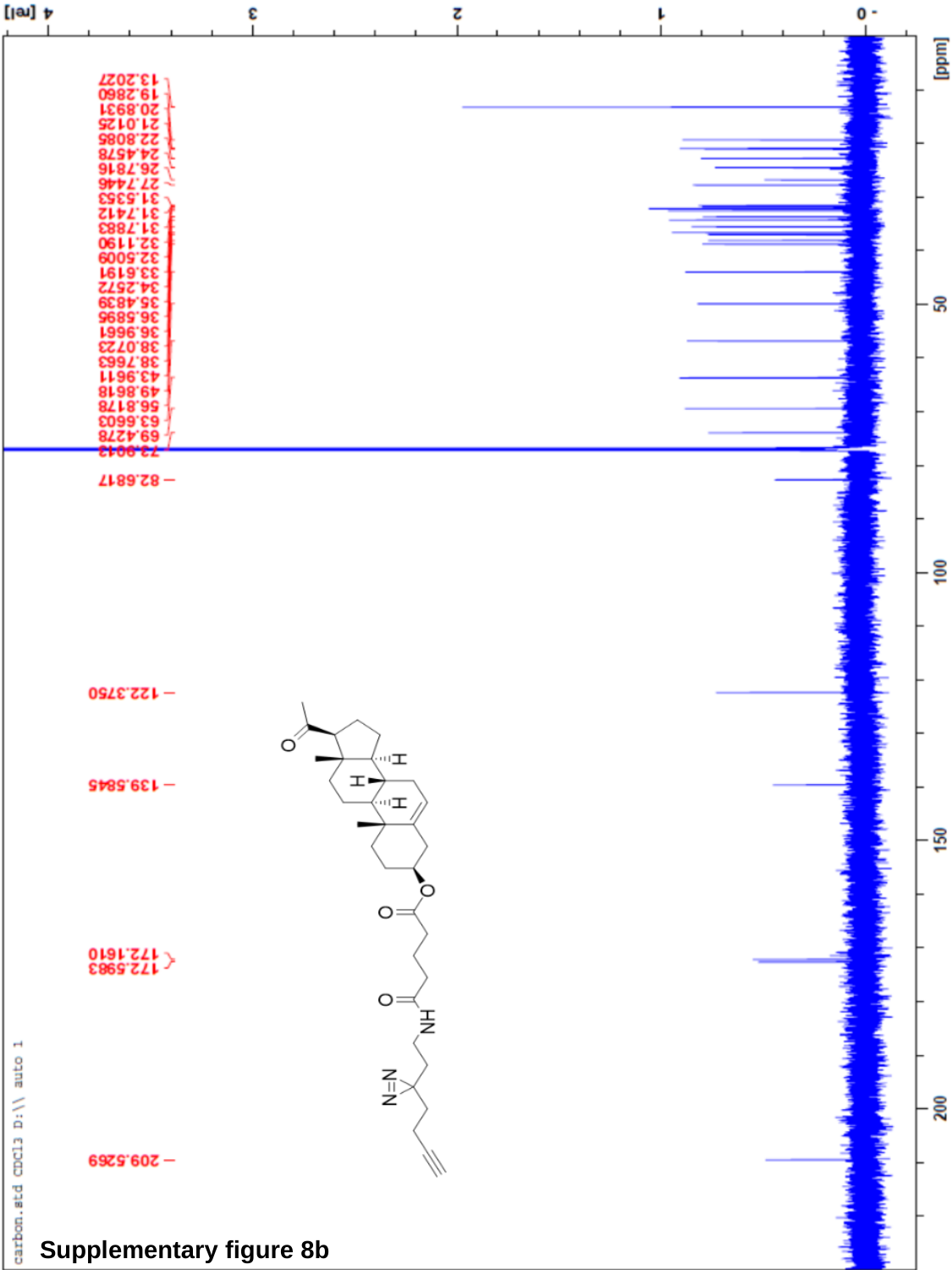

Supplementary figure 8b

### Slide 14
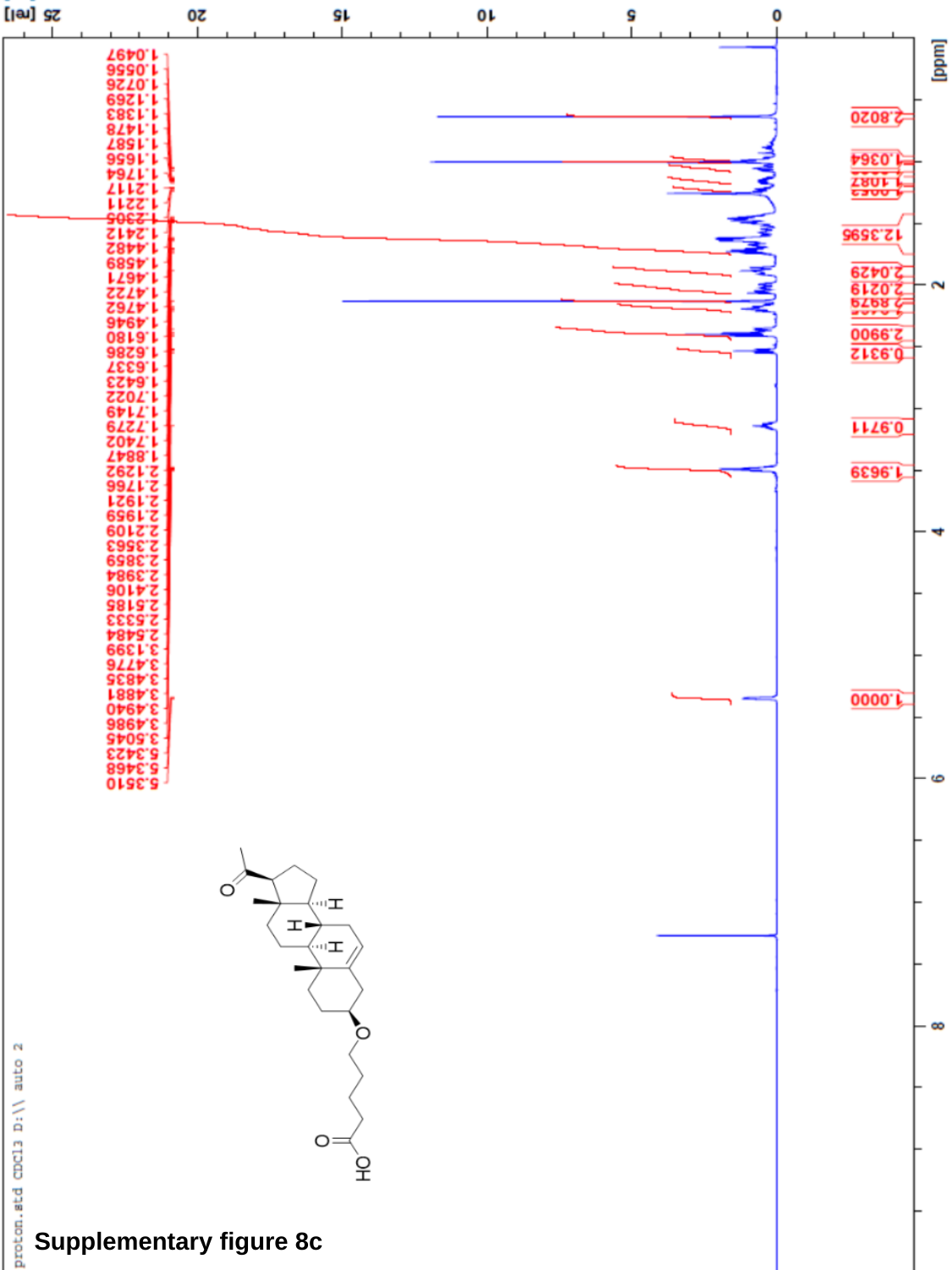

Supplementary figure 8c

### Slide 15
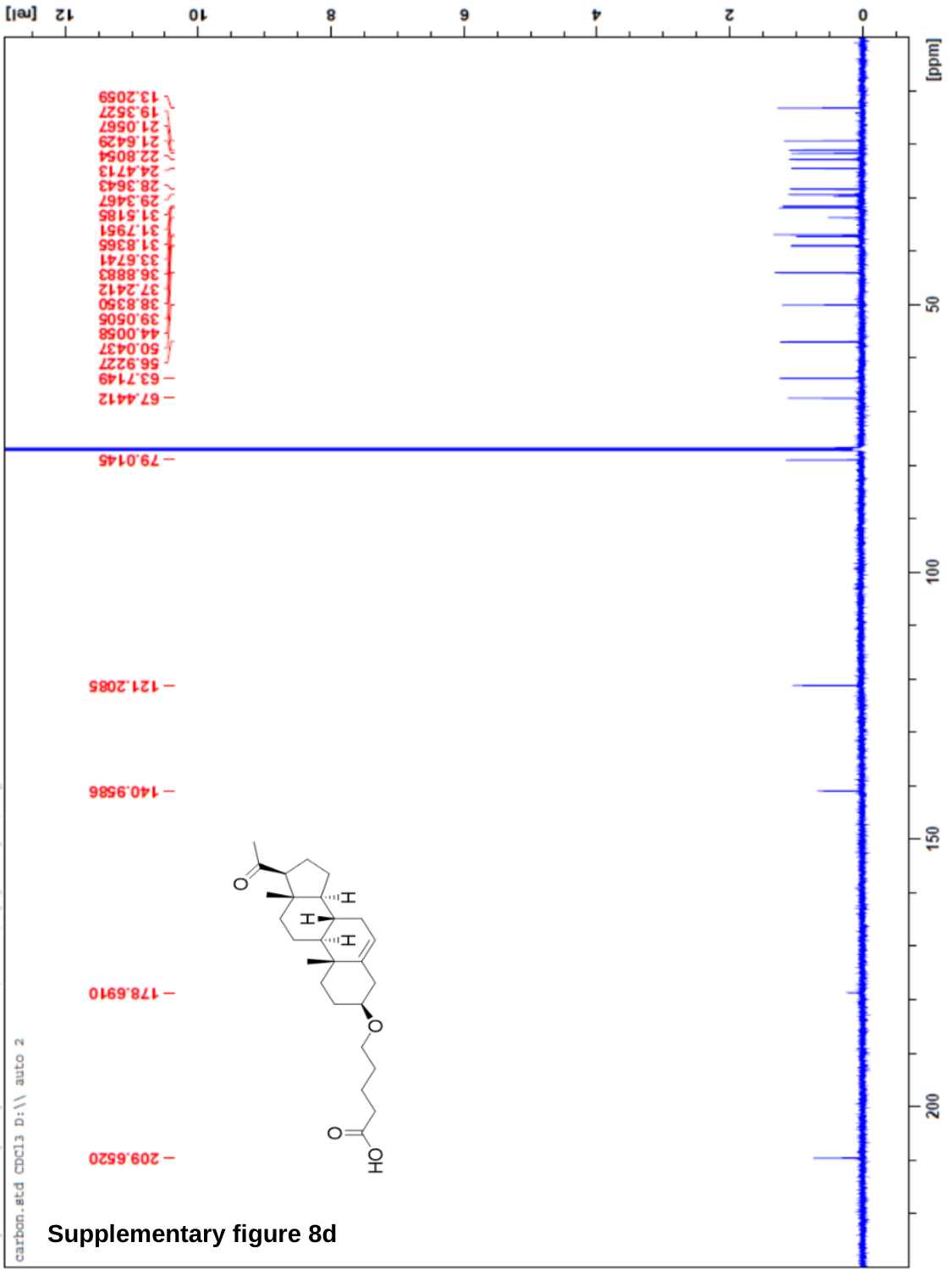

Supplementary figure 8d

### Slide 16
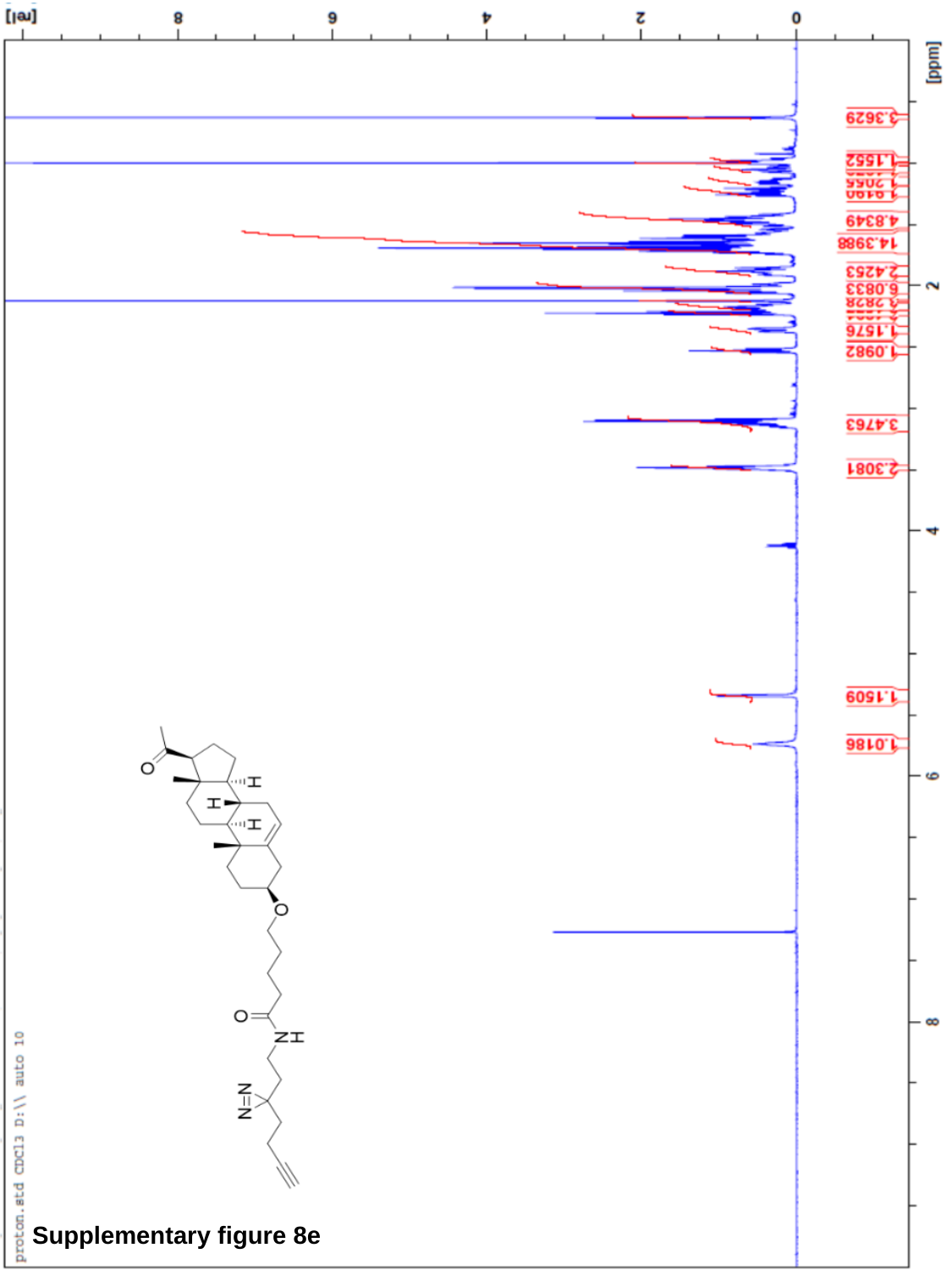

Supplementary figure 8e

### Slide 17
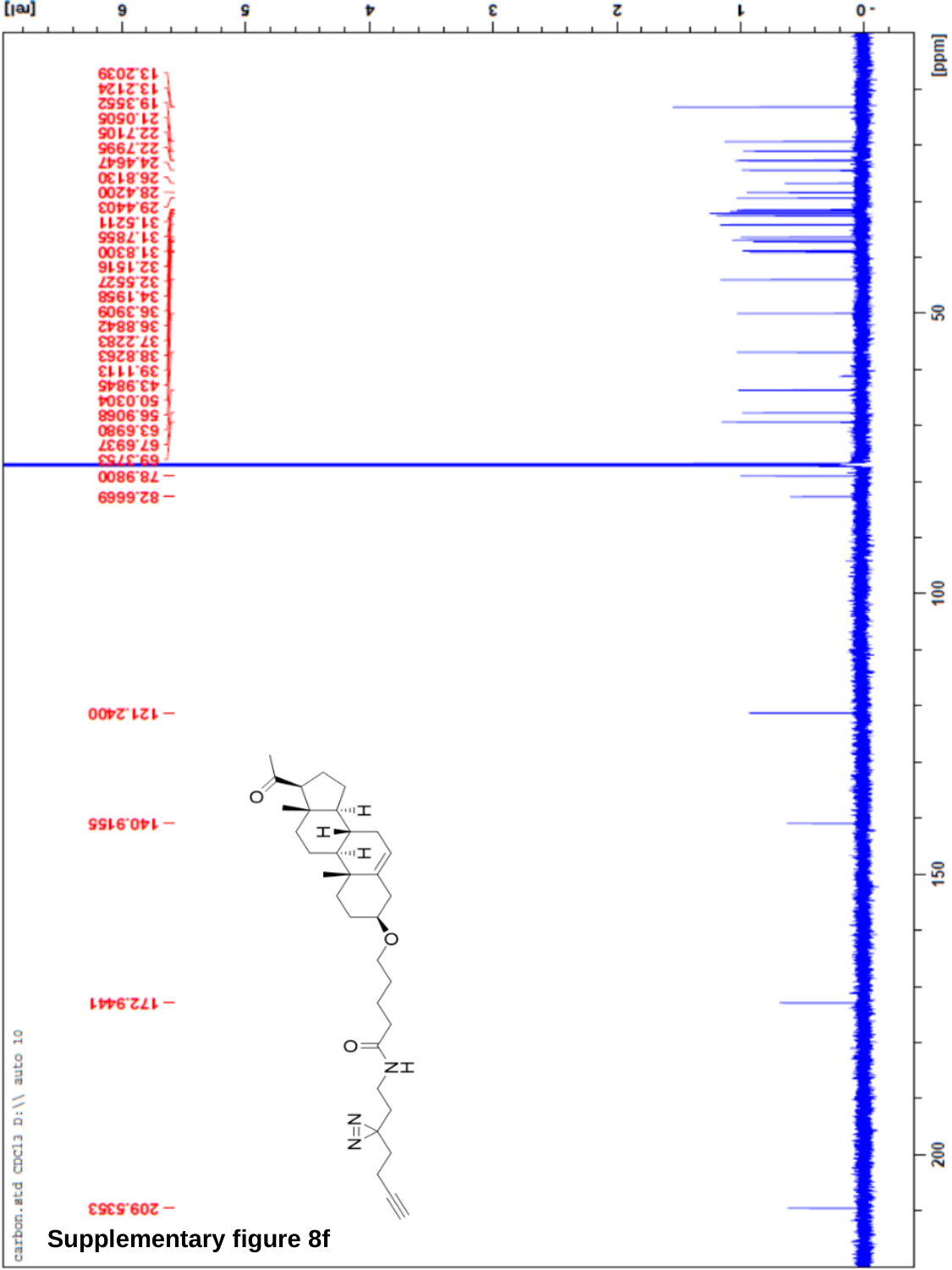

Supplementary figure 8f

### Slide 18
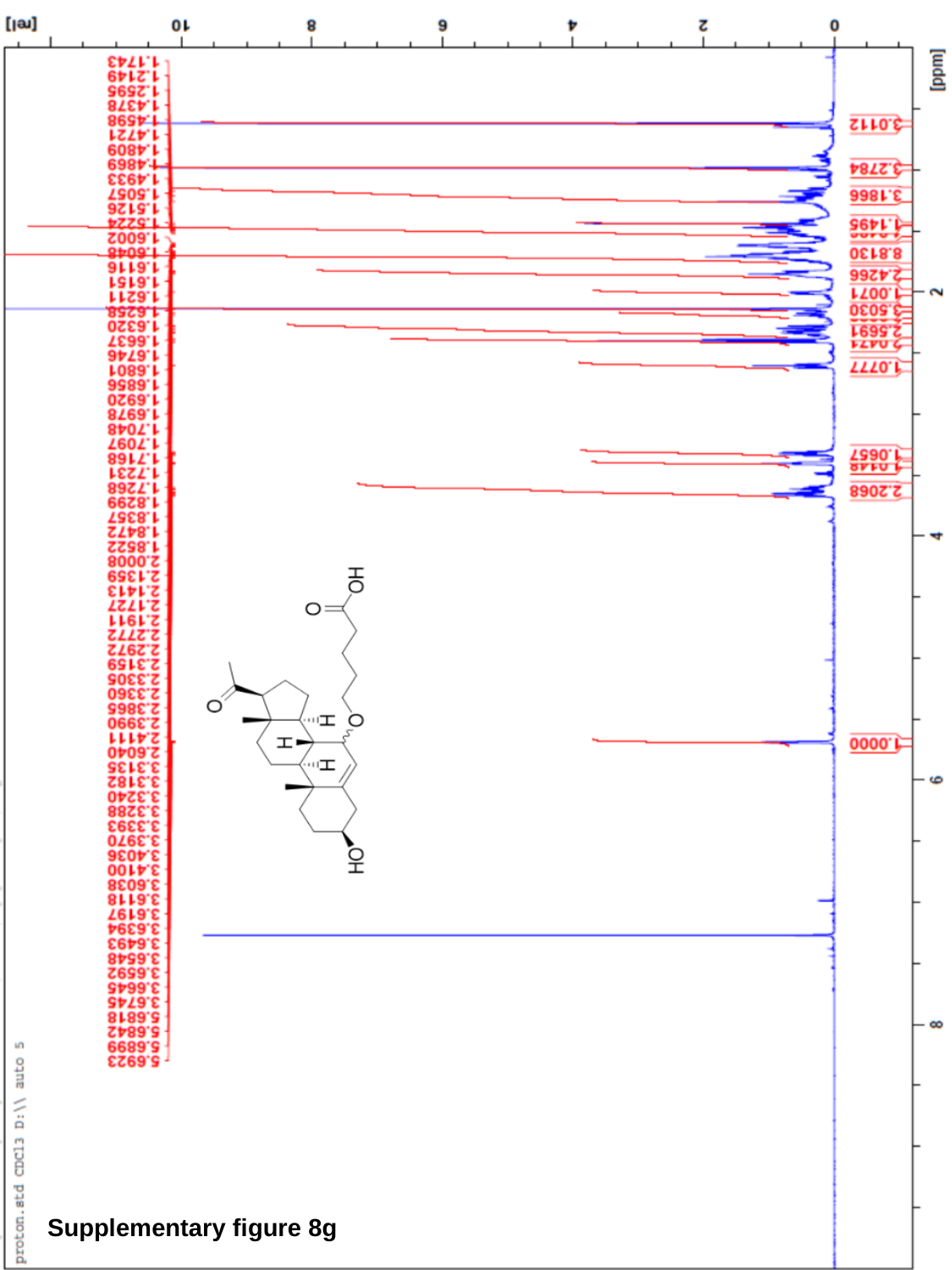

Supplementary figure 8g

### Slide 19
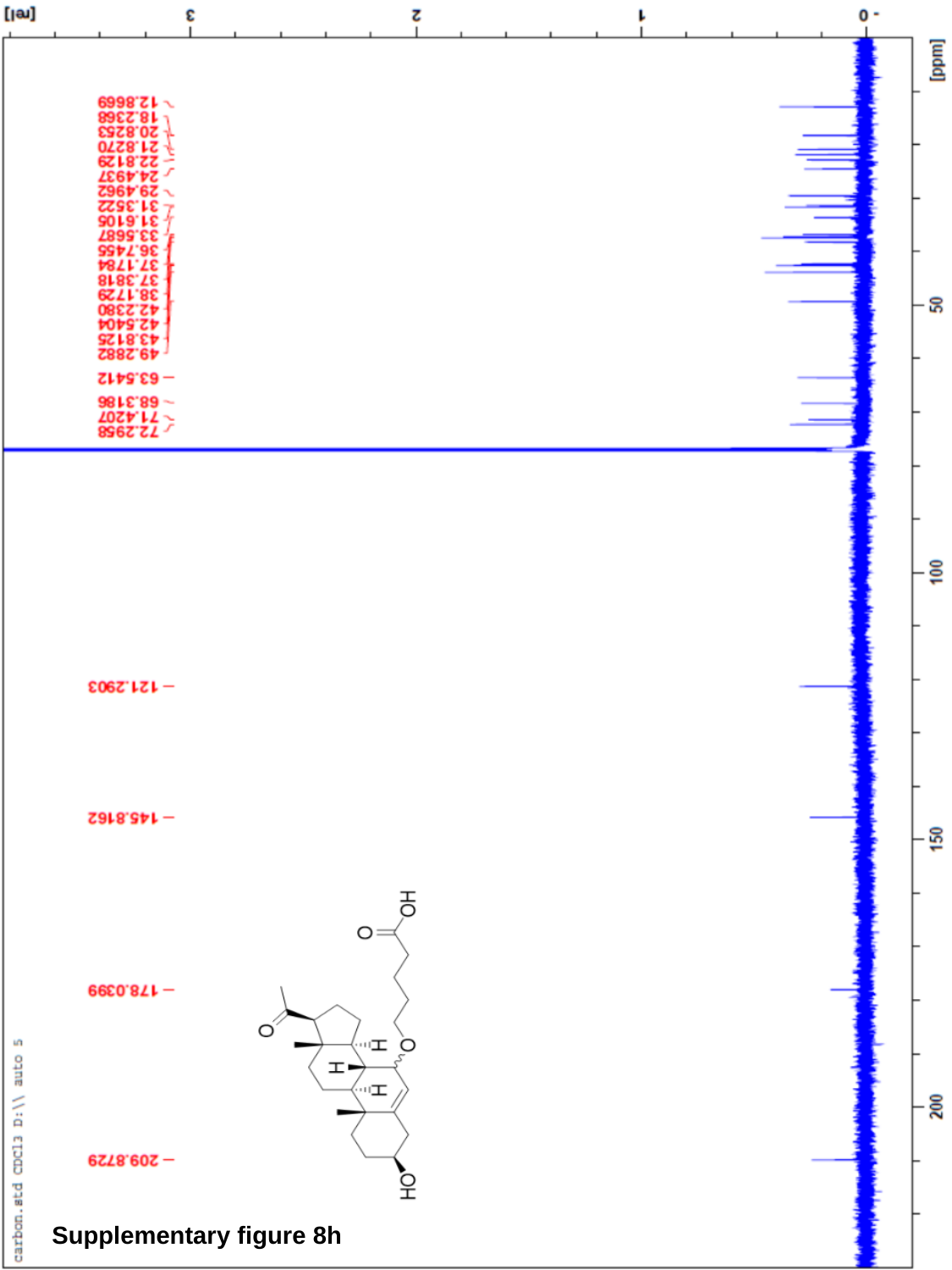

Supplementary figure 8h

### Slide 20
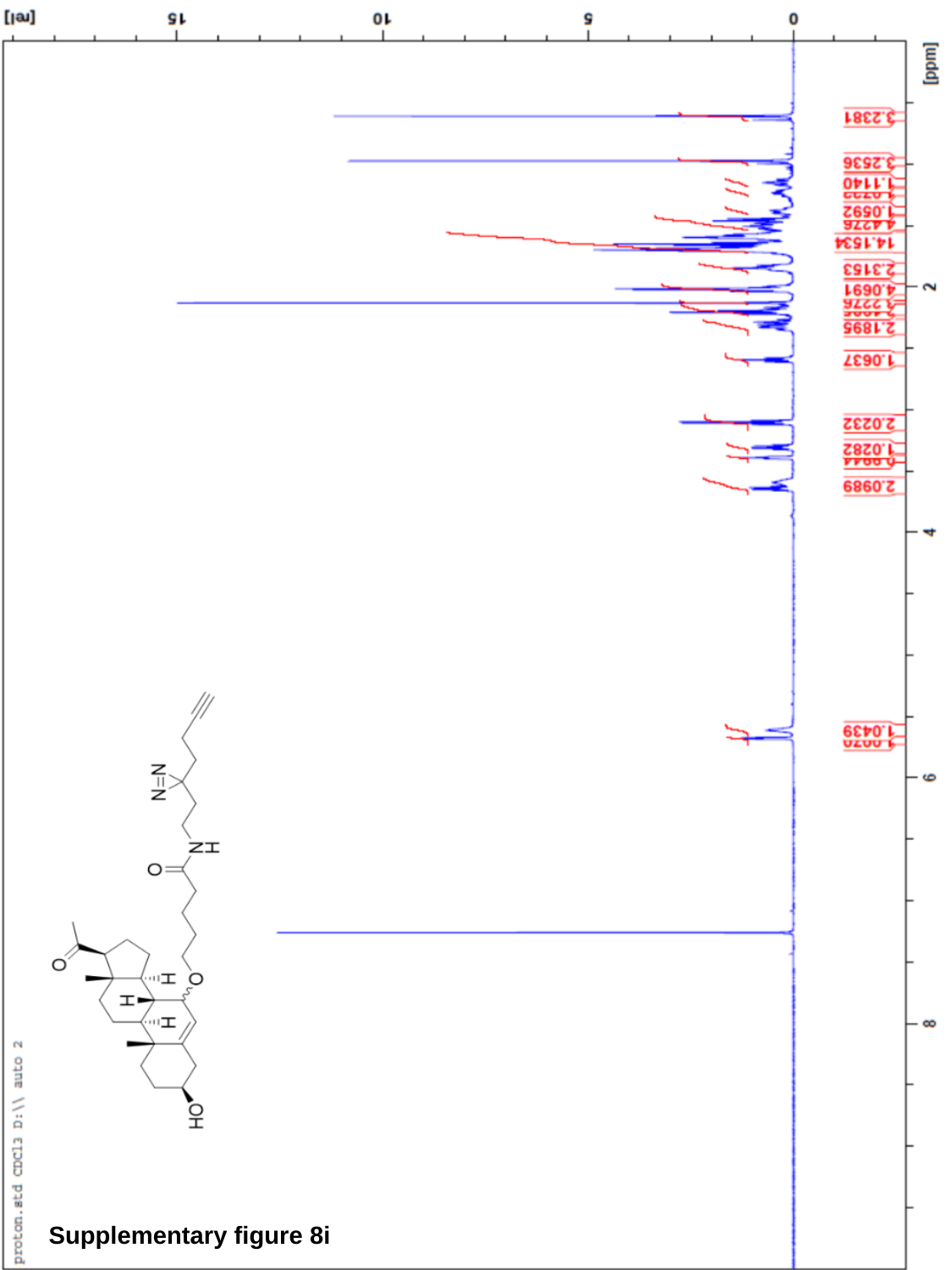

Supplementary figure 8i

### Slide 21
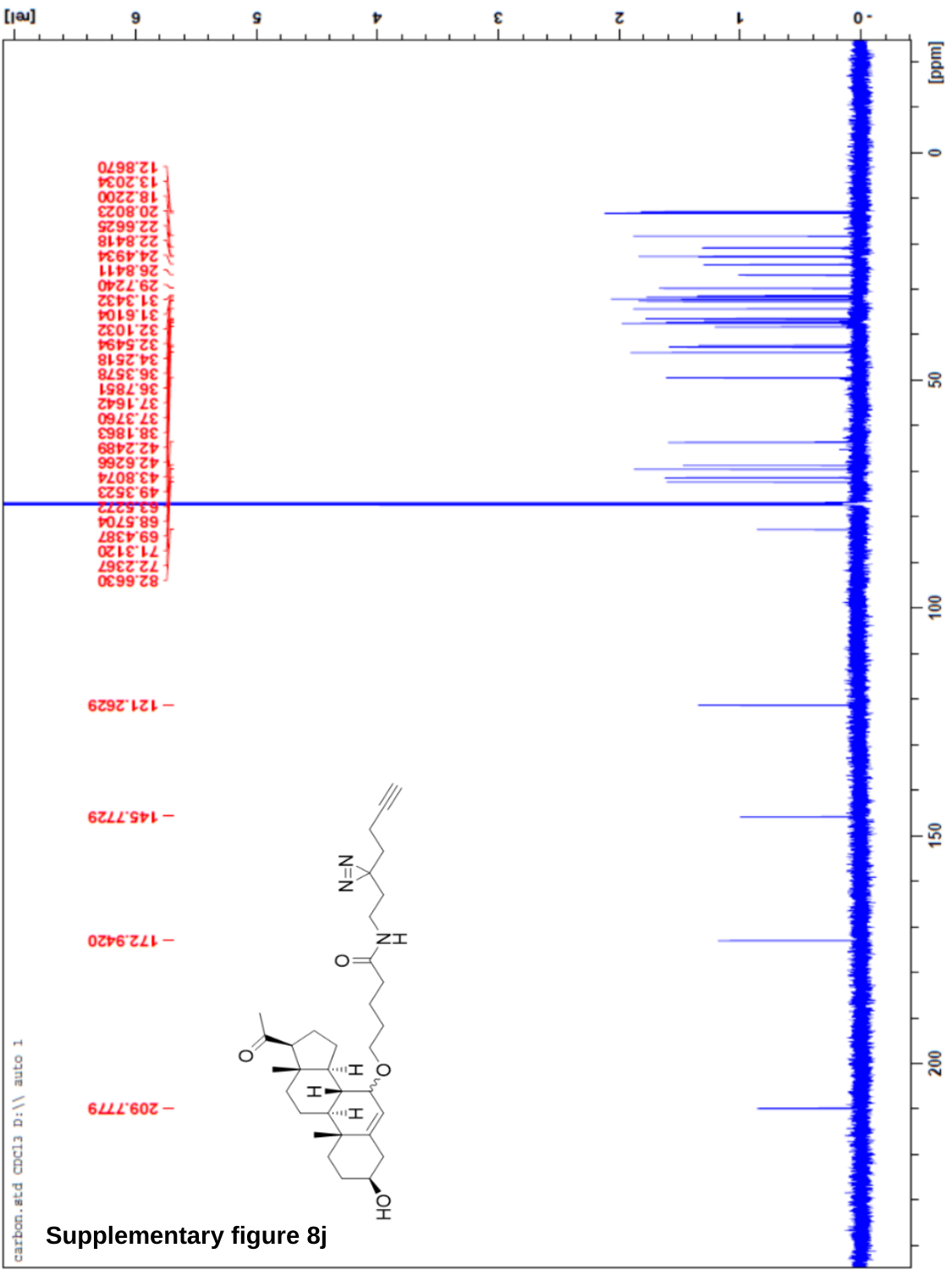

Supplementary figure 8j

### Slide 22
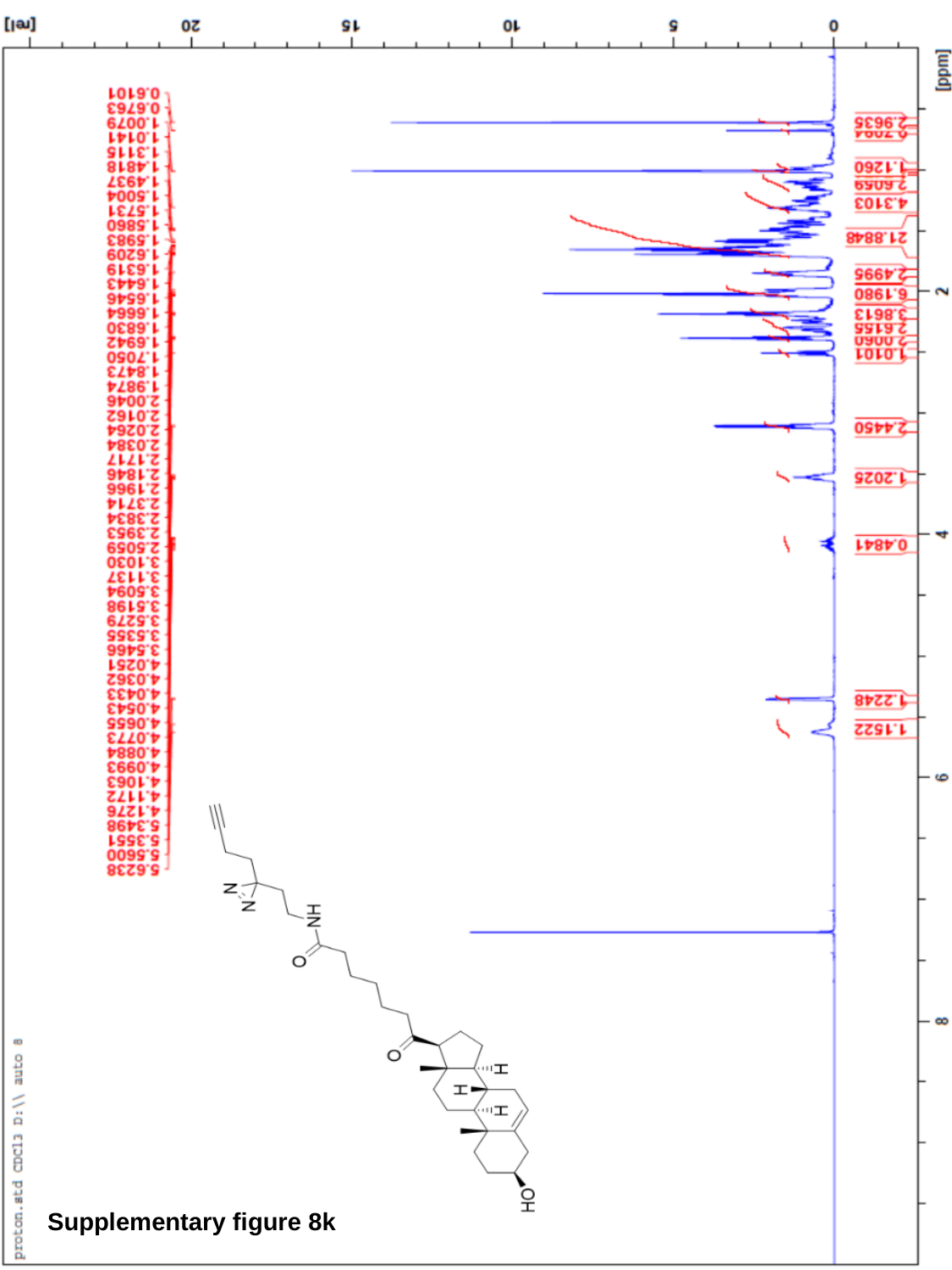

Supplementary figure 8k

### Slide 23
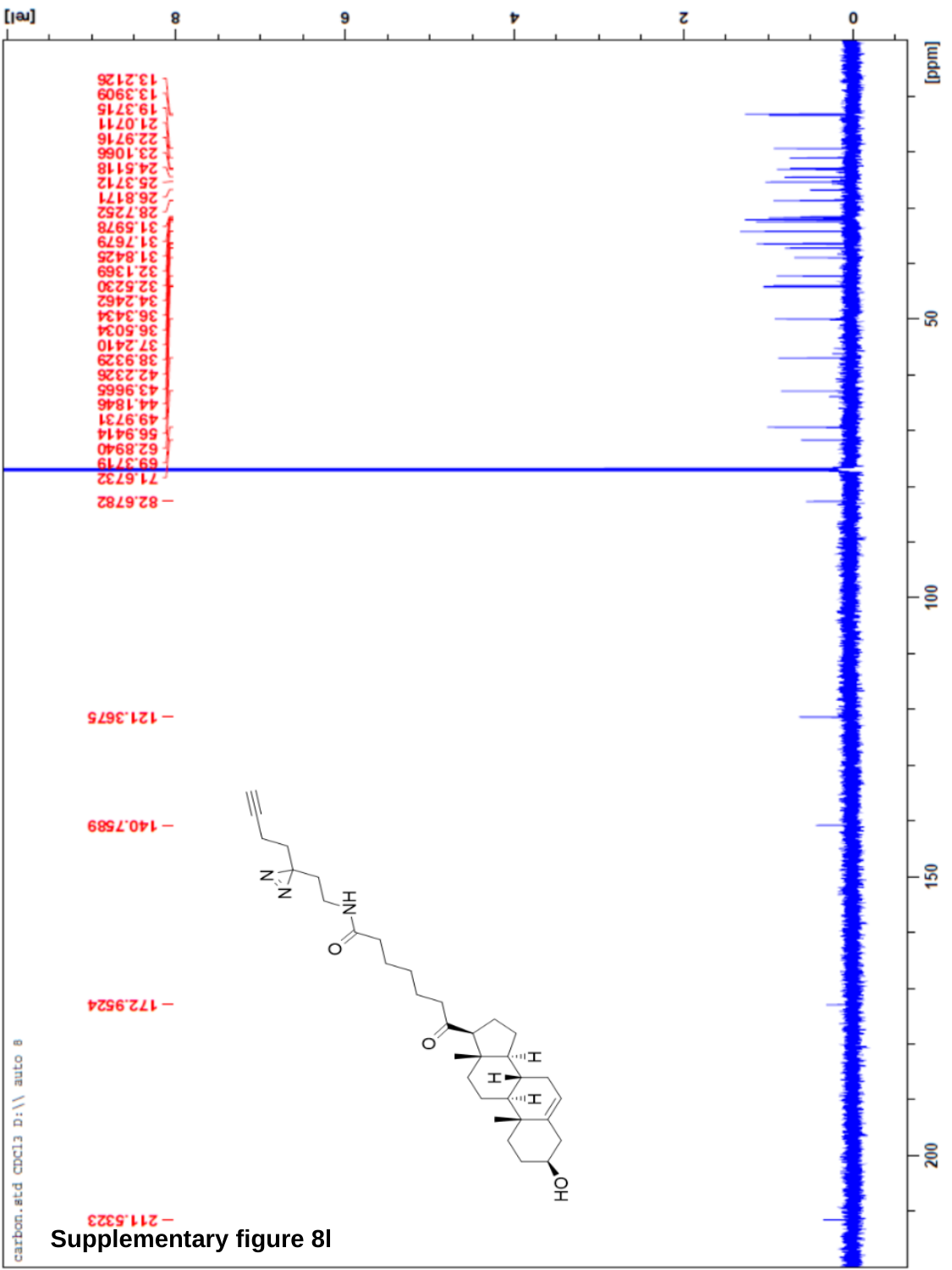

Supplementary figure 8l

### Slide 24
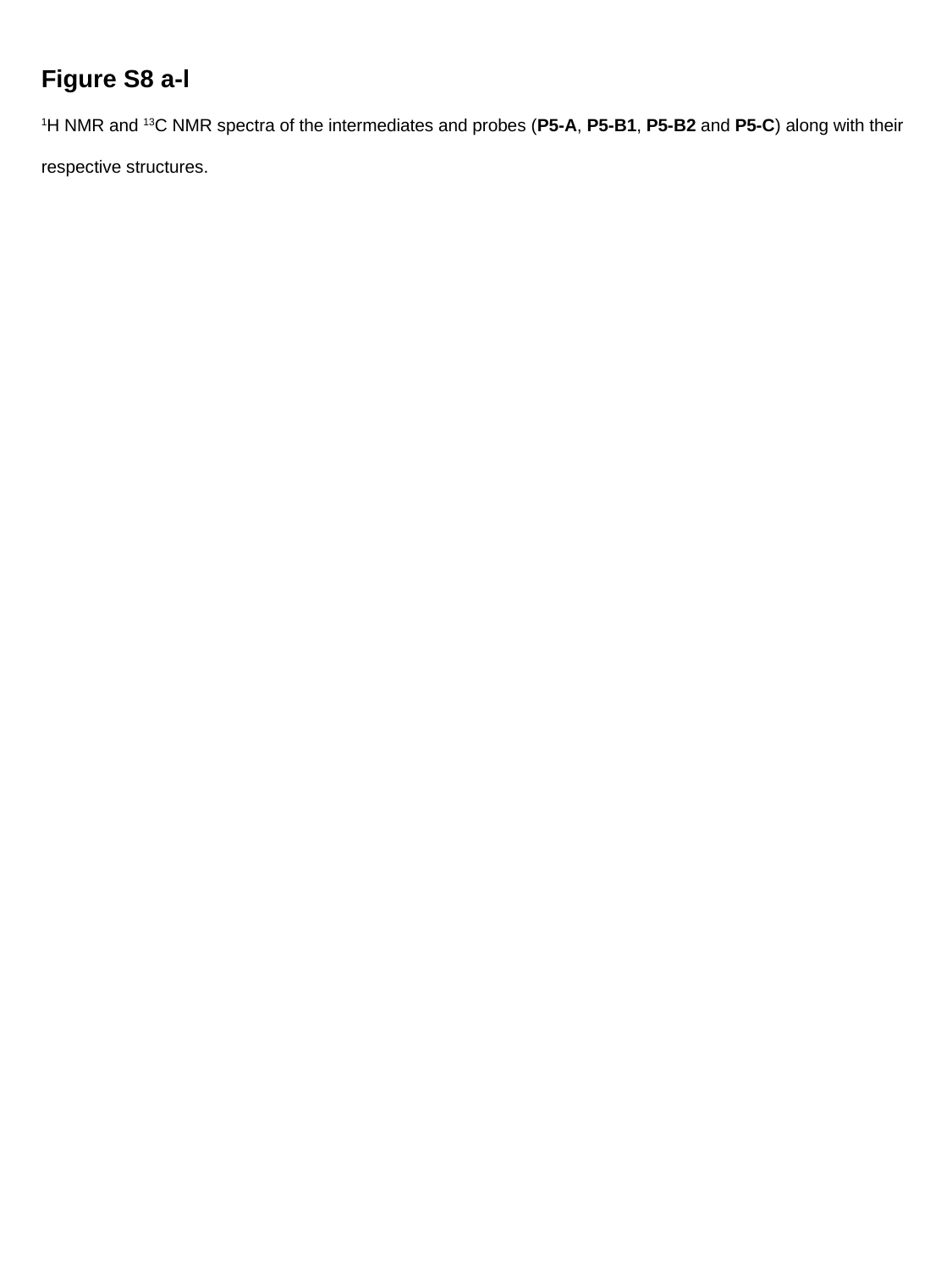

Figure S8 a-l
^1^H NMR and ^13^C NMR spectra of the intermediates and probes (P5-A, P5-B1, P5-B2 and P5-C) along with their respective structures.
